# Decoding heterogeneous aging clocks and disease risk stratification using MetAgeFormer

**DOI:** 10.64898/2026.05.18.725977

**Authors:** Yu Xu, Bohao Zou, Guoxiang Xie, Tianlu Chen, Wei Jia, Lu Zhang, the Alzheimer’s Disease Neuroimaging Initiative (ADNI), the Alzheimer Disease Metabolomics Consortium (ADMC)

## Abstract

Metabolomic aging clocks estimate biological age by modeling metabolite concentrations, thereby capturing aging signals from healthspan and adverse outcomes. However, existing clocks generally assume homogeneous aging trajectories and yield only a single age acceleration metric, limiting their capacity to capture inter-individual metabolic heterogeneity and characterize nuanced individual-level representations. To address these limitations, we proposed MetAgeFormer, a transformer-based metabolomic model pre-trained on nuclear magnetic resonance (NMR) metabolomic profiles from over 430,000 participants in UK Biobank via self-supervised learning. This large-scale pre-training enables MetAgeFormer to learn a metabolomic representation space that captures the complex, nonlinear structure of systemic metabolism as reflected in NMR data. Building on MetAgeFormer, we developed a mortality-informed metabolomic aging clock by fine-tuning an attached survival module, deriving age acceleration that demonstrates significant associations with multiple age-related diseases and factors. We further validated zero-shot transfer in the independent Alzheimer’s Disease Neuroimaging Initiative (ADNI) cohort. More importantly, we utilized embeddings generated by MetAgeFormer to identify 13 distinct metabolic subtypes and consolidated them into four meta-subtypes with markedly divergent susceptibility profiles for major age-related diseases, particularly type 2 diabetes and neurodegenerative disorders. This finding empirically demonstrated substantial metabolic heterogeneity across populations, persisting even at comparable levels of age acceleration. To enhance clinical applicability, we further employed contrastive learning to distill a lightweight model that approximates the learned metabolomic representation space using only 14 routine clinical blood test measurements as inputs. Both hold-out testing within UK Biobank and external validation in the China Health and Retirement Longitudinal Study replicated similar disease onset patterns across the identified subtypes, underscoring the robust generalizability of MetAgeFormer and supporting its translational potential as a scalable framework for metabolomic aging assessment and early disease risk stratification.

## 1 Introduction

Biological aging is a complex, dynamic process characterized by the progressive decline of physiological functions and increased susceptibility to age-related diseases. To quantify this process, various “aging clocks” have been developed using omics data, including epigenomics [1, 2], proteomics [3], and metabolomics [4, 5, 6]. Among these, metabolomics has emerged as a uniquely powerful lens, providing a dynamic readout of an individual’s physiological state [7]. This dynamic nature reflects an inherent metabolic heterogeneity, suggesting that biological aging is not a uniform process but rather a divergent set of pathways [8, 9]. However, the potential of existing metabolomic aging clocks is limited by a mismatch between biological complexity and methodological simplicity. First, most metabolomic clocks established using conventional computational models (e.g., LASSO [4, 5] and ElasticNet [5]) often fail to capture the complex, intrinsic, non-linear relationships among metabolites in large-scale metabolomic data. Consequently, these models typically yield only a single, aggregated aging score, thereby masking metabolic heterogeneity among individuals. These limitations are compounded by the prohibitive cost of metabolic profiling [10, 11], which currently limits its feasibility for routine clinical use.

With their large sample sizes, population-based metabolomics cohorts now enable robust learning of complex non-linear relationships among metabolites. Self-supervised pre-training on these data can extract context-dependent metabolic representations, and the resulting high-dimensional embeddings may also reflect inter-individual variation within a structured representation space. This capacity helps identify subpopulation-level signatures that define divergent aging trajectories. Additionally, knowledge distillation [12] can then transfer the learned representations of a large-scale pre-trained teacher into a compact student architecture. Such a framework can bridge biological fidelity and clinical utility by enabling inference of complex metabolic states from a dramatically reduced blood test biomarker panel.

In this study, we developed MetAgeFormer, a transformer-based metabolomic model pre-trained through self-supervised learning on nuclear magnetic resonance (NMR) profiles from over 430,000 participants from the UK Biobank (UKB) cohort, to decode heterogeneous metabolic aging clocks and disease stratification. The architecture of MetAgeFormer is based on the transformer-encoder architecture [13, 14], which effectively captures the context-dependent relationships among 107 metabolites from the UKB NMR panel [15]. By leveraging MetAgeFormer, we projected participants’ metabolic profiles into high-dimensional embeddings, establishing a metabolomic representation space. We further fine-tuned a survival module using these embeddings and mortality information to derive a robust measure of metabolomic aging acceleration and validated its associations with multiple age-related diseases. We also assessed zero-shot transfer in the independent ADNI cohort. Crucially, we identified 13 data-driven metabolic subtypes based on the metabolomic representation space that capture the inherent heterogeneity of metabolic dysregulation. These subtypes showed markedly divergent trajectories for age-related disease onset, with particularly pronounced differentiation in the onset risk for type 2 diabetes and neurodegenerative disorders. We further summarized these subtypes into four meta-subtypes for disease-incidence analyses. Our analysis further identified differential metabolic profiles that characterize these subtypes, providing candidate metabolite signatures of divergent aging trajectories rather than validated clinical biomarkers. To bridge the gap between MetAgeFormer and clinical utility, we employed a contrastive learning-based distillation framework [16]. This process transferred the deep insights of MetAgeFormer into a lightweight model capable of inferring aging acceleration and metabolic subtypes directly from 14 routine clinical blood measurements. The clinical utility and robustness of this framework were rigorously validated in an independent, ethnically diverse cohort, suggesting MetAgeFormer as a scalable framework for metabolomic aging assessment and early disease risk stratification.

## 2 Results

### 2.1 Metabolomics and Blood test from UK Biobank, ADNI, and CHARLS

UKB NMR metabolomics (UKB Category 220; version: November 2025) includes the quantification of 251 measures (including 107 non-derived and 63 composite metabolites, as well as 81 percentages and ratios) from 488,335 participants. A subset of 19,448 participants completed a second assessment, yielding a total of 507,783 plasma samples. Participants from the UKB assessment centers in England and Wales were used for model training (n=434,366), while participants from Scotland and those with repeated visits formed a hold-out test set (73,417 plasma samples). Moreover, to enable external validation, we incorporated plasma Nightingale NMR data from the Alzheimer’s Disease Neuroimaging Initiative (ADNI), acquired on the identical NMR platform as UKB NMR metabolomics. After harmonizing metabolite definitions with the UKB panel, 107 non-derived metabolites were available across 4,851 plasma samples from 1,524 participants. Among these, 1,489 participants completed at least two NMR assessments (2 to 9 visits per participant). ADNI is a disease-enriched cohort spanning cognitively normal individuals and participants with mild cognitive impairment or Alzheimer’s disease, providing a zero-shot test of model transfer under matched metabolomic profiling. In addition, targeted LC-MS metabolomic profiles measured by our Q300 platform [17] were available for a subset of ADNI samples (n=4,142), with nine chemically matched metabolites mapped to the NMR 107 non-derived metabolites. Additionally, we incorporated data from CHARLS [18], which comprises five follow-up waves conducted in 2011, 2013, 2015, 2018, and 2020. Routine blood test data are available for the 2011 (n=11,823) and 2015 (n=13,257) waves (**Figure 1A**). Among these, 7,539 participants contributed repeated blood samples across both waves. Between 2011 and 2015, 4,284 participants were lost to follow-up, and 5,718 new participants were enrolled in the 2015 wave.

**Figure 1.**
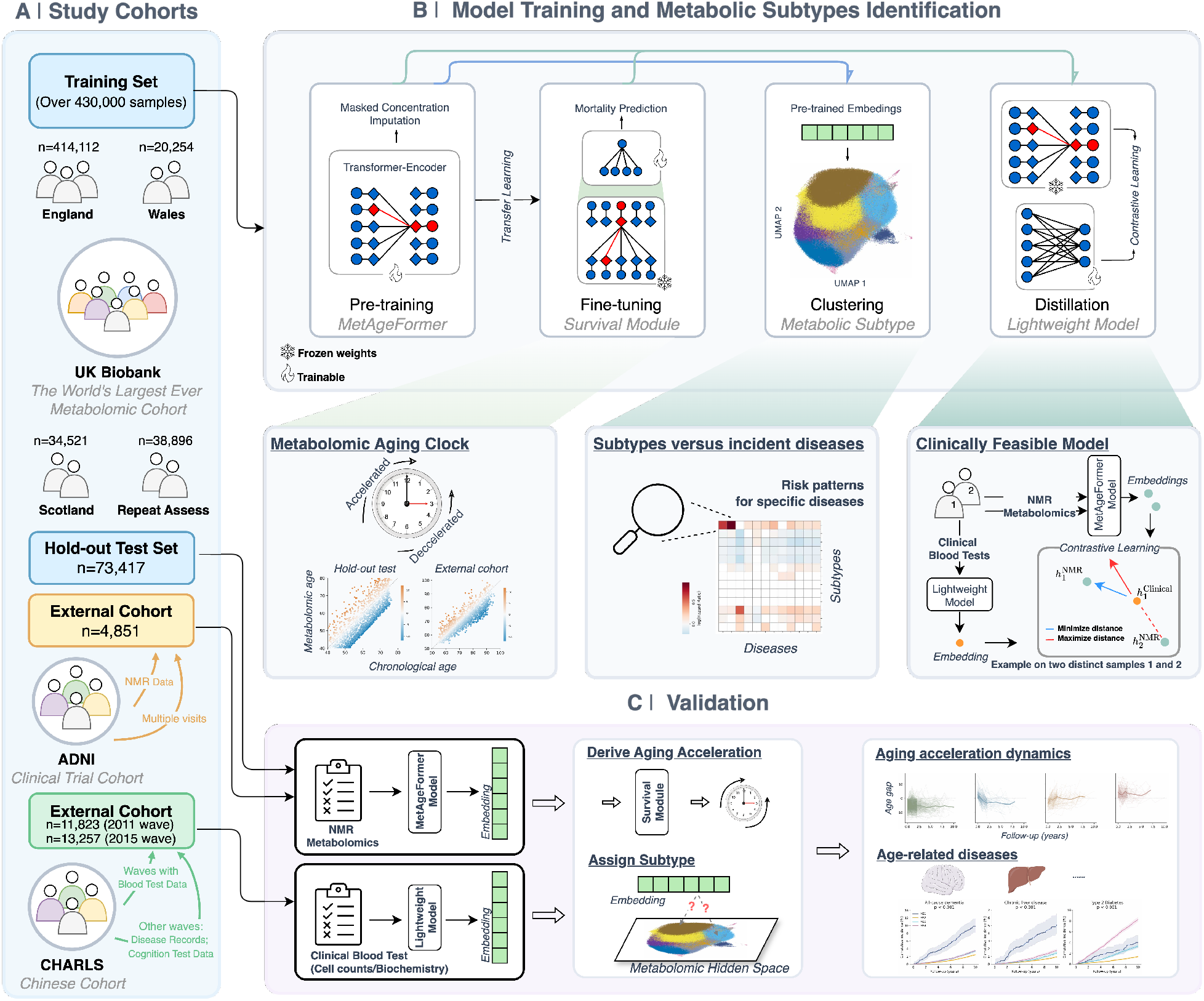
Overview of MetAgeFormer. **(A)** Metabolomic profiles from UK Biobank were divided into a training set and a hold-out test set (for internal validation). Participants with repeated assessments were excluded from the training set to prevent potential data leakage. External validation was performed using datasets derived from the ADNI and CHARLS cohorts. **(B)** The training set was used for four key steps: 1. pre-training MetAgeFormer to learn intrinsic relationships among metabolites; 2. fine-tuning the survival module to predict mortality risk, which was subsequently transformed into biologically interpretable age acceleration (in years); 3. identifying metabolic subtypes from the large-scale dataset of UK Biobank participants from England and Wales, representative of the general population; 4. distilling a lightweight model via contrastive learning between two modalities, NMR metabolomics and routine blood test measurements. **(C)** The hold-out test set and external datasets were used to validate metabolomic aging acceleration and the identified metabolic subtypes.

### 2.2 Overview of MetAgeFormer

The architecture of MetAgeFormer is a stack of six transformer-encoder layers [13] consisting of eight self-attention heads, with an embedding module designed for encoding metabolic profiles (**Methods,** **Figure 2**A, and **Supplementary Figure S1**). Specifically, the embedding module encodes metabolite identity and concentration through two sub-modules. Each metabolite identity is drawn from a fixed vocabulary, initially represented as a one-hot vector and then projected into a symbol embedding via a sub-module constructed using a fully-connected layer. Simultaneously, another sub-module, a stack of fully-connected layers, transforms scalar concentration values into concentration embeddings of matching dimensionality. The symbol and concentration embeddings are averaged for each metabolite to create a unified metabolite representation. Finally, the collection of these metabolite embeddings for an individual serves as the input for the initial transformer-encoder layer.

**Figure 2.**
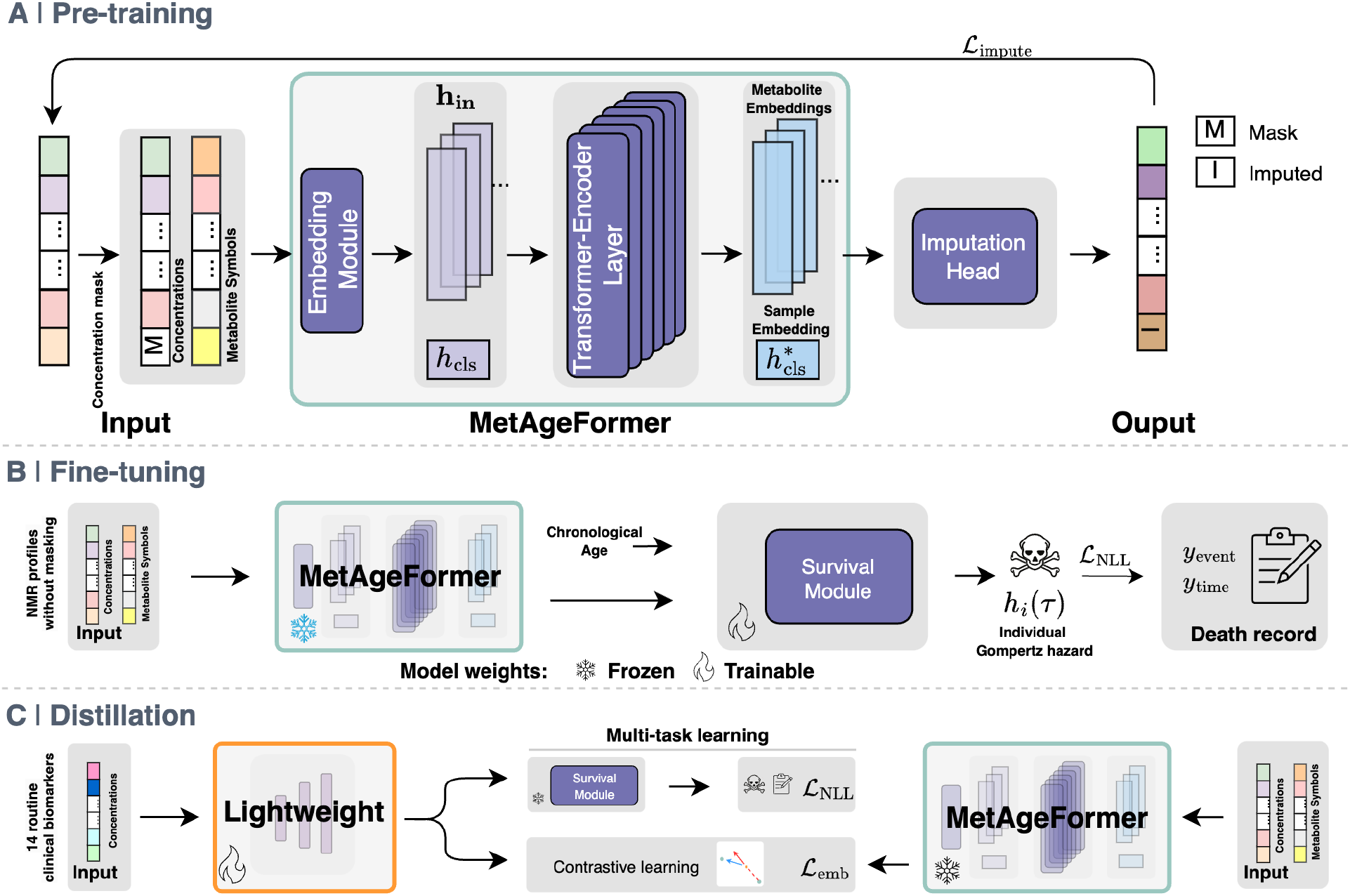
Model architecture and training strategy. **(A)** Masked metabolomic profiles imputation as the pre-training objective, enabling MetAgeFormer to learn intrinsic relationships among metabolites. ℒ_impute_ was computed as mean absolute error. **(B)** The DeepGompertz head of the survival module was fine-tuned to estimate the individual Gompertz hazard (*h*_*i*_(*τ*)). ℒ_NLL_ was computed as the negative log-likelihood (**Methods**). The weights of the pre-trained MetAgeFormer were frozen during fine-tuning of the survival module. **(C)** The lightweight model was distilled from the pre-trained MetAgeFormer via contrastive learning. ℒ_emb_ was computed as InfoNCE. ℒ_NLL_ was incorporated to ensure the lightweight model produced Gompertz hazard estimates comparable to those of the original MetAgeFormer. The weights of both the pre-trained MetAgeFormer and the fine-tuned survival module were frozen during the distillation process.

### 2.3 Pre-training MetAgeFormer on the population-level metabolomic data

MetAgeFormer was pre-trained in a self-supervised manner using masked concentration imputation on a panel of 107 non-derived metabolites (**Methods**, the left panel of **Figure 1B** and **Figure 2**A), comprising 73 measures from 14 lipoprotein subclasses, 9 amino acids, 7 fatty acids, 5 other lipids, 4 glycolysis-related metabolites, 4 ketone bodies, 2 apolipoproteins, 2 fluid balance biomarkers, and 1 inflammation biomarker. We performed an ablation study to assess whether incorporating derived measures (composite metabolites, metabolite ratios, and percentages) into pre-training can improve model capacity. We recomputed the full set of derived measures following the pipeline of Ritchie et al. [15], which yields 219 derived variables (135 percentages, 62 composite metabolites, and 22 ratios) and is therefore broader than the derived measures released directly by UKB. Combining these with the original 107 non-derived metabolites increased the input space to 326 variables (**Supplementary Table S1**). Despite increasing the input space to 326 variables (**Supplementary Table S1**), we observed comparable imputation performance for most metabolites (**Supplementary Figure S2**). This redundancy is consistent with a post hoc inspection of attention patterns (**Methods**), which indicated that the self-attention mechanism can represent covariation among metabolites from raw concentration data alone. Across transformer-encoder layers, attention patterns showed a shift from broadly distributed interactions in early layers toward more specialized interactions in deeper layers (**Supplementary Figure S3**). In exploratory analyses, some individual heads showed patterns concentrated among metabolites from known biochemical groups. For instance, unsaturated fatty acid metabolism (involving *ω*-3 fatty acids, *ω*-6 fatty acids, and docosahexaenoic acid (DHA)) was most prominently represented in head 3 in layer 1 and in heads 2 and 6 in layer 2, branched-chain amino acids (Leucine, Isoleucine, and Valine) in head 2 in layer 2, and ketone bodies (acetoacetate, *β*-hydroxybutyrate, and acetone) in head 8 in layer 3. More comprehensive layer-wise atlas is provided in **Supplementary Data 1**. Together, the ablation results and attention analyses support the practical choice of pretraining on non-derived metabolite concentrations, as many derived measures in the extended panel are computed directly from these non-derived metabolites (e.g., total concentration of branched-chain amino acids (UKB Data-Field 23464) as the sum of Leucine, Isoleucine, and Valine; polyunsaturated fatty acids (Data-Field 23446) as the sum of *ω*-3 and *ω*-6 fatty acids).

### 2.4 MetAgeFormer metabolomic age improves all-cause mortality risk stratification

To assess whether MetAgeFormer captures clinically meaningful biological aging beyond chronological age, we evaluated metabolomic age acceleration (ΔAge = metabolomic age-chronological age) to predict all-cause mortality in the UKB test set. We implemented this evaluation in head-to-head comparisons against existing aging clocks and markers (**Methods**). For each competitor, analyses included only plasma samples with non-missing values for both MetAgeFormer-derived ΔAge and that competitor. Sample sizes therefore differed by comparison (n = 53,843–73,417; death events: 5,114–6,319; details in **Supplementary Table S2**). Within these comparisons, MetAgeFormer achieved concordance index (C-index) of 0.762–0.767 and consistently outperformed metabolomic clocks supervised by chronological age (MileAge [5], C-index approximately 0.715–0.719), a metabolomic aging score (MetaboAge (Deelen) [19]: 0.747), a phenotypic aging clock (PhenoAge [1]: 0.748), Frailty Index [20] (0.741), and clinical biomarkers (Cystatin C: 0.731; HbA1c: 0.726) (**Figure 3**). For ProteomicAge (Hamilton) [3], comparison was limited to the 2,679 participants (**Supplementary Figure S4**) with paired NMR metabolomics and Olink proteomics measurements that were not used in the training of either model (342 death events). MetAgeFormer achieved a C-index of 0.767 vs. 0.739 for ProteomicAge (Hamilton).

**Figure 3.**
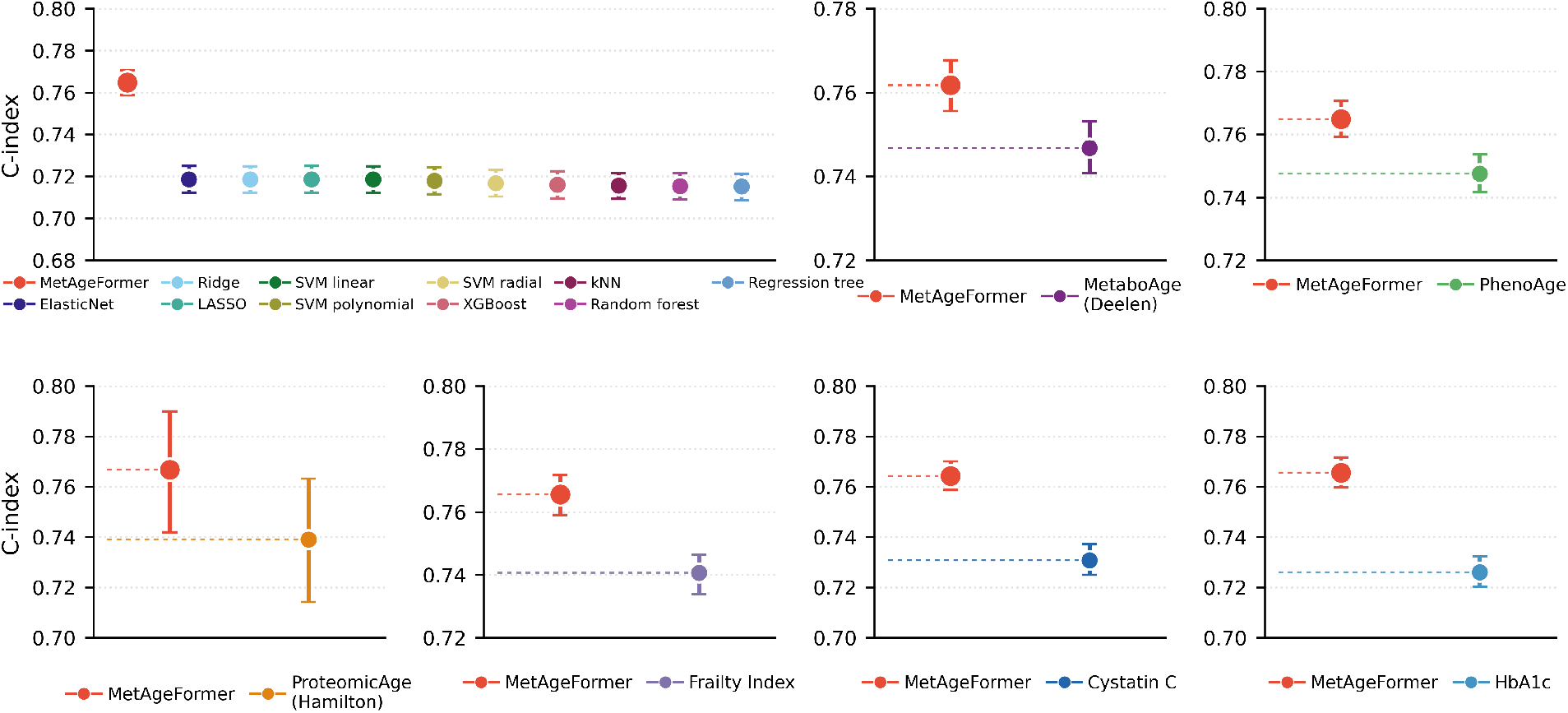
Performance of aging clocks and markers in predicting all-cause mortality. C-index values with 95% bootstrap confidence intervals were obtained from Cox regression models using ΔAge (or aging markers for non-aging clocks) as the predictor, adjusted for chronological age and sex. MetAgeFormer ΔAge demonstrated superior predictive performance, achieving the highest C-index among all evaluated aging clocks and markers.

Additionally, we conducted an ablation experiment to test whether unlocking the full model weights during fine-tuning further improves mortality prediction of ΔAge. Frozen fine-tuning (default: freezing the pre-trained transformer) and full fine-tuning yielded nearly identical C-index (0.765 [0.759, 0.771] vs. 0.764 [0.758, 0.770], p = 0.36), despite modest shifts in the ΔAge distribution (**Supplementary Figure S5**). Moreover, we evaluated 10-year mortality risk using calibration and decision-curve analysis (**Supplementary Figure S6**). MetAgeFormer showed good calibration (calibration slopes 0.94-1.03; expected calibration error ≤ 0.008; Brier 0.039-0.053) and higher net benefit than clinical, metabolomic, and proteomic competitors across threshold probabilities of 1-30%.

### 2.5 Disease associations and cross-cohort robustness of metabolomic age

Building on the mortality analyses above, we next evaluated whether MetAgeFormer-derived ΔAge also predicts incident age-related diseases. Using the identical competitor set as for mortality (MileAge, MetaboAge (Deelen), PhenoAge, ProteomicAge (Hamilton), Frailty Index, Cystatin C, and HbA1c), we assessed Cox models for 12 ICD-10-defined disease endpoints following Hamilton et al. [3], with C-index as the primary metric (**Methods**). MetAgeFormer-derived ΔAge showed broadly higher disease discrimination than alternative metabolomic and aging measures (**Figure 4A**). Because each competitor draws on different UK Biobank fields, every head-to-head comparison was restricted to participants with non-missing values for both measures; C-index values for MetAgeFormer therefore differ slightly between comparisons (**Methods**; sample and event counts in **Supplementary Table S2**). For each disease endpoint, we compared MetAgeFormer to the highest-performing MileAge variant. MetAgeFormer achieved a higher C-index for 11 of 12 diseases, with the largest gains for type 2 diabetes (C-index: 0.807 vs. 0.706 for MileAge (SVM radial)), chronic liver disease (0.720 vs. 0.631 for MileAge (ElasticNet)), and COPD (0.761 vs. 0.688 for MileAge (ElasticNet)). Similar advantages were observed over MetaboAge (Deelen) and PhenoAge for most endpoints.

**Figure 4.**
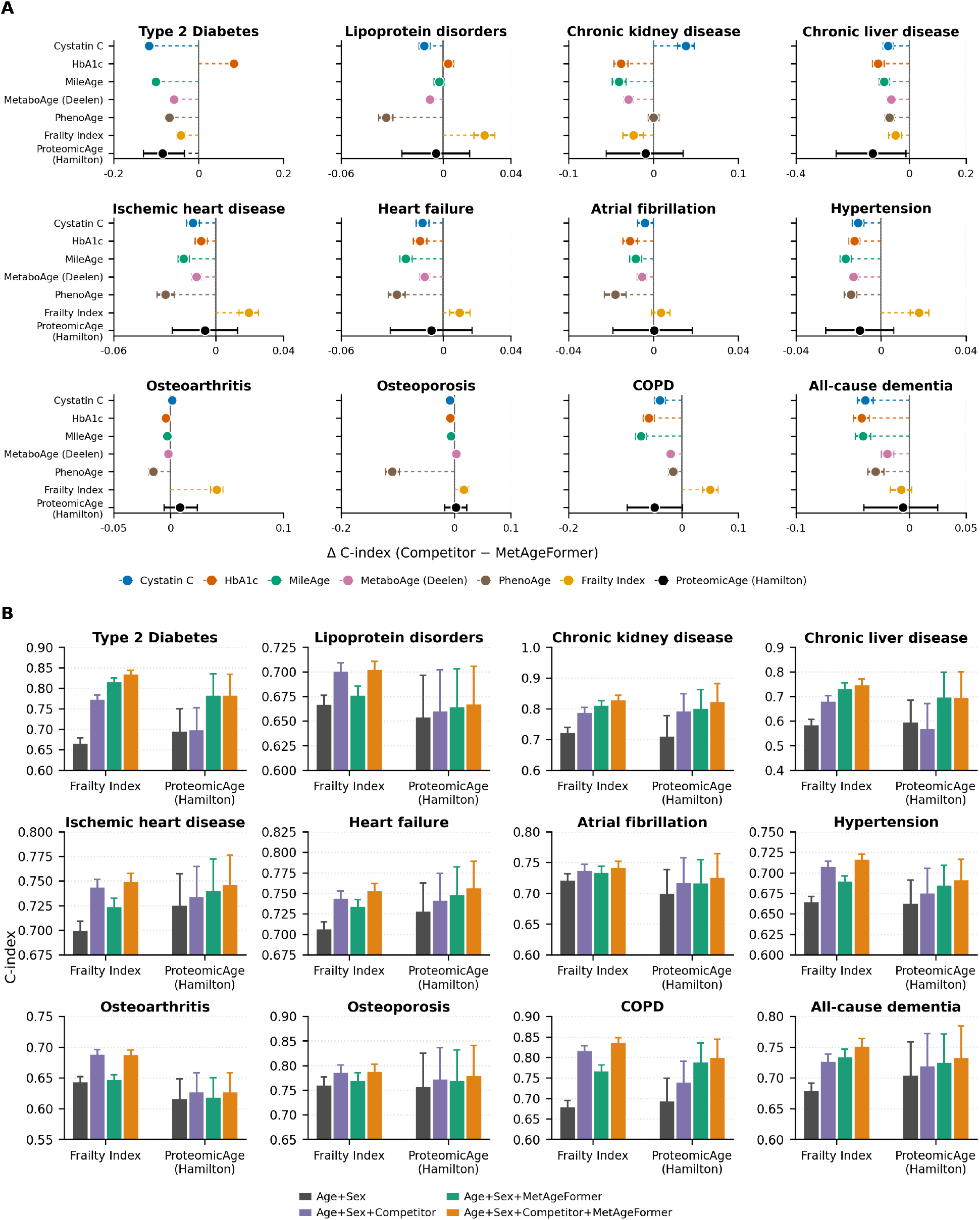
Performance of aging clocks in predicting disease onset. **(A)** Difference in concordance index (ΔC-index; competitor minus MetAgeFormer) across 12 age-related diseases. Negative values indicate a higher C-index for MetAgeFormer. Competitors include Cystatin C, HbA1c, MileAge, MetaboAge, PhenoAge, Frailty Index, and ProteomicAge. For MileAge, the highest-performing variant for each disease is shown. Points show estimates with 95% bootstrap confidence intervals (1,000 iterations). **(B)** Incremental predictive value of MetAgeFormer when added to baseline models that already include Frailty Index or ProteomicAge (Hamilton). Bars show mean C-index with standard deviations from 1,000 bootstrap iterations.

As expected for disease-proximal biomarkers, Cystatin C outperformed MetAgeFormer for chronic kidney disease (0.850 vs. 0.812), and HbA1c outperformed MetAgeFormer for type 2 diabetes (0.893 vs. 0.810). This HbA1c advantage is consistent with substantially higher baseline HbA1c in participants who later developed incident type 2 diabetes than in pure controls (see **Methods**; Cohen’s *d*=2.60; **Supplementary Figure S7A**). Nevertheless, adding MetAgeFormer to HbA1c further improved discrimination for type 2 diabetes (0.904; **Supplementary Figure S7B**). Comparisons with Frailty Index were more mixed: MetAgeFormer was stronger for metabolic outcomes such as type 2 diabetes and chronic liver disease, whereas Frailty Index showed a higher C-index for several musculoskeletal and cardiorespiratory endpoints (e.g., osteoarthritis and COPD). This pattern is expected in part because Frailty Index is constructed from 49 baseline self-reported items that explicitly include diagnosed diseases, disabilities, and related health deficits [20], whereas MetAgeFormer ΔAge is derived solely from NMR metabolomics. MetAgeFormer significantly outperformed ProteomicAge (Hamilton) for type 2 diabetes (0.784 vs. 0.699) and chronic liver disease (0.698 vs. 0.569), with comparable performance for the remaining diseases.

To examine whether these performance differences reflect distinct aging signals, we next correlated MetAgeFormer ΔAge with competitor clocks and markers (**Extended Data Figure 1**). MetAgeFormer ΔAge showed moderate positive correlations with MetaboAge (Pearson r=0.591) and PhenoAge (r=0.482), and weaker correlations with Frailty Index, Cystatin C, and HbA1c (r=0.295 to 0.372), indicating shared components of biological aging. Among the metabolomic aging signals, however, the correlations diverged markedly: MetaboAge (Deelen), which was trained on mortality using NMR metabolites, remained moderately correlated with MetAgeFormer (r=0.591), whereas MileAge (ElasticNet), trained on chronological age, was nearly uncorrelated (r=–0.064). This contrast strongly suggests that the near-zero correlation with MileAge arises primarily from differences in the supervision target (chronological age vs. mortality) rather than from the metabolomic input alone. In contrast, the correlation with ProteomicAge (Hamilton) was also near zero (r=0.044), implying that MetAgeFormer captures aging-related variation not recapitulated by these alternative omics clocks.

Given that ProteomicAge (Hamilton) showed little overlap with MetAgeFormer ΔAge, while Frailty Index has demonstrated complementary disease-specific performance, these distinct patterns led us to test whether MetAgeFormer provides incremental predictive information beyond either of these measures alone. For each disease, we compared four Cox models (**Figure 4B**): (1) demographic baseline: chronological age and sex only; (2) demographic baseline plus Frailty Index or ProteomicAge (Hamilton); (3) demographic baseline plus MetAgeFormer; and (4) demographic baseline plus both MetAgeFormer and the respective competitor. Inclusion of MetAgeFormer consistently improved the C-index over the demographic baseline across diseases. Furthermore, inclusion of MetAgeFormer together with either competitor yielded the highest C-index for nearly all endpoints (11 of 12 diseases) relative to the competitor alone, including COPD for Frailty Index and chronic kidney disease for ProteomicAge (Hamilton). These results indicate that metabolomic age acceleration provides complementary prognostic information to both frailty and proteomic aging signals for incident disease prediction.

We further assessed zero-shot transfer of MetAgeFormer to an external cohort with NMR metabolomics, ADNI [21]. MetAgeFormer-derived metabolomic age positively correlated with chronological age (r=0.776, **Extended Data Figure 2A**). For comparison, we evaluated MileAge (ElasticNet), the only other metabolomic clock trained on the identical NMR metabolite panel that yields age-unit outputs directly comparable to MetAgeFormer. MileAge (ElasticNet) showed a poor correlation with chronological age (r=-0.206; **Extended Data Figure 2B**), consistent with its supervision by chronological age and the marked shift in age distribution between the UKB and ADNI cohorts (**Extended Data Figure 2C**). We observed that higher MetAgeFormer-derived ΔAge was cross-sectionally linked to worse Functional Activities Questionnaire (FAQ) scores, reduced fluorodeoxyglucose standardized uptake value ratio (FDG SUVR), and lower executive function (EF) composite, indicating a possible association with cognitive decline without implying causality (**Extended Data Figure 2D**). In this cross-sectional comparison, mean ΔAge showed a stepwise increase across diagnostic groups from early mild cognitive impairment (EMCI) to late mild cognitive impairment (LMCI) to Alzheimer’s disease (AD) (2.06, 2.25, and 3.43 years, respectively; **Extended Data Figure 2E**). Diagnostic groups differed in mean chronological age (**Extended Data Figure 2F**). CN participants were on average older than the early impairment groups, which likely reflects the ADNI study design that recruits older cognitively normal participants as controls in this disease-enriched cohort. In regression analyses, ΔAge tended to increase with chronological age across all diagnostic groups, with a steeper slope in AD (**Extended Data Figure 2G**), yielding a cross-over pattern across age windows: CN was highest at ages 60–70, groups largely overlapped at 70–80, and CN was lowest at 80–90 (**Extended Data Figure 2H-J**). Given large within-group variance, we emphasize the coarser MCI-to-AD contrast, for which AD showed the highest ΔAge, especially at older chronological ages.

To evaluate robustness to incomplete metabolite coverage across cohorts and assays, we assessed MetAgeFormer under simulated missingness. In the UKB test set, correlation with metabolomic age derived from the full set of 107 non-derived metabolites remained high across missing ratios from 0.2 to 0.8 (Pearson r = 0.989 to 0.909), with only a modest decline in mortality C-index (0.756 to 0.735) (the left panel of **Extended Data Figure 3** and **Supplementary Figure S8A**). In ADNI, the corresponding correlation decreased from 0.965 to 0.806 at a missing ratio of 0.8 (the right panel of **Extended Data Figure 3** and **Supplementary Figure S8B**). Beyond platform-random masking, we further compared metabolomic age derived from ADNI LC-MS using only nine metabolites (**Methods**) overlapping with the NMR panel against predictions from the NMR 107 non-derived metabolites on the same samples (n = 4,142). The experimental results showed that MetAgeFormer retained substantially higher correlation than MileAge (ElasticNet) (r = 0.713 vs. 0.207; **Supplementary Figure S9**). These results indicate that MetAgeFormer retains a substantial fraction of its aging signal under incomplete metabolite coverage, including cross-platform sparse input.

### 2.6 Association of heterogeneous metabolic subtypes with age-related diseases

Although ΔAge provides a concise scalar summary of the biological aging clock, its inherent homogeneity among individuals may neglect the underlying complexity of physiological decline. To better understand aging clock heterogeneity, we applied unsupervised Leiden clustering (**Methods**) to metabolomic embeddings for participants from England and Wales centers (the training set). We searched a grid of the number of neighbors and resolution values and selected the Leiden clustering setting that maximized the minimum cluster size (**Supplementary Figure S10**), which yielded 13 data-driven metabolic subtypes (**Figure 5A** and **Supplementary Figure S11**). We then performed Cox analyses of age-related diseases across these subtypes (**Methods**), identifying subtypes with elevated disease risks (**Figure 5B**). Specifically, subtype 8 showed associations with increased risk of all-cause dementia (Hazard ratio (HR) = 2.578, p < 0.001) and chronic liver disease (HR = 1.946, p < 0.001). This pattern is consistent with previous studies [22, 23, 24, 25, 26] that link hepatic dysfunction to cognitive decline, suggesting that MetAgeFormer may capture biologically coherent comorbidity patterns from metabolic profiles. Meanwhile, subtype 4 was associated with increased risk of the onset of type 2 diabetes (HR = 1.769, p < 0.001).

**Figure 5.**
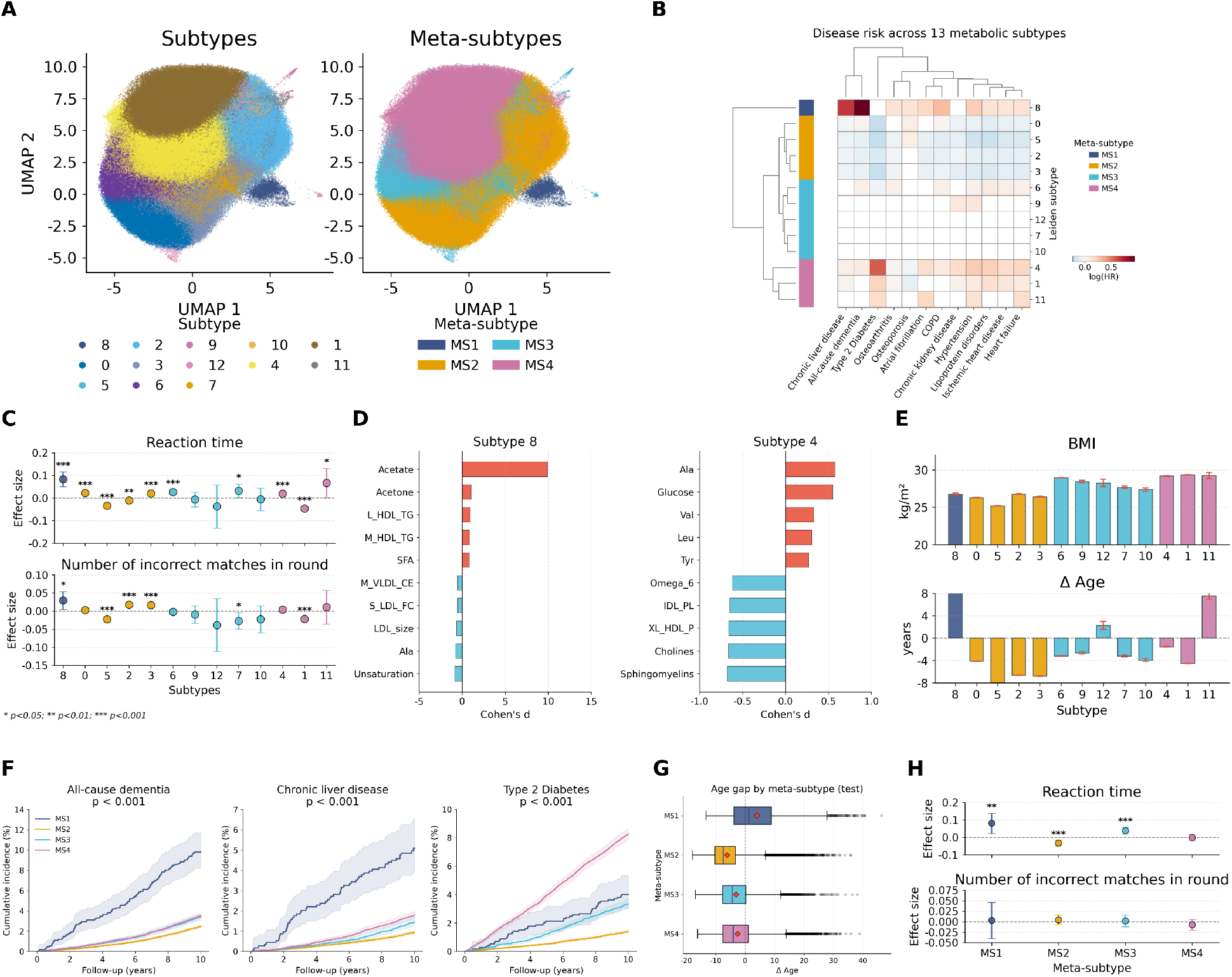
Identification and characterization of metabolic subtypes. **(A)** UMAP visualizations of the metabolomic representation space annotated by the identified subtypes (left) and four meta-subtypes (right). Each point represents a unique participant. **(B)** Heatmap of log_2_(HR) for multiple age-related diseases in 13 subtypes, estimated by Cox models adjusted for chronological age, sex, and BMI. Compared to other subtypes, subtype 8 showed the highest incidence risk for all-cause dementia and chronic liver disease, whereas subtype 4 demonstrated the highest risk for type 2 diabetes. Hierarchical clustering was used to group subtypes with similar disease risk profiles into meta-subtypes. Row colors indicate meta-subtype membership. Blank elements indicate non-significance. **(C)** Associations of subtypes with two cognitive assessments: reaction time and pairs matching (incorrect matches). Effect sizes were estimated by regressing the measures of cognitive assessment on each specific subtype, adjusted for chronological age, sex, and BMI. **(D)** Bar plots of the top 10 metabolites with significantly different profiles (5 positives and 5 negatives) for subtypes 8 and 4. The x-axis represents Cohen’s d effect sizes, and the y-axis represents metabolite abbreviations (e.g., Ala: Alanine, Val: Valine, and Leu: Leucine; more details listed in **Supplementary Table S1**). **(E)** Distribution of BMI and ΔAge across metabolic subtypes. **(F)** Cumulative incidence for three diseases in the hold-out test set, stratified by four meta-subtypes. P-values were calculated using the log-rank test. **(G)** Distribution of ΔAge from the hold-out test set in meta-subtypes. Black lines and red diamonds within boxes represent median and mean values, respectively. **(H)** Validation of cognitive associations of meta-subtypes in the hold-out test set, estimated as in (C).

Besides, we observed distinct risk patterns across the other age-related diseases (**Figure 5B**). Specifically, heart failure, ischemic heart disease, and lipoprotein disorders tended to show higher risk in subtypes 4, 1, 8, and 6, but lower risk in subtypes 0, 5, 2, and 3. Regarding chronic kidney disease and hypertension, subtypes 4 and 1 showed higher risk for both outcomes, with subtype 8 additionally elevated for hypertension, whereas subtypes 5, 2, and 3 showed lower-risk patterns. For atrial fibrillation and COPD, subtypes 8, 4, and 11 were associated with higher risks, with subtype 6 also elevated for COPD, whereas subtypes 5, 2, and 3 showed lower susceptibility. Analysis of musculoskeletal disorders revealed that osteoarthritis risk was higher in subtypes 8, 6, 4, and 1, but lower across subtypes 5, 2, and 3. Meanwhile, osteoporosis risk was notably higher in subtypes 8, 0, and 5, with subtypes 4 and 1 showing lower risk than most other subtypes.

Furthermore, we assessed the associations of metabolic subtypes with two cognitive functions: reaction time and pairs matching, which were used to develop dementia prediction models [27]. Our analysis revealed that subtype 8 showed the strongest associations with both cognitive measures (**Figure 5C**), characterized by significantly slower reaction times (p<0.001) and a greater number of incorrect matches in the card-pair identification task (p<0.05). These patterns are consistent with early cognitive decline characteristic of certain neurodegenerative trajectories.

### 2.7 Characterization of subtypes using metabolic profiles

To characterize metabolite signatures associated with metabolic subtypes, we analyzed differential metabolic profiles across them (**Methods**). We noticed that acetate concentrations were markedly elevated (Cohen’s *d*=9.92) in subtype 8 (the left panel of **Figure 5D**), consistent with recent observational evidence linking circulating short-chain fatty acids (including acetate) to dementia-related phenotypes [28]. Given this extreme elevation, we assessed the sensitivity of clustering and disease-risk associations to acetate perturbation (**Supplementary Methods**). Denoting the disease risk profiles of the original subtypes as the reference, we observed that removing acetate from the input recovered a moderate concordance in the disease risk structure (Spearman’s *ρ*=0.68 vs. the reference, **Supplementary Figure S12A and B**). Winsorizing acetate yielded higher concordance (*ρ*=0.86, **Supplementary Figure S12C**). By contrast, excluding metabolomic samples with extreme acetate values did not recover the original subtype 8, despite a modest overall concordance in the disease risk structure (*ρ*=0.73, **Supplementary Figure S12D**). Focusing on acetate perturbation clusters matched to the original subtype 8, all-cause dementia and chronic liver disease associations remained elevated after acetate removal or winsorization, whereas exclusion of extreme acetate samples decreased the peak disease associations (**Supplementary Figure S12E**). The sizes of these perturbation clusters further illuminated this pattern. Winsorizing acetate recovered a cluster of comparable size to the original subtype 8 (n=3,437 vs. 3,700), whereas removing acetate recovered the same disease-risk pattern in a substantially smaller cluster (857 samples, a 77% reduction; **Supplementary Figure S12F**). We therefore regard the disease-risk structure as reproducible under winsorization and only partially recovered after acetate removal. In contrast, the exclude-outliers setting failed to recapitulate the elevated disease associations of the original subtype 8. This pattern, with winsorization preserving the signal and removal recovering it only partially while exclusion weakened it, suggests that the excluded extreme acetate samples largely constitute subtype 8 itself, rather than representing spurious outliers that artificially drive the disease associations. Additionally, Nightingale NMR acetate and targeted LC-MS acetic acid showed a positive correlation in the ADNI cohort, and samples assigned to subtype 8 exhibited elevated acetate levels on both platforms (**Supplementary Figure S12G**). These analyses support acetate elevation as a reproducible observational feature of subtype 8, but do not establish acetate as a clinical biomarker. In subtype 4, we observed significantly elevated concentrations in glucose (*d*=0.55) and branched-chain amino acids (BCAAs; including valine (*d*=0.33) and leucine (*d*=0.31)) (the right panel of **Figure 5D**), in line with established metabolic dysregulation in diabetes-related conditions [29, 30]. Alanine (*d*=0.57) was also among the most elevated subtype-associated metabolites in subtype 4, despite the lack of direct evidence linking it to type 2 diabetes. Nevertheless, the role of alanine is biologically plausible within the context of BCAA-driven insulin resistance [31]. Specifically, enhanced BCAA catabolism has been shown to increase pyruvate transamination to alanine, thereby contributing to the development of glucose intolerance in obesity. Additionally, circulating alanine concentration was found to be significantly elevated in obese individuals [31], a pattern highly consistent with our observations in subtype 4. The coordinated elevation of BCAAs, alanine, and glucose is therefore consistent with a multi-metabolite signature previously linked to insulin resistance, although causal pathway inference is beyond the scope of this observational analysis. Furthermore, most of the downregulated metabolites in subtype 4 have been previously implicated in the pathogenesis of type 2 diabetes, including sphingomyelins [32], cholines [33], concentration of very large HDL particles (XL_HDL_P) [34], phospholipids (IDL_PL) [35], and omega-6 fatty acids [36].

Beyond the highlighted subtypes 4 and 8, we observed distinct metabolic signatures across the remaining subtypes (**Extended Data Figure 4**). Subtype 1 presented markedly elevated concentrations in VLDL-related metabolites, consistent with reports of enhanced hepatic lipid export [37] and atherogenic risk [38]. Subtype 11 was characterized by elevated glycoprotein acetyls (GlycA) [39], alongside reduced glutamine (Gln) [40, 41], albumin [42], a pattern previously associated with chronic inflammation. Our analysis of subtypes 0, 2, and 12 revealed a complex relationship of ketone bodies (acetone, acetoacetate, and *β*-hydroxybutyrate) and health outcomes. Subtype 0 was characterized by significantly elevated ketone bodies consistent with sustained ketogenesis, coupled with an average BMI of 26.28 kg/m2 (the upper panel of **Figure 5E**). This physiological profile was associated with modestly higher osteoporosis risk (HR=1.074, p<0.001), although the impact of chronic ketosis on bone mineral density remains under debate [43]. Subtype 12 represented an extreme but rare phenotype (n=398, <0.1% of the cohort), marked by exceptionally high concentrations of both ketone bodies and pyruvate. This profound ketogenic state was accompanied by an elevated ΔAge of 2.26 years (the bottom panel of **Figure 5E**) and an average BMI of 28.24 kg/m2, significantly larger than subtype 0 (the upper panel of **Figure 5E**). Such a convergence of extreme metabolic dysregulation and accelerated metabolomic aging is compatible with a rare, extreme metabolic state. In contrast, subtype 2 showed low ketone body levels, a pattern reported in association with more favorable metabolic profiles [44]. Subtype 3 displayed reduced triglycerides, generally favorable for cardiovascular health [45]. Subtype 5 demonstrated a lipoprotein profile often regarded as favorable, with high HDL-related and low VLDL-related metabolites. However, elevated apolipoprotein A1 (ApoA1), glycine, and low BMI levels (average 25.16 kg/m2; lowest among all subtypes) were nevertheless associated with increased osteoporosis risk (HR=1.073, p<0.001) [46, 47, 48]. Subtype 6 showed elevated saturated fatty acids and decreased unsaturation, linked to cardiovascular risk [49]. Subtype 9 demonstrated elevated triglycerides, an independent risk factor consistent with observed hypertension risk [50, 51]. For subtypes 7 and 10, we observed bidirectional deviations in albumin and HDL composition, with a marginal increase in biological age (the bottom panel of **Figure 5E**) but no difference in disease risk (**Figure 5B**), which could reflect compensatory metabolic adjustments that maintain homeostasis [52]. These changes might represent subclinical aging processes that have not yet reached a threshold for disease manifestation, or they may fall within a non-linear physiological range where risk remains stable despite statistically significant metabolic differences.

### 2.8 Characteristics and validation of metabolic meta-subtypes based on age-related diseases and aging clocks

We further categorized these 13 subtypes into four meta-subtypes based on their disease onset risk (log(HR)) in the training set using hierarchical clustering [53] (**Figure 5B**) to increase the sample size of each subtype. This facilitated the validation of our findings in independent cohorts. These four meta-subtypes represented distinct major disease risk signatures: meta-subtype 1, characterized by relatively higher log(HR) for dementia and chronic liver disease; meta-subtype 2, a comparatively low-risk group across most disease outcomes; meta-subtype 3, a baseline risk profile; and meta-subtype 4, associated with elevated relative risk of type 2 diabetes. We evaluated cumulative incidence across meta-subtypes in the hold-out test set to assess associations between meta-subtypes and age-related diseases (**Figure 1C**). Meta-subtype assignment for participants in the hold-out set was performed by utilizing a multi-layer perceptron (MLP) classifier fitted on the training set (**Methods**). As shown in **Figure 5F**, the evaluation results were consistent with the patterns observed in the training set, supporting the reproducibility of these associative risk profiles. To examine whether these findings remained consistent across demographic groups, we performed a series of stratified analyses by chronological age intervals, sex, and socioeconomic status, using the Townsend deprivation index (**Methods**). For all-cause dementia, chronic liver disease, and type 2 diabetes, the stratified groups (age intervals, sex, and different levels of social deprivation) showed trends that were broadly consistent with our initial findings (**Supplementary Figure S13**), further supporting the consistency of these observational associations. Besides the three diseases shown in **Figure 5F**, we also observed the associations between meta-subtypes and the other diseases (**Supplementary Figure S14**) in the test set. Meta-subtype 1 showed relatively higher cumulative incidence of osteoporosis, atrial fibrillation, and COPD, while meta-subtype 2 also showed relatively elevated cumulative incidence for osteoporosis. In contrast, meta-subtypes 3 and 4 exhibited relatively higher cumulative incidence of osteoarthritis than meta-subtypes 1 and 2. Notably, meta-subtype 4 was associated with relatively higher cumulative incidence across most of the remaining cardiometabolic outcomes.

We further derived ΔAge from the hold-out test samples and assessed metabolomic aging levels across meta-subtypes (**Figure 5G**). Meta-subtype 1 showed the highest mean ΔAge, with a mean (median) of 3.99 (1.25) years, consistent with a relatively accelerated metabolomic aging profile in this group. In contrast, meta-subtype 2 showed the lowest mean ΔAge (−6.15 years), consistent with its comparatively lower disease-risk profile. Although meta-subtype 4 had a substantially lower average ΔAge (−2.58 years) compared to meta-subtype 1 (3.99 years) and a value close to that of meta-subtype 3 (−3.12 years), participants in this meta-subtype showed higher cumulative incidence of type 2 diabetes than meta-subtypes 1 and 3. This observation highlights the limitations of relying solely on an aging metric (ΔAge) and emphasizes the need for subtype-specific characterization to better describe associations between metabolomic aging and disease risk. Moreover, we evaluated the associations between meta-subtypes and the two cognitive functions in the hold-out test set. Meta-subtype 1 was associated with significantly longer reaction time, whereas associations with pairs matching were not significant (**Figure 5H**), partially replicating the patterns of subtype 8 observed in the training set (**Figure 5C**).

### 2.9 Longitudinal dynamics of metabolomic aging and metabolic subtypes

We examined the longitudinal patterns of metabolomic aging and metabolic subtypes among a subset of participants who attended repeat visits (n=19,448). By leveraging MetAgeFormer-derived ΔAge, we categorized participants into four distinct groups (**Extended Data Figure 5A**): stable decelerated (consistently negative ΔAge, n=13,562), transition to accelerated (moving from negative to positive ΔAge, n=3,113), transition to decelerated (moving from positive to negative ΔAge, n=973), and stable accelerated (consistently positive ΔAge, n=1,800). With the stable decelerated group serving as a healthy longitudinal reference for Cox analysis, we found that all other groups generally demonstrated a higher risk of disease onset (**Extended Data Figure 5B**). This elevation in risk was most pronounced in the stable accelerated and transition to accelerated groups. Notably, the transition to decelerated group showed hazard ratios close to 1.0 for several diseases, including lipoprotein disorders, ischemic heart disease, osteoarthritis, and osteoporosis, with none reaching statistical significance. Participants whose ΔAge decreased between visits therefore showed a more favorable incident-risk profile than those who remained accelerated. Because these analyses are observational and ΔAge was not experimentally manipulated, they should not be read as evidence that intervening to lower ΔAge would reduce disease risk.

We further investigated the longitudinal stability of metabolic subtypes and found that only 26.6% of participants retained their original subtype over an average 10-year follow-up period (**Extended Data Figure 5C**). When using meta-subtypes as the object of analysis, we observed that 58.0% of individuals retained their original meta-subtype, implying that 31.4% of participants changed subtype while remaining within the same meta-subtype disease-risk class.

To assess whether these ΔAge trajectory groups generalize beyond UKB, we extended the same quadrant-based classification to ADNI participants (n=1,489 individuals with at least two visits). Although ADNI follow-up comprised multiple scheduled visits rather than a single paired repeat assessment, each participant was assigned to one of the four acceleration groups according to ΔAge at their first and last available NMR measurement time points, using the same rules as in UKB. We then plotted all available intermediate ΔAge, expressed relative to the first NMR visit, as individual trajectories within each group (**Extended Data Figure 5**D-G; stable decelerated, n=957; transition to decelerated, n=181; transition to accelerated, n=167; stable accelerated, n=184). Group-mean trajectories were overlaid only at scheduled visits with ≥3 participants per group, where sample size permitted. These group-mean trajectories generally followed the directions implied by the endpoint-based definitions. They remained predominantly below zero in the stable decelerated group (**Extended Data Figure 5**D), declined toward negative values in the transition-to-decelerated group (**Extended Data Figure 5**E), rose toward positive values in the transition-to-accelerated group (**Extended Data Figure 5**F), and remained predominantly above zero in the stable accelerated group (**Extended Data Figure 5**G). These trends offer support for the external reproducibility of MetAgeFormer-derived age-acceleration dynamics in an independent clinical cohort.

### 2.10 A lightweight clinical model to estimate age acceleration and metabolic subtypes

To enable clinical applications of our aging clock and metabolic subtypes, we distilled a lightweight model from the pre-trained MetAgeFormer and validated its effectiveness using the CHARLS cohort (**Methods, Figure 1C**). We derived ΔAge from the CHARLS 2011 and 2015 waves using the distilled lightweight model, evaluating its predictive validity. For each of two waves, we stratified participants into three groups (low-, mid-, and high-aging groups) by ΔAge tertiles, and performed survival analysis using Kaplan-Meier curves. Our analysis demonstrated that the high-aging group showed a markedly lower survival probability compared to the other two groups (**Figure 6A**), which was consistently observed in both the 2011 and 2015 datasets. The mid-aging group showed moderately reduced survival compared to the low-aging group, a pattern that aligns with trends observed in existing metabolomic aging clocks [5, 6]. These results underscore the robustness and generalizability of ΔAge estimated by our proposed framework.

**Figure 6.**
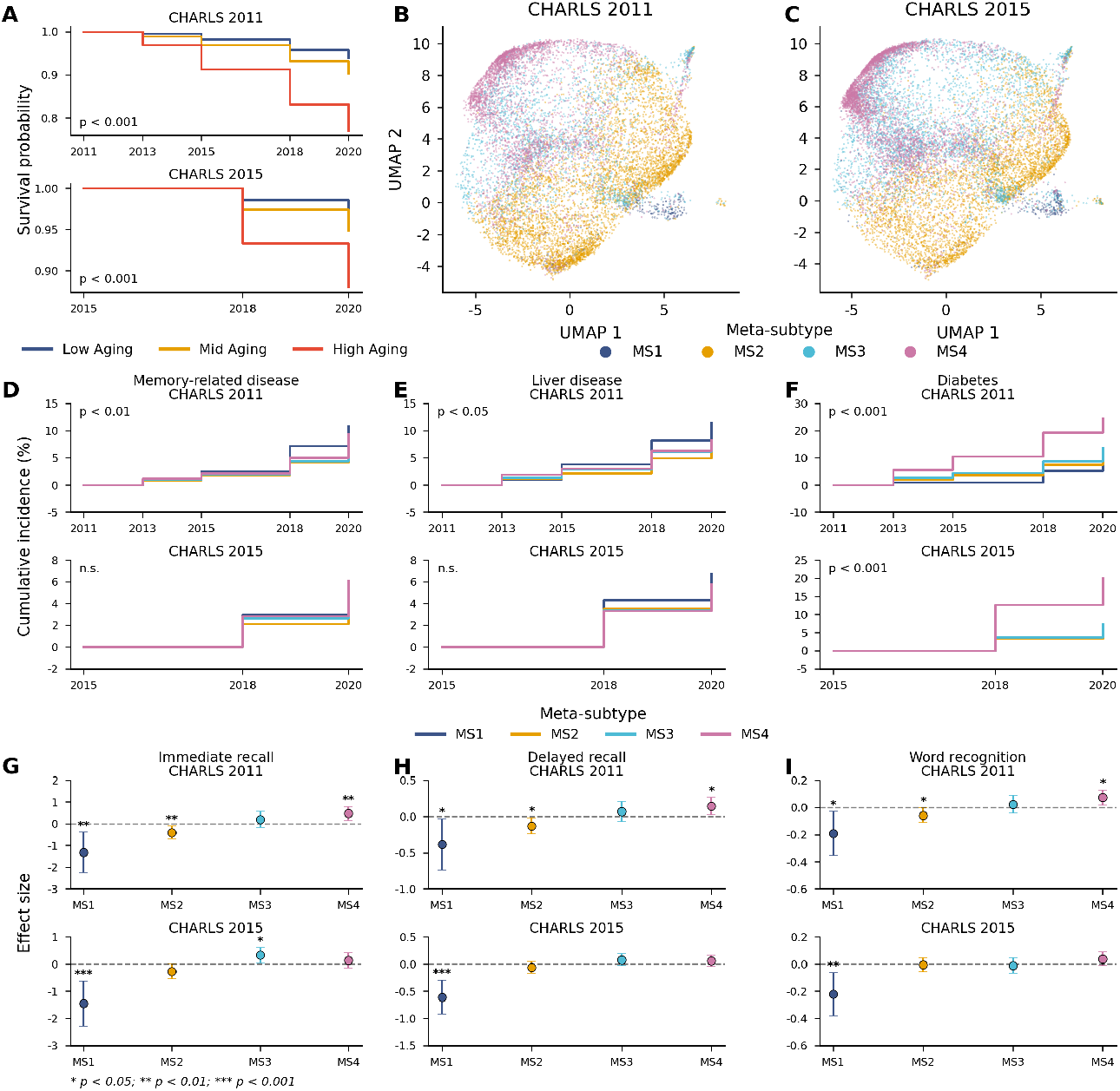
Validation in the CHARLS cohort using the distilled lightweight model. **(A)** Survival curves stratified by ΔAge tertiles in CHARLS 2011 and 2015. P-value was calculated using the log-rank test. **(B-C)** UMAP visualization of the metabolic representation space in CHARLS 2011 (B) and 2015 (C), colored by meta-subtypes. **(D-F)** Cumulative incidence of memory-related disease (D), liver disease (E), and diabetes (F) by meta-subtype in CHARLS 2011 and 2015. P-values were calculated using the log-rank test. ‘n.s.’: not significant. **(G-I)** Associations of meta-subtypes with immediate recall (G), delayed recall (H), and word recognition (I). Effect sizes were estimated by regressing each specific meta-subtype on the performance of memory tests, adjusted for chronological age, sex, and BMI.

We next assigned subtypes for the CHARLS participants by leveraging the MLP subtype classifier fitted on the UKB training set (**Methods**). As illustrated in **Figure 6B–C**, the structure of the metabolomic space observed in the UKB data (**Figure 5A**) is preserved in both the CHARLS datasets, suggesting that the identified subtypes reflect a generalizable pattern of metabolic heterogeneity rather than dataset-specific artifacts. We also conducted an ablation study to evaluate the effectiveness of the distillation process. We observed that the embedding space generated by the lightweight model with distillation showed greater uniformity and aligned more closely with the original metabolomic representation space than the version without distillation (**Supplementary Figure S15**).

Moreover, we analyzed the CHARLS datasets based on meta-subtypes as we did in the UKB test set. We first evaluated the levels of ΔAge across meta-subtypes. In both the 2011 and 2015 waves, the ordering of mean ΔAge across meta-subtypes reproduced that observed in the UKB test set (meta-subtype 1 > 4 > 3 > 2; **Supplementary Figure S16**), although the absolute values were shifted upward. The trend of ΔAge among meta-subtypes in the 2015 dataset mirrors that observed in the UKB test set (the right panel of **Supplementary Figure S16**). Additionally, we noticed that the overall ΔAge in CHARLS was higher than in UKB (**Figure 5G, Supplementary Figure S17**). This ΔAge difference was biologically meaningful, as our ΔAge estimates were comparable across cohorts, which was achieved through multi-task learning during distillation that constrained the lightweight model to predict mortality risk aligned with those of the original MetAgeFormer. The lower ΔAge in UKB compared to CHARLS implies that the UKB participants are generally healthier, consistent with the conclusion from Anna Fry et al. [54]. This is further reflected in the higher disease incidence rates observed in CHARLS (**Figure 5F** and **Figure 6D–F**) over a comparable follow-up duration (10 years in UKB versus 9 years in CHARLS).

We further investigated the incidence patterns of three diseases in CHARLS (memory-related disease, liver disease, and diabetes). In long-term follow-up, the cumulative incidence of high-risk meta-subtypes showed broadly consistent patterns between UKB and CHARLS (**Figure 6D–F**). Specifically, in the CHARLS 2011 dataset, meta-subtype 1 showed the highest observed cumulative incidence for both memory-related and liver diseases. Meta-subtype 4 had a markedly higher observed cumulative incidence of diabetes than the other meta-subtypes, suggesting elevated susceptibility. In the CHARLS 2015 dataset, meta-subtypes 1 and 4 continued to show relatively high observed cumulative incidence for liver disease and diabetes, respectively. However, the cumulative incidence curves for memory-related and liver diseases no longer showed significant differences across meta-subtypes (the bottom panel of **Figure 6D**). This discrepancy may be attributed to differential attrition during the follow-up period from 2011 to 2015 (a total of 4,284 participants were lost). The loss-to-follow-up rates were 38.32% in meta-subtype 1, 35.69% in meta-subtype 2, 36.03% in meta-subtype 3, and 36.95% in meta-subtype 4, respectively. Moreover, participants in meta-subtype 1 presented lower survival probability compared with those in other meta-subtypes (**Supplementary Figure S18**), suggesting a higher mortality rate that may have disproportionately removed incident cases from the at-risk population and thereby biased the observed cumulative incidence for memory-related diseases. To validate the clinical relevance of meta-subtypes in memory-related diseases, we assessed their association with cognitive performance using data from CHARLS wave 2018. Cognitive assessment included CERAD word recall (immediate and delayed) and word recognition tasks. We quantified recall performance as the total number of correctly recalled words and calculated d-prime scores for the recognition task (**Supplementary Methods**), where higher values indicate superior recognition memory. Regression analyses revealed that meta-subtype 1 was associated with the poorest memory capability across all three cognitive domains (**Figure 6G-I**). Notably, blood samples were collected in 2011 and 2015, and memory tests were administered in 2018, enabling prospective evaluation of meta-subtype predictive utility. The effect size for meta-subtype 1 was stronger when using 2015 blood test data (p<0.001 for immediate and delayed recall; p<0.01 for word recognition) compared to 2011 data (p<0.01 for immediate recall; p<0.05 for delayed recall and word recognition), suggesting that proximal measurements yield more robust associations (3-year versus 7-year intervals). These findings emphasized the importance of periodic health monitoring and demonstrated that our lightweight model, which requires only routine blood test data, offers a practical approach to clinical risk stratification of memory decline.

Additionally, we examined longitudinal patterns of metabolomic aging and metabolic subtypes within a subset of the CHARLS cohort (n=7,539) who provided blood samples in both 2011 and 2015 waves. As an observational validation of the UKB repeat-visit analyses, participants were assigned to four aging trajectories based on ΔAge at the 2011 and 2015 visits (**Extended Data Figure 5H**): stable decelerated, transition to accelerated, transition to decelerated, and stable accelerated. In exploratory Cox analysis, diabetes onset appeared more frequent among participants with stable accelerated or transition-to-accelerated aging patterns relative to the stable decelerated reference (**Extended Data Figure 5I**); transition to accelerated aging was also associated with memory-related disease in this cohort, whereas liver disease associations did not reach statistical significance. Investigation of metabolic subtype stability over follow-up was broadly consistent with our initial UKB results (**Extended Data Figure 5J**): 34.2% of participants retained the same subtype and 54.5% remained within the same meta-subtype, suggesting that most longitudinal shifts occurred between profiles with similar risk rather than across distant meta-subtypes.

## 3 Discussion

In this study, we introduced MetAgeFormer, a transformer-based metabolomic model designed to quantify biological aging and stratify disease risk. By pre-training on the extensive UK Biobank NMR metabolomics cohort, MetAgeFormer appears to learn a representation space that captures the complex interdependencies among metabolites. We leveraged this representation to fine-tune a survival module, yielding a measure of metabolomic age acceleration that may serve as an indicator of individual biological aging. Beyond metabolomic aging, unsupervised clustering of these informative representations identified distinct metabolic subtypes. These data-driven clusters may help characterize specific disease onset profiles and their underlying metabolic drivers, potentially connecting molecular signatures with clinical outcomes. In sensitivity analyses with simulated metabolite masking, MetAgeFormer appeared to maintain metabolomic age prediction even at high missing ratios (**Extended Data Figure 3**). Building on this potential tolerance to incomplete metabolomic input, we next explored an exploratory lightweight model distilled from MetAgeFormer via contrastive learning. This model uses only 14 routine blood test measurements yet appears to retain the ability to predict age acceleration and assign metabolic subtypes with reasonable accuracy. Together, our analyses suggest that MetAgeFormer may provide a scalable and interpretable framework for metabolic health assessment, suggesting it may offer a useful approach in precision medicine and population health monitoring.

Biological aging clocks trained on chronological age are prone to bias from age-related confounders. For example, omega-3 fatty acids demonstrated contradictory effect sizes when predicting chronological age versus mortality risk in the UK Biobank [5], likely reflecting the tendency of older participants to consume supplements and maintain healthier diets, especially in health-conscious populations like the UK Biobank cohort [54]. Recent work by Fong et al. [55] has highlighted that age gaps derived from aging clocks trained on chronological age may reflect training artifacts rather than genuine biological aging signals. Their findings demonstrated that ‘risk-equivalent’ aging clocks provide superior predictive performance for clinical outcomes compared to aging clocks trained on chronological age. Building on these insights, we developed a mortality-based aging clock that directly optimizes for survival prediction rather than chronological age, thereby minimizing the confounding effects inherent in age-based training approaches.

Our comparative analysis reveals that MetAgeFormer, ProteomicAge (Hamilton), and Frailty Index capture distinct yet complementary facets of biological aging (**Extended Data Figure 1**). The superior performance of MetAgeFormer in predicting metabolic and systemic conditions, such as type 2 diabetes and chronic liver disease, likely reflects the sensitivity of the metabolome to real-time metabolic flux and acute physiological dysregulation. In contrast, the predictive strength of ProteomicAge (Hamilton) for hypertension and osteoarthritis suggests that proteomic signatures may more effectively capture long-term structural or regulatory changes. Similarly, Frailty Index, a deficit score built from phenotypic and functional measures rather than molecular profiles, outperformed MetAgeFormer for several cardiovascular, musculoskeletal, and respiratory endpoints, including ischemic heart disease, heart failure, and COPD. These findings highlight that aging signals derived from non-omics data types can capture cumulative functional vulnerability that complements molecular clocks. Notably, the significant performance boost observed across nearly all clinical endpoints when integrating MetAgeFormer with ProteomicAge (Hamilton) or Frailty Index underscores a powerful synergy between molecular and phenotypic aging measures (**Figure 4B**). This improvement demonstrated that metabolomic aging signals provide non-redundant information relative to proteomic and clinical deficit indices, where their combination offers a more granular and holistic representation of the aging process than any single measure alone. This complementarity is consistent with hallmarks-of-aging frameworks, in which biological aging arises from multiple interrelated processes [56, 57]. Metabolomic age acceleration should therefore not be interpreted as a direct measure of any one hallmark, but may relate to these frameworks as a systemic metabolic readout that complements other molecular and functional aging measures.

Despite the absence of explicit sex information during pre-training, female and male participants appeared to separate within the metabolomic representation space (**Supplementary Figure S15**). The apparent emergence of sex-specific patterns from NMR profiles alone is consistent with the well-established knowledge of pronounced sex differences in the human metabolome [58, 59, 60]. This observation may support the biological plausibility of the continuous metabolomic representation captured by MetAgeFormer. By capturing nuanced metabolic variations in a compact, information-rich representation, the metabolomic embeddings may have facilitated the identification of subtypes that appear to reflect population-level patterns with potential generalizability. This capacity to identify potentially reproducible subtypes was further supported by the large-scale cohort (half a million participants), which may have provided the statistical power needed to distinguish metabolic signatures from cohort-specific artifacts. This may offer an advantage over smaller datasets that could yield subtypes of limited generalizability.

While traditional case-control experimental designs enable direct causal inference between disease status and biomarkers, our data-driven metabolic subtyping approach offers a complementary advantage: the ability to identify participants at elevated risk of diseases before clinical manifestation. By stratifying participants into metabolic subtypes based on their metabolomic profiles rather than disease status, we can identify individuals residing in high-risk sub-types for specific diseases despite remaining asymptomatic. This prospective identification is particularly powerful for uncovering metabolites critically involved in early disease development pathways, whose perturbations are detectable before clinical diagnosis rather than only among individuals with established disease; whether such perturbations are causally involved in pathogenesis cannot be determined from these observational data. Furthermore, assessing metabolite patterns with distinct profiles within specific metabolic subtypes, collectively as integrated signatures, rather than as isolated biomarkers, provides substantially richer biological insight into the systemic metabolic dysregulation underlying disease susceptibility. This systems-level perspective not only enhances the biological interpretability of our findings but also frames metabolic alterations as interconnected network phenomena, which may help prioritize candidates for future mechanistic and interventional studies. Additionally, the analyzed results for longitudinal participants in both the UKB and CHARLS cohorts showed that subtype membership is not stable over years of follow-up (**Extended Data Figure 5**). Because the metabolomic representation is continuous, part of this label turnover is expected for individuals near subtype boundaries, and measurement variability may also contribute; we therefore regard the four meta-subtypes, at which level all external validation was performed, as the appropriate granularity for longitudinal use, and interpret repeated assessment as a way to update risk classification over time, as periodic reassessments are essential to capture these transitions and provide accurate, up-to-date health evaluations.

For the lightweight model, we selected 14 routine blood measurements shared between the UKB and CHARLS cohorts. To evaluate the relationship between these clinical markers and the underlying metabolome, we calculated Pearson correlation coefficients and mutual information across 107 non-derived metabolites. Our analysis revealed that only a specific subset of these routine measurements showed considerable associations with NMR metabolites (**Supplementary Figure S19**). This suggests that future iterations could enhance the lightweight model by prioritizing clinical variables with higher mutual information, thereby more effectively capturing the metabolic signals required for metabolomic aging and subtype prediction.

Several limitations should be considered when interpreting these findings. First, all analyses are observational. Although incident outcomes were ascertained prospectively, ΔAge and metabolic subtype membership were not experimentally manipulated, so the reported associations do not establish that intervening on metabolomic aging would alter disease risk. Second, pre-training used a targeted NMR panel dominated by lipoprotein measures, which does not cover bile acids, microbial metabolites, neurotransmitter-related metabolites, tryptophan, purine and pyrimidine metabolism, acylcarnitines, or detailed phospholipid and sphingolipid species; the representation learned here should therefore be read as a lipid-weighted view of systemic metabolism rather than a comprehensive one. Third, external validation with directly measured NMR metabolomics was available only in ADNI, which is disease-enriched and substantially older than UK Biobank, whereas the CHARLS replication relies on the distilled 14-blood test measures model rather than on measured metabolomics; the subtype findings therefore await confirmation in an independent population-based cohort with a full metabolomic panel. Fourth, the subtypes are data-driven partitions of a continuous representation space rather than clinically defined categories, and the modest longitudinal retention of subtype labels indicates that individual assignments near subtype boundaries should be treated as provisional. Fifth, differential metabolite profiles and attention patterns are descriptive and neither identify mechanisms nor establish clinical biomarkers. Finally, prospective evaluation, together with a formal assessment of cost-effectiveness and implementation feasibility, would be required before any clinical use; both are beyond the scope of the present study. Additional ablation, robustness and quality-control analyses are reported in **Supplementary Notes 1–7**, including the contribution of self-supervised pre-training, comparison with non-linear Gompertz-based baselines, sensitivity of the subtypes to clustering parameters and embedding normalization, de novo clustering in CHARLS, assay reproducibility, missing data characteristics, and the sensitivity of the BMI adjustment.

Integrating MetAgeFormer-derived metabolomic age with complementary signals from other data types, including ProteomicAge (Hamilton) and Frailty Index, enhances the prediction of disease onset. However, rather than simply aggregating independent scores, a deeper integration of these aging layers is necessary to uncover the complex biological interactions across molecular and phenotypic domains. In future research, we aim to employ advanced fusion techniques, such as Contrastive Language-Image Pre-training (CLIP) [61], to enable joint modeling of multiomics profiles together with readily available clinical aging markers. This approach will facilitate a more nuanced understanding of the cross-talk between the metabolome, proteome, and phenotypic measures of functional decline, ultimately providing deeper insights into the systemic drivers of biological aging.

## 4 Methods

### 4.1 Study Cohorts

#### UK Biobank

For NMR metabolomics, we performed the quality control pipeline (Ritchie et al. [15]) to correct technical variation in non-derived metabolite concentrations and compute composite metabolites, ratios, and percentages. Following the previous study (Julkunen, Heli, et al. [62]), we identified disease cases by utilizing 3-character ICD-10 codes (Chapters A-N) to screen both primary and secondary diagnoses from HES (UKB Category 2002). ICD-9 codes in HES were converted to corresponding ICD-10 codes using mappings from the Center for Disease Control (https://ftp.cdc.gov/pub/Health_Statistics/NCHS/Publications/ICD10CM/2018/2018_I9gem.txt). Prevalent cases were defined by diagnoses occurring up to 25 years before the UKB assessment visit (UKB Data-Field 53), while incident cases were those diagnosed within 10 years after the visit. For incident analyses, prevalent cases were excluded to ensure a focus solely on new diagnoses for each disease. Furthermore, any disease with fewer than 50 cases was excluded from all analyses in this study. Adopting the definitions from Hamilton et al. [3], we defined the following disease groups: Type 2 diabetes (E11), Ischemic heart disease (I20–I25), Chronic liver disease (K70, K73–K76), Chronic kidney disease (N18), All-cause dementia (A81, F00–F03, F05, F10, G30, G31, I67), Osteoporosis (M80, M81), Osteoarthritis (M15–M19), COPD (J44), Lipoprotein disorders (E78), Hypertension (I10–I13, I15), Heart failure (I11, I25, I42, I50), and Atrial fibrillation (I48). Pure controls were defined as participants with no prevalent record of any of the above diseases at baseline, and remained free of incident events for all of the above diseases during follow-up. The follow-up for hospitalizations was completed by October 31, 2022 (in England), August 31, 2022 (in Scotland), and May 31, 2022 (in Wales). The incident events for each of the above disease groups in UK Biobank are summarized in **Supplementary Figure S20**, with prevalent cases excluded. Mortality data were retrieved from death registries (UKB Category 100093), with follow-up completed by November 30, 2022. Chronological ages of the participants were calculated using their date of birth (UKB Data-Fields 34 and 52) and the date they attended the UKB assessment centers, which coincided with the date of blood sample collection. Participant baseline characteristics and mortality outcomes in UK Biobank are summarized in **Supplementary Table S3**. All data were downloaded from the UKB Research Analysis Platform (UKB-RAP) using the DNAnexus Platform SDK ‘dx-toolkit’.

Participants recruited from the UKB assessment centers in England (n=414,112) and Wales (n=20,254) were used for model training (denoted as *D*_train_, n=434,366), excluding those with repeat visits. Participants from UKB centers in Scotland (n=34,521), along with all individuals who had repeat visits (n=19,448*2), formed the hold-out test set (*D*_test_, n=73,417). This rigorous split ensures that no individual appears in both training and testing sets and that participants from the Scotland UKB centers represent distinct geographic regions, consistent with established protocols [4]. We further split 1% from the training set (*D*_train_) as a validation set (*D*_val_, n=4,320) to enable early stopping during model training. Specifically, we applied multi-label stratified sampling [63] to ensure a balanced distribution of disease cases among the remaining training 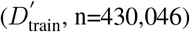 and validation (*D*_val_) sets.

#### ADNI

We constructed an external NMR dataset from ADNI, with metabolomics measured on the same platform as the UKB cohort. The same 107 non-derived metabolites were Z-score normalized using the mean and standard deviation estimated from the UKB training set 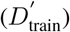 We excluded samples with missing chronological age, yielding 4,851 metabolomic samples from 1,524 participants (1,489 with repeated measurements). Chronological age at the time of each sample was calculated as the baseline age at the earliest assessment plus the number of years between that assessment and the sample collection. Five diagnostic groups are defined in ADNI: cognitively normal (CN; n=1,308), subjective memory complaint (SMC; n=173), early mild cognitive impairment (EMCI; n=811), late mild cognitive impairment (LMCI; n=1,251), and Alzheimer’s disease (AD; n=1,308). For the main analyses, SMC was pooled with CN.

To assess the performance of MetAgeFormer aging clock, we also incorporated several additional metrics, namely the Functional Activities Questionnaire (FAQ), the fluorodeoxyglucose standardized uptake value ratio (FDG SUVR), and an executive function composite (EF). Concentration distribution comparisons of NMR metabolites between the UKB and ADNI cohorts are shown in **Supplementary Data 2**. In supplementary analyses, targeted LC-MS Q300 metabolomic profiles [17] were utilized to assess cross-platform concordance of metabolomic aging, using the ADNI samples with both NMR and LC-MS profiles (n=4,142). Nine chemically matched metabolites were mapped to the NMR 107 non-derived metabolites: alanine, glycine, histidine, leucine, valine, tyrosine, citrate, acetate, and *β*-hydroxybutyrate. Concentration distribution comparisons of these metabolites between NMR and LC-MS platforms in the ADNI cohort are shown in **Supplementary Data 2**.

#### CHARLS

We established two datasets using baseline data from the CHARLS 2011 and 2015 waves, which included clinical blood data measurements. To ensure consistency with UKB data, we standardized the units for blood biomarkers in the CHARLS 2011 and 2015 datasets. HbA1c values, originally in percent (%), were converted to mmol/mol using the standard IFCC (International Federation of Clinical Chemistry and Laboratory Medicine) equation: HbA1c (mmol/mol) = (HbA1c (%) −2.15) / 0.0915. Other biomarkers measured in mg/dL were converted to mmol/L by dividing by their respective molecular weight-based conversion factors: glucose (18.0), triglycerides (88.6), creatinine (11.3), HDL and total cholesterol (38.7), and urate (16.8). The distribution comparison of clinical blood biomarker concentrations across the UKB and CHARLS cohorts is listed in **Supplementary Data 2**.

To prepare the data for survival analysis, we linked mortality information from the 2013, 2015, 2018, and 2020 follow-up waves to the CHARLS 2011 baseline dataset. The 2011 wave served as time zero. Participants who died between waves were assigned the event time of the subsequent wave (i.e., 2, 4, 7, or 9 years). Participants who survived were right-censored at 9 years (2020 wave). This entire procedure was replicated using the CHARLS 2015 dataset as the baseline. Participant baseline characteristics and mortality outcomes in CHARLS are summarized in **Supplementary Table S3**.

For the CHARLS 2011 dataset, prevalent cases were defined as participants with a self-reported physician diagnosis at baseline. To ensure reliability, these self-reports were verified using a disease confirmation questionnaire from the 2013 wave. If a participant’s 2013 response contradicted their 2011 report, their baseline status was deemed unreliable and coded as missing. Incident cases were defined as initially disease-free participants who developed a specific disease during follow-up (2013, 2015, 2018, 2020), with time-to-event assigned to the first wave of diagnosis (2, 4, 7, or 9 years). Follow-up data reliability was managed similarly: for the 2013 and 2015 waves, we used the confirmation questionnaire to verify the status of the prior wave, excluding any records that became unreliable. Since the 2018 and 2020 waves lacked this questionnaire, we applied a cumulative data check, removing any participant with unreliable or missing data from any previous wave. For the 2020 wave, we performed an additional historical consistency check against the lifetime disease history questionnaire, excluding records with discrepancies. Disease-free participants with complete, consistent data were right-censored at 9 years. For the CHARLS 2015 dataset, we followed the same principles for two follow-up waves (2018 and 2020), with time-to-event at approximately 3 or 5 years. As the 2018 wave lacked the confirmation questionnaire, incidence was determined by a direct transition from disease-free status in 2015. For the 2020 wave, we applied the same rigorous cumulative data and historical consistency checks as used for the CHARLS 2011 dataset. The resulting incident events are summarized in **Supplementary Figure S20**, with prevalent cases excluded.

### 4.2 Input embeddings and model architecture

Targeted metabolomic data are presented as a concentration matrix **x** ∈ ℝ^*n*×*k*^ (sample-by-metabolite), where each element is the absolute concentration value for a specific metabolite in a given sample. *n* and *k* represent the number of samples and metabolites, respectively. Inspired by natural language processing (NLP) methodologies, we assigned a unique identifier to each metabolite *m*_*j*_ (*j* ≤ *k*) and created an identifier vocabulary. This allowed us to conceptualize individual metabolites (with their concentration values) as “words” and whole samples as “sentences.” Each metabolite is represented as an integer **id**(*m*_*j*_) indicating its position in the vocabulary. Therefore, the input metabolites for each sample are represented by a vector **m** = [**id**(*m*_1_), **id**(*m*_2_), …, **id**(*m*_*k*_)]. The units of absolute concentrations in the NMR metabolomics are unified as millimoles per litre (mmol/L) and grams per litre (g/L), leading to various value scales among metabolites. To alleviate the effect of value scales, we adopted Z-score normalization to rescale concentration values for each metabolite *m*_*j*_. The normalized input concentration for sample *i* is 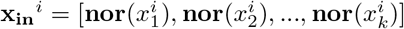. The embedding module (**Supplementary Figure S1A**) comprises two submodules: **emb**_**m**_ for metabolite identifiers (**m**) and **emb**_**x**_ for concentrations (**x**_**in**_). **emb**_**m**_ employs PyTorch’s embedding layer (https://pytorch.org/docs/stable/generated/torch.nn.Embedding.html) to map metabolite identifiers to dense vector representations. **emb**_**x**_ projects normalized concentrations into vector embeddings by fully connected neural networks, which is constructed as follows:

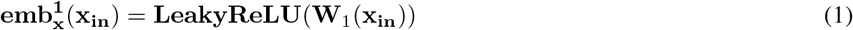

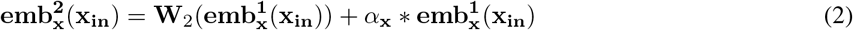

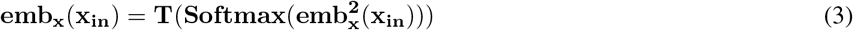

where **W**_1_ ∈ ℝ^1×*b*^, **W**_2_ ∈ ℝ^*b*×*b*^, and *α*_**x**_ (scaling factor) are trainable parameters. **LeakyReLU** is the activation function implemented by the function from PyTorch (https://pytorch.org/docs/stable/generated/torch.nn.LeakyReLU.html) for capturing non-linear relationships. The output of **emb**^**2**^ is normalized by **Softmax** to produce attention weights, which are used for computing a weighted average of vectors from a look-up table **T** ∈ ℝ^*b*×*d*^. *b* (100 by default) is the pre-defined number of vectors in the look-up table, and *d* (512 by default) is the hidden embedding dimension. This approach for encoding scalar values has been demonstrated by Hao et al. [64] in their work on encoding gene expression values. Subsequently, the outputs of these two sub-modules **emb**_**m**_ and **emb**_**x**_ are merged as model input embeddings (**Equation 4**).

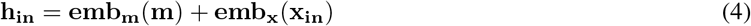

Furthermore, we incorporated three specific trainable embeddings *h*_cls_, *h*_mask_, and *h*_pad_ ∈ ℝ^*d*^ to perform sample representation learning, masked concentration imputation, and padding for mini-batch operation, respectively. During model pre-training, if a sample contains metabolites with zero or missing concentration values, we exclude those specific metabolites in constructing input embeddings from that sample.

The backbone of MetAgeFormer is a stack of six transformer-encoder layers [13], each with 8 (*d/*64) self-attention heads, which utilize self-attention mechanism to model metabolite relationships from input embeddings (**Supplementary Figure S1B**). The dimension of feed-forward layers within each transformer-encoder layer is 2048 (*d* × 4) [65]. The number of model parameters is over 19 millions. The dropout rate and the activation function within transformer-encoder layers are 0.1 and ReLU. The model forward pass can be formulated as

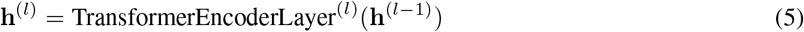

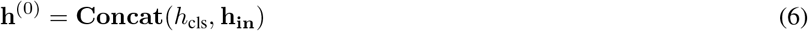

**h**^(0)^ represents the input to the first transformer-encoder layer, which is generated by concatenating *h*_cls_ and **h**_**in**_. **h**^(*l*)^ represents the corresponding output embeddings of the *l*-th Transformer-encoder layer.

### 4.3 MetAgeFormer pre-training

For masked concentration imputation, a subset of metabolite concentration embeddings (**emb**_**x**_(**nor**(*x*_*j*_)), *j* ∈ [1, …, *k*]) is randomly replaced with a specific embedding *h*_mask_ to enable self-supervised learning (**Figure 2A**). Unlike BERT’s fixed 15% masking rate [14], we adopt a dynamic masking rate *γ*(*r*) based on the cosine function [66]: *γ*(*r*) = *cos*(*r*∗*π*∗ 0.5), where *r* ~ Uniform[0, 1) and *γ*(*r*) ∈ (0, 1]. To ensure sufficient information retention, *γ*(*r*) is scaled by 0.8, limiting the maximum masking ratio to 80%. The attention mechanism is adapted to a mixed-directional approach (**Supplementary Figure S21**): masked positions are excluded as attention keys for all other tokens, so a masked concentration never contributes to the representation of any other metabolite. Each masked position may attend only to the unmasked concentrations and to itself, and not to other masked positions, whereas unmasked positions attend bidirectionally among themselves. A fully connected imputation head (**Supplementary Figure S1C**) predicts masked concentrations:

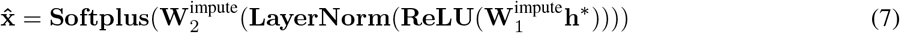

where **h**^∗^ denotes the final transformer-encoder output, and 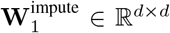 and 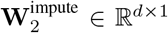 are trainable parameters. Predicted values 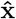 are compared to **x**_**in**_ using mean absolute error (MAE) loss ℒ_impute_, excluding predictions from padding embeddings (*h*_pad_).

We pre-trained the model on an Nvidia RTX-4080 GPU card using a mini-batch size of 128. An early-stopping strategy was employed with a maximum of 300 epochs and a patience of 10 epochs, monitoring the pre-training loss (ℒ_impute_) in the validation set to prevent overfitting. For learning rate scheduling, we adopted a warm-up phase where the learning rate linearly increased from 1e-5 to a peak of 1e-4 over the first 30 epochs, followed by a cosine decay schedule reducing the learning rate to 0 for the remaining pre-training. AdamW optimizer [67] was employed for model optimization.

### 4.4 Analysis of attention patterns

To characterize the dependency structure learned by MetAgeFormer, we analyzed attention scores derived from the pre-trained model using the hold-out test set. These scores represent the relative contribution of each metabolite, with higher values indicating greater weight within the self-attention mechanism. To summarize layer- and head-specific attention structure across the UKB NMR metabolomics panel, we aggregated attention scores across all test samples, resulting in 48 self-attention matrices (R^107×107^) corresponding to the 8 attention heads across 6 transformer-encoder layers. For each attention head, we summarized metabolite-level attention by computing the mean attention each metabolite received from all others, including itself. These aggregates were used for descriptive visualization only and were not interpreted as evidence of causal biological mechanisms.

### 4.5 Survival module fine-tuning

The survival module (**Supplementary Figure S1D**) comprises a population-level Gompertz baseline and a Deep-Gompertz head attached to the pre-trained MetAgeFormer model, enabling the full metabolomic profile to inform the mortality hazard while retaining a shared chronological age effect (**Figure 2B**). The DeepGompertz head is attached downstream of the transformer-encoder layers and takes metabolomic embeddings as inputs. Before fine-tuning, we first fit the population-level Gompertz baseline on the UKB training set to characterize the reference age–mortality relationship:

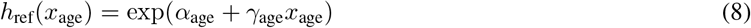

where *α*_age_ is the log-hazard intercept, *γ*_age_ is the increase in log-hazard per year of chronological age, and *h*_ref_(*x*_age_) represents the reference mortality hazard at chronological age *x*_age_.

Given a metabolomic embedding 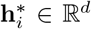 and a scalar chronological age *x*_age,*i*_ for individual *i*, the DeepGompertz head defines the instantaneous mortality hazard at follow-up time *τ* as

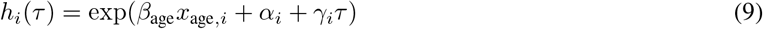

where *β*_age_ is a population-shared coefficient, *α*_*i*_ is the individual log baseline hazard level, and *γ*_*i*_ is the individual Gompertz aging rate. The parameters *α*_*i*_ and *γ*_*i*_ are predicted from 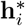 by two parallel feed-forward heads:

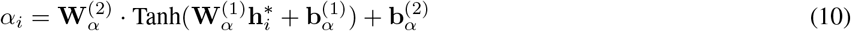

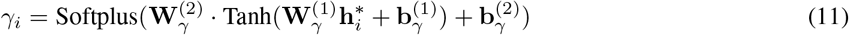

Where 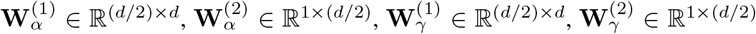, and the corresponding bias terms are learnable parameters. During fine-tuning, *τ* is set to each individual’s observed follow-up time in the survival likelihood.

Under right censoring, the log-likelihood of individual *i* is determined by the hazard at the end of follow-up and by the cumulative hazard accrued over the observed interval:

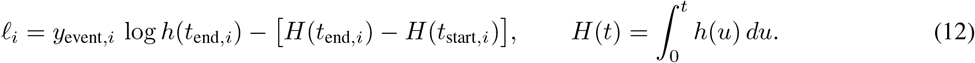

The training loss is the negative mean log-likelihood:

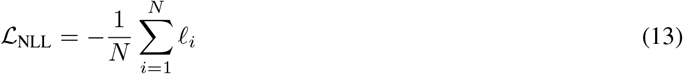

For the Gompertz baseline, the time axis is chronological age. Conditioning on survival to enrollment implies left truncation, so *t*_start,*i*_ = *x*_age,*i*_ and *t*_end,*i*_ = *x*_age,*i*_ + *y*_time,*i*_. Substituting *h*_ref_ (**Equation 8**) and integrating yields:

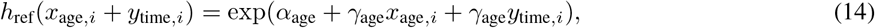

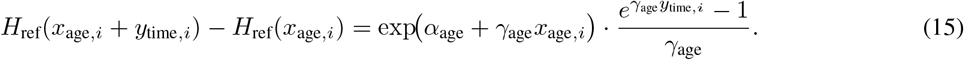

By summarizing **Equations 12, 13, 14, and 15**, the parameters of the Gompertz baseline are therefore optimized by minimizing:

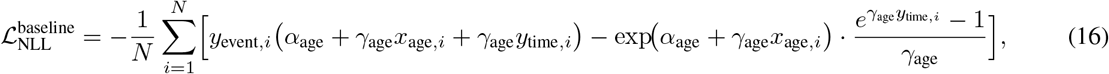

where *x*_age,*i*_ is chronological age at enrollment, *y*_time,*i*_ is follow-up time, and *y*_event,*i*_ is the death indicator.

For DeepGompertz, the time origin is enrollment, so *t*_start,*i*_ = 0 and *t*_end,*i*_ = *y*_time,*i*_. Substituting *h*_*i*_(*τ*) (**Equation 9**) and integrating over [0, *y*_time,*i*_] yields

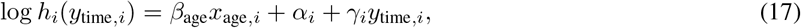

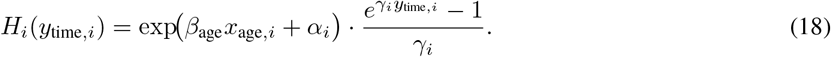

By summarizing **Equations 12, 13, 17, and 18**, the weights of the DeepGompertz head are therefore optimized by minimizing:

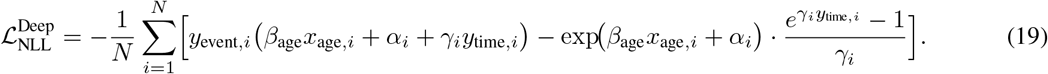

This objective constrains the network to prioritize mortality-associated metabolomic variation through individualized hazard estimation.

The population-level Gompertz baseline was first fitted on 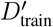 by minimizing 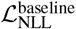 with a fixed learning rate of 1e-3. We froze the pre-trained weights and fine-tuned the DeepGompertz head with a fixed learning rate of 1e-4 and a mini-batch size of 512. For both steps, an early-stopping strategy was applied with a patience of 5 epochs, monitoring the validation negative log-likelihood in the validation set (*D*_val_) to prevent overfitting, up to a maximum of 1,000 epochs.

### 4.6 Metabolomic age estimation

Metabolomic age was derived at inference by mapping each individual’s predicted mortality probability to the population age scale defined by the Gompertz baseline. At inference, *τ* is replaced by a fixed 10-year window, consistent with the window used in PhenoAge [1]. This fixed window ensures that mortality probabilities are comparable across individuals. The corresponding 10-year cumulative hazard for a reference individual of age (*x*_age_) is:

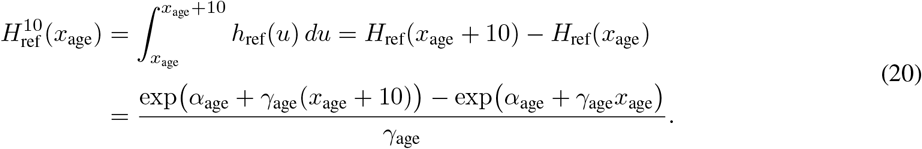

Throughout, *H*(·) denotes the cumulative hazard accrued from the origin of the corresponding time axis, as defined in **Equation 12**. The superscript 10 denotes the cumulative hazard accrued over a 10-year window, which starts at chronological age *x*_age_ for the Gompertz baseline and at enrollment for the DeepGompertz head. The mortality probabilities derived below are therefore conditional on survival to the start of that window.

This quantity was converted to a 10-year mortality probability by

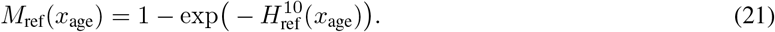

For individual *i* at inference, the 10-year cumulative hazard from enrollment was computed as

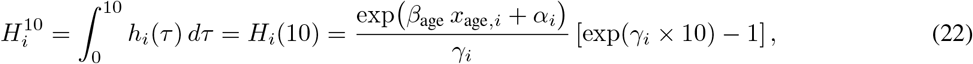

and the corresponding 10-year mortality probability was

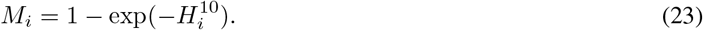

Metabolomic age (*x*_met_age,*i*_) was defined as the chronological age at which the Gompertz baseline (**Equation 20**) yields the same 10-year mortality probability as the individual prediction:

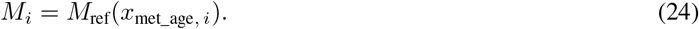

Equivalently, by substituting **Equation 20**, *x*_met_age,*i*_ satisfies

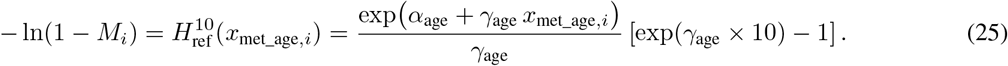

Solving for *x*_met_age,*i*_ gives

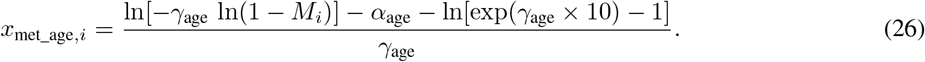

where *α*_age_ and *γ*_age_ are fitted parameters from the Gompertz baseline, *M*_*i*_ can be calculated from the output of the DeepGompertz head. Thus, *x*_met_age,*i*_ is the age on the Gompertz baseline scale that corresponds to the individualized 10-year mortality probability *M*_*i*_. When the individualized parameters (*α*_*i*_, *γ*_*i*_) predicted from metabolomic embeddings shift *M*_*i*_ away from the baseline probability expected at chronological age *x*_age,*i*_, the age gap captures the corresponding displacement along the reference age–mortality relationship. Chronological age at enrollment (*x*_age,*i*_) enters the DeepGompertz hazard through the population-shared coefficient *β*_age_, and the term (*β*_age_ *x*_age,*i*_) accounts for the cohort-wide increase in hazard with chronological age. This age-related hazard adjustment is combined with the individualized parameters (*α*_*i*_, *γ*_*i*_) from metabolomic embeddings to determine *M*_*i*_, and the conversion to *x*_met_age,*i*_ expresses this combined 10-year mortality probability as an equivalent age under the reference age–mortality relationship.

### 4.7 Identification of metabolic subtypes

Metabolic subtypes were identified using the Leiden clustering algorithm applied to the raw metabolomic embeddings from the training set (*D*_train_). We searched a grid of the number of neighbors (`n_neighbors` ∈ {5, 10, 15}) and resolution values (`resolution` ∈ {0.5, 1.0, 2.0}) and selected the Leiden clustering setting that maximized the minimum cluster size. The neighbor graph for Leiden clustering was constructed with cosine distance and the selected parameters of n_neighbors=15, and clustering was performed at the parameter of resolution=1, yielding 13 data-driven metabolic subtypes. To enable subtype assignment for new samples, we fitted a MLP classifier with focal loss [68] (focal *γ* = 2.0, class-balanced loss weighting, and a balanced mini-batch sampler; PyTorch implementation) using the embeddings from the training set and their corresponding subtype labels derived from Leiden clustering, with the following parameters: hidden_layer_sizes=(128, 64), activation=‘relu’, dropout=0.1, batch_size=2,048, learning_rate=10^−3^, max_epochs=500, validation_fraction=0.1, patience=30, random_state=42, early_stopping=True. This classifier allowed any new sample (e.g., the hold-out test set and participants from the CHARLS cohort) with a metabolomic embedding to be assigned to one of the identified subtypes. For downstream analyses requiring broader groupings, the 13 MLP-assigned subtypes were merged into four meta-subtypes according to a predefined mapping informed by hierarchical clustering of subtype-specific disease incidence profiles. For visualization purposes, UMAP dimension reduction was applied to the embeddings with the following parameters: n_neighbors=15, min_dist=0.5, spread=1.0, metric=‘cosine’, and n_components=2.

### 4.8 Distillation for the lightweight model

A total of 14 clinical blood biomarkers were used to distill the lightweight model, including 5 blood cell counts (white blood cells, haemoglobin concentration, haematocrit percentage, mean corpuscular volume, and platelet count from UKB Category 100081) and 9 biochemistry measures (C-reactive protein, glycated haemoglobin, cholesterol, HDL cholesterol, triglycerides, creatinine, glucose, urate, and Cystatin C from UKB Category 17518). The architecture of the lightweight model is built on transformer-encoder layers (**Supplementary Figure S1E**), which maps clinical biomarkers **x**_clin_ ∈ ℝ^14^ into a dense embedding **h**_clin_ ∈ ℝ^512^. Each scalar biomarker value is linearly projected into a 512-dimensional embedding and added to a learnable positional embedding. Missing values are encoded by a learnable mask embedding rather than imputation. Similar to the pre-trained MetAgeFormer, a CLS embedding is prepended to the biomarker embeddings and updated by two transformer-encoder layers (4 self-attention heads, GELU feed-forward with 1,024 dimensions, and dropout rate of 0.1). The model forward pass can be formulated as

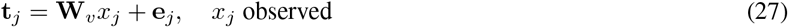

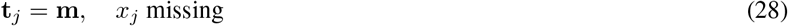

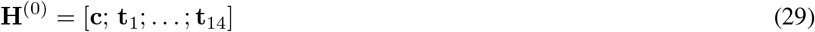

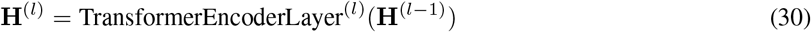

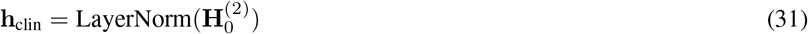

**x**_clin_ denotes the input vector of 14 clinical biomarkers, and *x*_*j*_ is the *j*-th biomarker value. **W**_*v*_ ∈ ℝ^512^ is the shared linear projection weight for observed values; **e**_*j*_ ∈ ℝ^*d*^ is the learnable positional embedding for the *j*-th biomarker; **m** ∈ ℝ^512^ is the learnable mask embedding for missing values; and **c** ∈ ℝ^512^ is the CLS embedding. **t**_*j*_ is the token embedding for the *j*-th biomarker. **H**^(0)^ represents the input to the first transformer-encoder layer, formed by concatenating the CLS embedding with the 14 biomarker token embeddings. **H**^(*l*)^ represents the output embeddings of the *l*-th transformer-encoder layer. 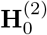 denotes the updated CLS embedding at the final layer, and **h**_clin_ is its layer-normalized representation used as the clinical embedding.

The mask embedding **m** is shared across biomarkers and, unlike observed tokens, does not receive a positional embedding. Missing tokens are additionally masked out as attention keys, so they are never attended to by other tokens and make no direct contribution to the updated CLS embedding. Consequently, the missing pattern of an individual is conveyed implicitly, through which positional embeddings are present among the attended tokens, rather than through the content of the mask tokens themselves. Formally, the self-attention in **Equation 30** is computed with a key-padding mask ℳ_*i*_ = {*j* : *x*_*j*_ is missing}, and attention weights to positions in ℳ_*i*_ are set to zero. This design keeps the token sequence at a fixed length (one CLS embedding and 14 biomarker tokens).

To align the dense embedding space generated from the lightweight model with the metabolomic representations learned by MetAgeFormer, we employ a distillation objective combining symmetric InfoNCE loss for embedding alignment (**Figure 2C**). Additionally, we added a DeepGompertz negative log-likelihood loss 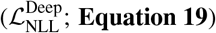 to ensure that metabolomic age and mortality risk predicted by the lightweight model remain consistent with those of the original MetAgeFormer, allowing seamless reuse of the fine-tuned DeepGompertz head. The distillation objective is formulated as:

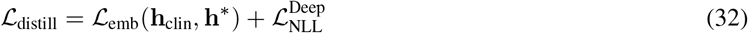

where 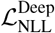 is computed by passing **h**_clin_ through the teacher-initialized DeepGompertz head, which was kept frozen during distillation. ℒ_emb_ was implemented by symmetric InfoNCE loss with temperature *τ* = 0.2, which is deliberately chosen over alternative metric-learning objectives because we aim to enable clinical biomarkers to faithfully mimic the metabolomic embedding geometry, instead of merely preserving pairwise similarities. Model optimization was implemented using a fixed learning rate of 10^−4^, a mini-batch size of 512, and an early-stopping strategy with a patience of 10 epochs, monitored via the distillation loss (ℒ_distill_) in the validation set (*D*_val_), up to a maximum of 300 epochs.

### 4.9 Comparison of aging clocks and markers

We implemented MileAge following the chronological age regression protocol of the original publication [5] using the scikit-learn Python library [69]. Metabolite predictors comprised 168 Nightingale NMR biomarkers selected to match the original MileAge predictor set. Missing values were imputed by mean values for each metabolite. Metabolite concentrations were standardized using mean and standard deviations from the training set. We established a diverse MileAge variants based on a set of machine learning models including LASSO, Ridge, ElasticNet, k-nearest neighbours (kNN), support vector machine (SVM) regressors with linear, polynomial, and radial kernels, regression trees, random forests, and XGBoost. Hyperparameters were optimized using a randomly selected 10% subsample of the training set (*D*_train_) through 3-fold cross-validation with mean absolute error as the selection metric. Subsequently, models were re-trained on the complete training set using the optimal hyperparameters. Specifically, we employed grid search for penalized linear models. For LASSO, *α* spanned 20 logarithmically spaced values from 10^−4^ to 10^1^. For Ridge, *α* spanned 25 logarithmically spaced values from 10^−4^ to 10^4^. For ElasticNet, *α* spanned 12 logarithmically spaced values from 10^−4^ to 10^1^ with l1_ratio ∈ {0.2, 0.5, 0.7, 0.9}. For kNN, we searched over n_neighbors ∈ {5, 10, 25, 50}, weights ∈ {uniform, distance}, and Minkowski distance exponents *p* ∈ {1, 2}. For linear SVM regressors (LinearSVR), we searched over *C* ∈ {0.01, 0.1, 1.0, 10.0} and *ϵ* ∈ {0.0, 0.1, 0.5}. For polynomial and radial SVM regressors, we used a Nyström kernel approximation coupled with stochastic gradient descent and conducted randomized search over 12 configurations. For polynomial SVM, the search space covered n_components ∈ {100, 300, 600}, degree ∈ {2, 3}, *γ* ∈ {0.01, 0.1, 1.0}, and L2 regularization *α* ∈ {10^−5^, 10^−4^, 10^−3^}. For radial SVM, the search space covered n_components ∈ {100, 300, 600}, *γ* ∈ {0.001, 0.01, 0.1, 1.0}, and L2 regularization *α* ∈ {10^−5^, 10^−4^, 10^−3^}. For regression trees, we conducted randomized search over 20 configurations with max_depth ∈ {3, 5, 10, 20, unrestricted}, min_samples_leaf ∈ {10, 50, 100, 500}, and min_samples_split ∈ {20, 100, 500}. For random forests, we conducted randomized search over 20 configurations with n_estimators drawn uniformly from integers in [100, 500), max_depth ∈ {10, 20, 30, unrestricted}, min_samples_split drawn uniformly from integers in [20, 500), min_samples_leaf drawn uniformly from integers in [5, 100), and max_features drawn uniformly from [0.3, 1.0]. For XGBoost, we conducted randomized search over 20 configurations with n_estimators drawn uniformly from integers in [100, 600), max_depth drawn uniformly from integers in [2, 8), learning_rate drawn uniformly from [0.01, 0.20], subsample and colsample_bytree drawn uniformly from [0.6, 1.0], and reg_lambda ∈ {0.1, 1.0, 10.0}. Predicted age was converted to metabolomic age using linear regression bias correction fitted on the combined training and validation sets. Age acceleration was defined as the difference between bias-corrected metabolomic age and chronological age, and this definition was applied consistently across all MileAge models. Beyond MileAge, we included additional published aging clocks and clinical biomarkers for comparison. Phenotypic age (PhenoAge) was computed using the pyaging implementation [1] from nine blood biochemistry and blood count biomarkers together with chronological age. The Deelen metabolic mortality score was computed using the MiMIR R package [19] from 14 Nightingale metabolites, and the resulting mortality score was used as the aging metric. Frailty Index was computed at baseline (instance 0) using the ‘di’ R package https://CRAN.R-project.org/package=di from 49 deficit items. Single-marker clinical aging proxies were derived from Cystatin C and glycated haemoglobin (HbA1c) measured in UKB blood biochemistry. Rather than re-training from scratch, we directly implemented the trained weights of ProteomicAge (Hamilton) as established in the original study [3]. The pre-processing of proteomic profiles, including quality control and missing value imputation, strictly adhered to the protocols defined in the original study [3], resulting in 2917 proteins for deriving proteomic age acceleration. To ensure a rigorous and unbiased comparison, our analysis was restricted to a shared test set of 2,679 participants who were entirely excluded from the training stages of both MetAgeFormer and ProteomicAge (Hamilton) (**Supplementary Figure S4**).

For each head-to-head comparison, analyses were restricted to the UK Biobank hold-out test set samples with non-missing MetAgeFormer metabolomic age acceleration and each competitor under evaluation. Because competitors draw on different UK Biobank fields (e.g., routine blood biochemistry, frailty phenotypes, published aging clocks, proteomic aging scores), sample sizes differ across comparison groups and should not be compared directly between panels. Sample and event counts for every disease endpoint and comparison are reported in **Supplementary Table S2**.

### 4.10 Statistical analysis

To analyze disease onset, we performed Cox proportional hazards regression using the CoxPHFitter function from the lifelines Python package (https://lifelines.readthedocs.io/en/latest/fitters/regression/CoxPHFitter.html). Linear associations were assessed via Ordinary Least Squares (OLS) regression using the statsmodels package https://www.statsmodels.org/stable/generated/statsmodels.regression.linear_model.OLS.html, while associations between pairs matching and metabolic subtypes were evaluated using Zero-inflated Poisson regression (https://www.statsmodels.org/stable/generated/statsmodels.discrete.count_model.ZeroInflatedPoisson.html) to account for the high proportion of participants with zero matching errors. Cumulative incidence curves for age-related diseases were generated using the KaplanMeierFitter within the lifelines framework (https://lifelines.readthedocs.io/en/latest/fitters/univariate/KaplanMeierFitter.html). The effect sizes of differential metabolites across subtypes were quantified using Cohen’s *d*. For multiple testing, p-values were adjusted using the Bonferroni correction.

To minimize reverse causation bias in disease onset analyses, participants with prevalent diseases at baseline were excluded. For the analysis of memory-related cognitive functions, we excluded participants with all-cause dementia from the UKB cohort and those with memory-related diseases from the CHARLS cohort at baseline. In subtype-specific analyses, the target subtype was encoded as 1, with all other subtypes pooled and encoded as 0 to serve as the general population reference. Head-to-head comparisons of aging clocks and markers for all-cause mortality and incident disease (**Figure 3** and **Figure 4**) were adjusted for chronological age and sex, following the evaluation protocol of Hamilton et al. [3]. All subtype-, meta-subtype- and ΔAge-trajectory-level analyses of incident disease, in both UK Biobank and CHARLS (**Figure 5B,C, and H, Extended Data Figure 5B and I**), were additionally adjusted for BMI, because BMI differs systematically across metabolic subtypes (**Figure 5E**) and is therefore a potential confounder of subtype–outcome associations. Associations in ADNI were adjusted for chronological age and sex. The CHARLS cognitive analyses (Figure 6G–I) were adjusted for chronological age, sex, and BMI. No other covariates were included unless otherwise specified.

## Data availability

The source code of MetAgeFormer is publicly available at the GitHub repository (https://github.com/ericcombiolab/MetAgeFormer). UKB NMR metabolomic profiles and participants’ data can be assessed from the UKB-RAP (https://ukbiobank.dnanexus.com/landing) through approved research projects. ADNI data can be accessed (https://adni.loni.usc.edu/) through the approved request. CHARLS data is accessible (https://charls.pku.edu.cn/en/) through the approved research request.

## Competing interests

The authors declare that they have no competing interests.

## Author contributions statement

LZ, WJ, and YX conceived the study; YX designed MetAgeFormer, implemented the algorithm, and conducted the experiments; YX, LZ, and WJ analyzed the results; YX wrote the original article; LZ and WJ reviewed and edited the paper; BHZ contributed to discussions regarding the design of MetAgeFormer; GXX contributed to discussions concerning the results. TLC contributed to discussions regarding the external validation data. All authors read and approved the final manuscript.

## Acknowledgments

This work was supported by Young Collaborative Research grant (No. C2004-23Y and C2005-24Y), HMRF grant (No. 11221026), HKBU RCMS (RCMS/24-25/03). We gratefully acknowledge funding support from the Shenzhen-Hong Kong Jointly Funded Project from Shenzhen Science and Technology Program (Category A; SGDX20230116093201002). This study analyzed data provided by UK Biobank via application 60434. We extend our sincere thanks to all participants and their families, as well as to the investigators and staff of UK Biobank. The metabolomics data of ADNI cohort were provided by the Alzheimer’s Disease Metabolomics Consortium (ADMC). As such, the investigators within the ADMC, not listed specifically in this publication’s author’s list, provided data along with its pre-processing and prepared it for analyses, but did not participate in analyses or writing of this manuscript. A complete listing of ADMC investigators can be found at: https://sites.duke.edu/adnimetab/team/. Sample and information collection for ADNI cohorts was funded by the Alzheimer’s Disease Neuroimaging Initiative (ADNI) (National Institutes of Health Grant U01 AG024904) and DOD ADNI (Department of Defense award number W81XWH-12-2-0012). ADNI is funded by the National Institute on Aging, the National Institute of Biomedical Imaging and Bioengineering, and through generous contributions from the following: AbbVie, Alzheimer’s Association; Alzheimer’s Drug Discovery Foundation; Araclon Biotech; BioClinica, Inc.; Biogen; Bristol-Myers Squibb Company; CereSpir, Inc.; Eisai Inc.; Elan Pharmaceuticals, Inc.; Eli Lilly and Company; EuroImmun; F. Hoffmann-La Roche Ltd and its affiliated company Genentech, Inc.; Fujirebio; GE Healthcare; IXICO Ltd.; Janssen Alzheimer Immunotherapy Research & Development, LLC.; Johnson & Johnson Pharmaceutical Research & Development LLC.; Lumosity; Lundbeck; Merck & Co., Inc.; Meso Scale Diagnostics, LLC.; NeuroRx Research; Neurotrack Technologies; Novartis Pharmaceuticals Corporation; Pfizer Inc.; Piramal Imaging; Servier; Takeda Pharmaceutical Company; and Transition Therapeutics. The Canadian Institutes of Health Research is providing funds to support ADNI clinical sites in Canada. Private sector contributions are facilitated by the Foundation for the National Institutes of Health (https://fnih.org). The grantee organization is the Northern California Institute for Research and Education, and the study is coordinated by the Alzheimer’s Disease Cooperative Study at the University of California, San Diego. Data used in preparation of this article were obtained from the Alzheimer’s Disease Neuroimaging Initiative (ADNI) database (https://adni.loni.usc.edu). As such, the investigators within the ADNI contributed to the design and implementation of ADNI and/or provided data but did not participate in analyses or writing of this report. A complete listing of ADNI investigators can be found at: http://adni.loni.usc.edu/wp-content/uploads/how_to_apply/ADNI_Acknowledgement_List.pdf.

---

**Extended Data Figure 1.**
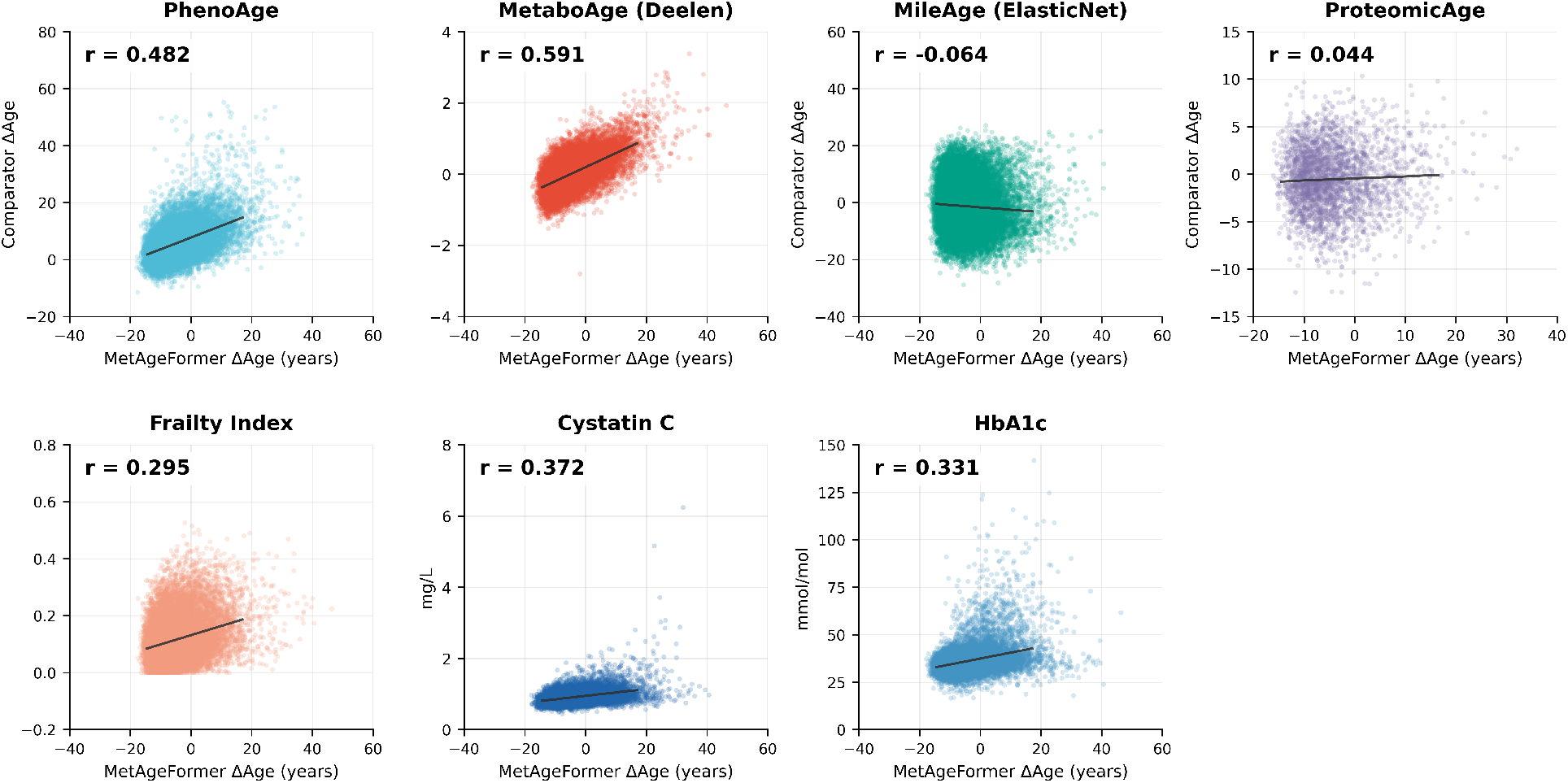
Correlation of MetAgeFormer ΔAge with competitor aging measures and biomarkers in the UKB test set. Each panel shows MetAgeFormer ΔAge (x-axis) versus the indicated competitor (y-axis), with a line fitted by ordinary least-squares regression and Pearson r. Sample sizes vary by competitor availability.

**Extended Data Figure 2.**
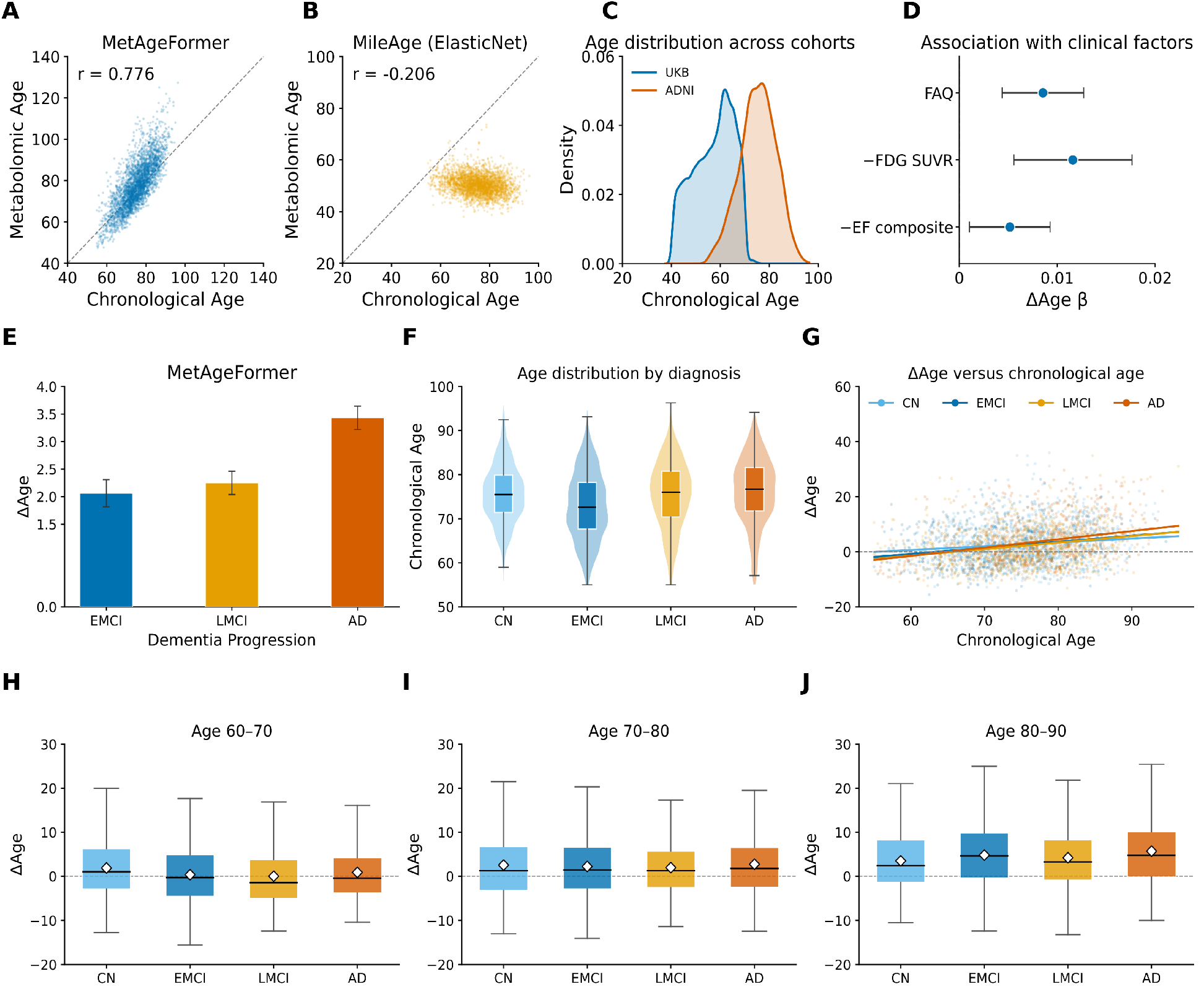
Zero-shot external validation of MetAgeFormer in ADNI. **(A-B)** Predicted metabolomic age vs. chronological age for MetAgeFormer and MileAge (ElasticNet). **(C)** Chronological age distributions in UKB and ADNI cohorts. **(D)** Associations of ΔAge with FAQ, FDG SUVR, and the EF composite. Points show ΔAge *β* with 95% confidence intervals, adjusted for chronological age and sex. **(E)** Mean ΔAge across dementia progression (EMCI, LMCI, and AD). **(F)** Chronological age distributions by diagnosis (CN, EMCI, LMCI, and AD). **(G)** ΔAge versus chronological age by diagnosis. **(H–J)** Diagnosis-stratified ΔAge within chronological age windows 60–70, 70–80, and 80–90 years. Black lines and white diamonds indicate medians and means, respectively.

**Extended Data Figure 3.**
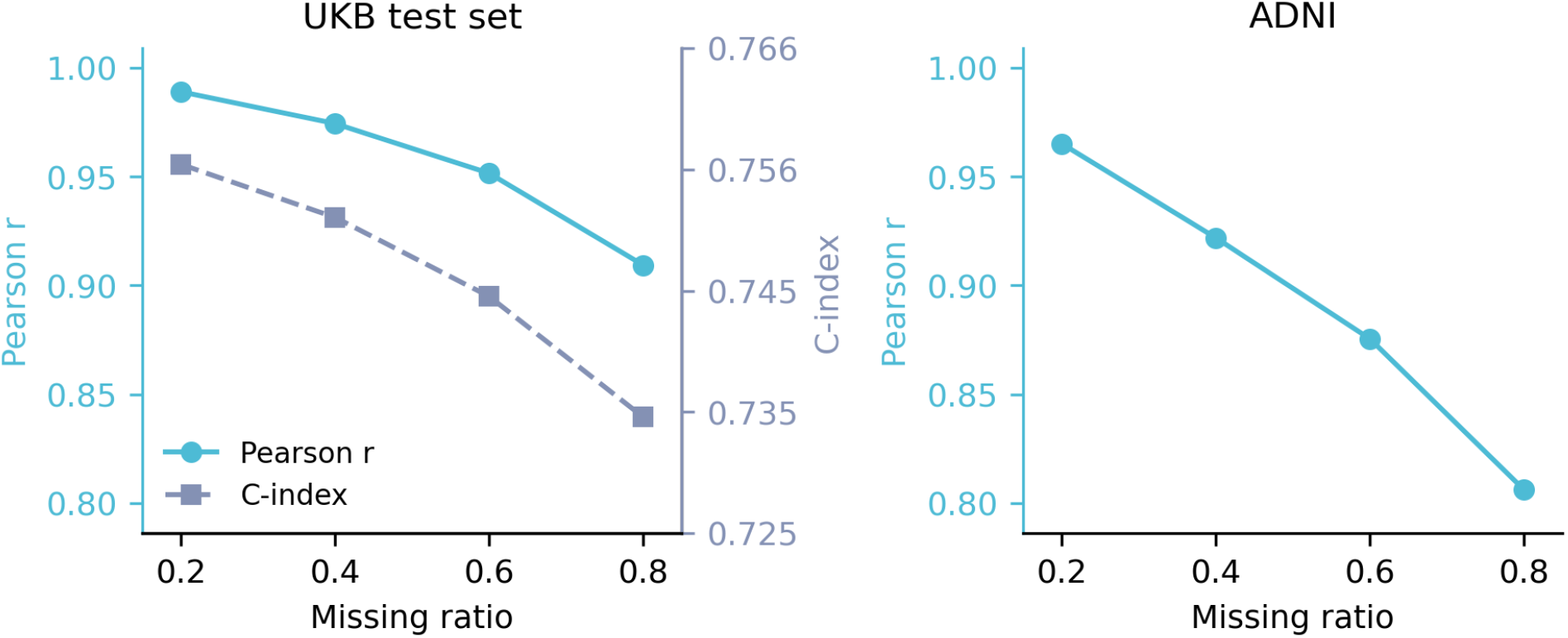
Robustness of MetAgeFormer metabolomic age to simulated missing metabolites. **(Left)** UKB hold-out test set (73,417 plasma samples): Pearson correlation between metabolomic age under masked inputs and metabolomic age under the full metabolite panel (left axis), and mortality C-index (right axis), as a function of missing ratio. **(Right)** ADNI (4,851 plasma samples from 1,524 participants): corresponding Pearson correlation with full-panel metabolomic age.

**Extended Data Figure 4.**
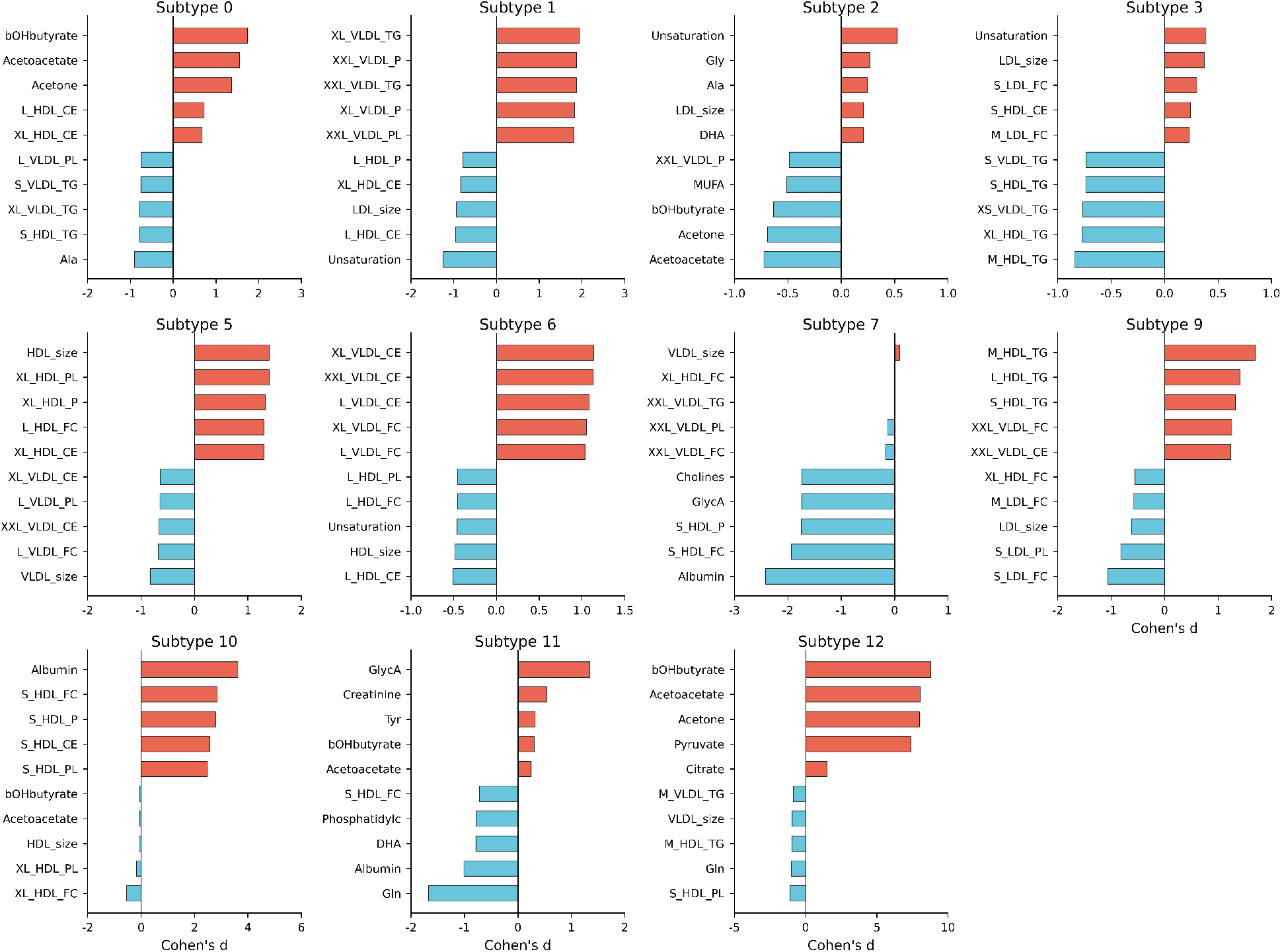
Bar plots of the top 10 significantly differential metabolites (5 positives and 5 negatives) for metabolic subtypes. The x-axis represents Cohen’s d effect sizes, and the y-axis represents metabolite abbreviations. Subtypes 4 and 8 are shown in **Figure 5D**.

**Extended Data Figure 5.**
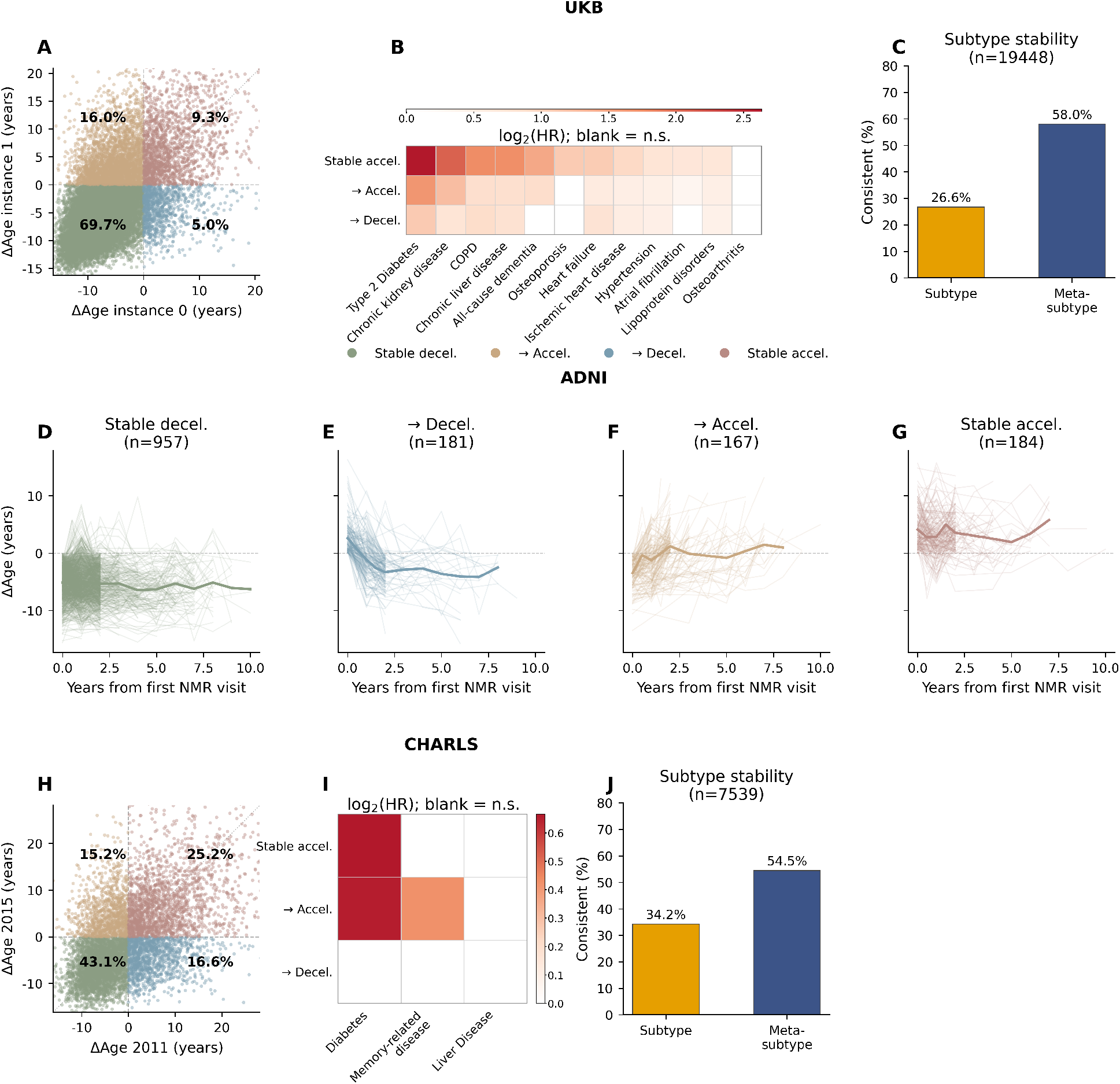
Longitudinal dynamics of metabolic age acceleration and health outcomes across the UK Biobank (UKB), ADNI, and CHARLS cohorts. **(A)** Scatter plot of MetAgeFormer-derived ΔAge trajectories between instance 0 (x-axis) and instance 1 (y-axis) in UKB (n=19,448). Four quadrants define distinct aging patterns: Stable Decelerated (both timepoints ΔAge *<* 0), Transition to Accelerated (ΔAge *<* 0 at instance 0 and ΔAge ≥ 0 at instance 1), Transition to Decelerated (ΔAge ≥ 0 at instance 0 and ΔAge *<* 0 at instance 1), and Stable Accelerated (both timepoints ΔAge *>* 0). Quadrant percentages are shown; colors denote acceleration group (green, Stable decelerated; orange, Transition to accelerated; blue, Transition to decelerated; red, Stable accelerated). **(B)** Heatmap of log_2_(hazard ratios) from Cox proportional-hazards models for incident disease risk across non-reference acceleration groups (reference: Stable Decelerated), adjusted for age, sex, and BMI at instance 1 among participants disease-free at both instances; diseases are ordered by maximum significant log_2_(HR). Cells are colored only for significant associations (*p <* 0.05); blank cells indicate non-significance. **(C)** Metabolic subtype stability between instances 0 and 1, shown as the percentage of participants with consistent subtype or meta-subtype assignment at both time points. **(D–G)** Individual ΔAge trajectories in ADNI, stratified by the same four acceleration groups defined from first-to-last paired NMR visits (n=957, 181, 167, and 184, respectively). Thin lines show subject-level trajectories (≥ 2 visits); thick lines show group means. **(H–J)** Parallel analyses in CHARLS (n=7,539): **(H)** ΔAge scatter between 2011 and 2015 with the same quadrant definitions; **(I)** Cox heatmap (log_2_ HR) for incident diabetes, memory-related disease, and liver disease (reference: Stable Decelerated; covariates: age, sex, and BMI at 2015); **(J)** Subtype and meta-subtype stability between 2011 and 2015.

## Notes

### Competing Interest Statement

The authors have declared no competing interest.

### Summary of Updates

The major changes include: (1) renaming the model from MetFoundation to MetAgeFormer to describe its scope more accurately; (2) a redesigned DeepGompertz survival module that separates the shared chronological age effect from metabolome-specific risk; (3) external validation in the ADNI cohort using directly measured Nightingale NMR data, including cross-platform validation against an independent LC-MS assay; (4) head-to-head benchmarking against established aging clocks and clinical biomarkers (PhenoAge, MetaboAge (Deelen), ProteomicAge (Hamilton), Frailty Index, Cystatin C, and HbA1c), together with calibration and decision-curve analyses; (5) comprehensive ablation and sensitivity analyses covering pre-training, clustering parameters, covariate adjustment, and acetate robustness; and (6) tempered language regarding clinical utility and mechanistic interpretation throughout.

