## Supplementary Data 1 for "Decoding heterogeneous aging clocks and disease risk stratification using MetAgeFormer"

**Layer-wise attention connection atlas for MetAgeFormer**

**Supplementary Data 1 | Layer-wise attention connection atlas for MetAgeFormer.**

Arc diagrams visualize self-attention connectivity among the 107 non-derived NMR metabolites in the pre-trained MetAgeFormer encoder (six transformer-encoder layers; eight attention heads per layer). For each head, metabolites are ordered along the horizontal axis (blue markers). Red arcs connect each query metabolite to its top three attended metabolites; arc opacity and line width are scaled by the corresponding attention weight. The [CLS] token was excluded from all plots. Each page shows one transformer-encoder layer; within each page, attention heads 1–8 are arranged top to bottom. Metabolite names are shown on the bottom head panel only. These full-resolution layer-wise maps complement the cross-layer summary in Supplementary Figure 3 and are provided for illustrative inspection of head- and layer-specific connectivity patterns.

**Layer 1**

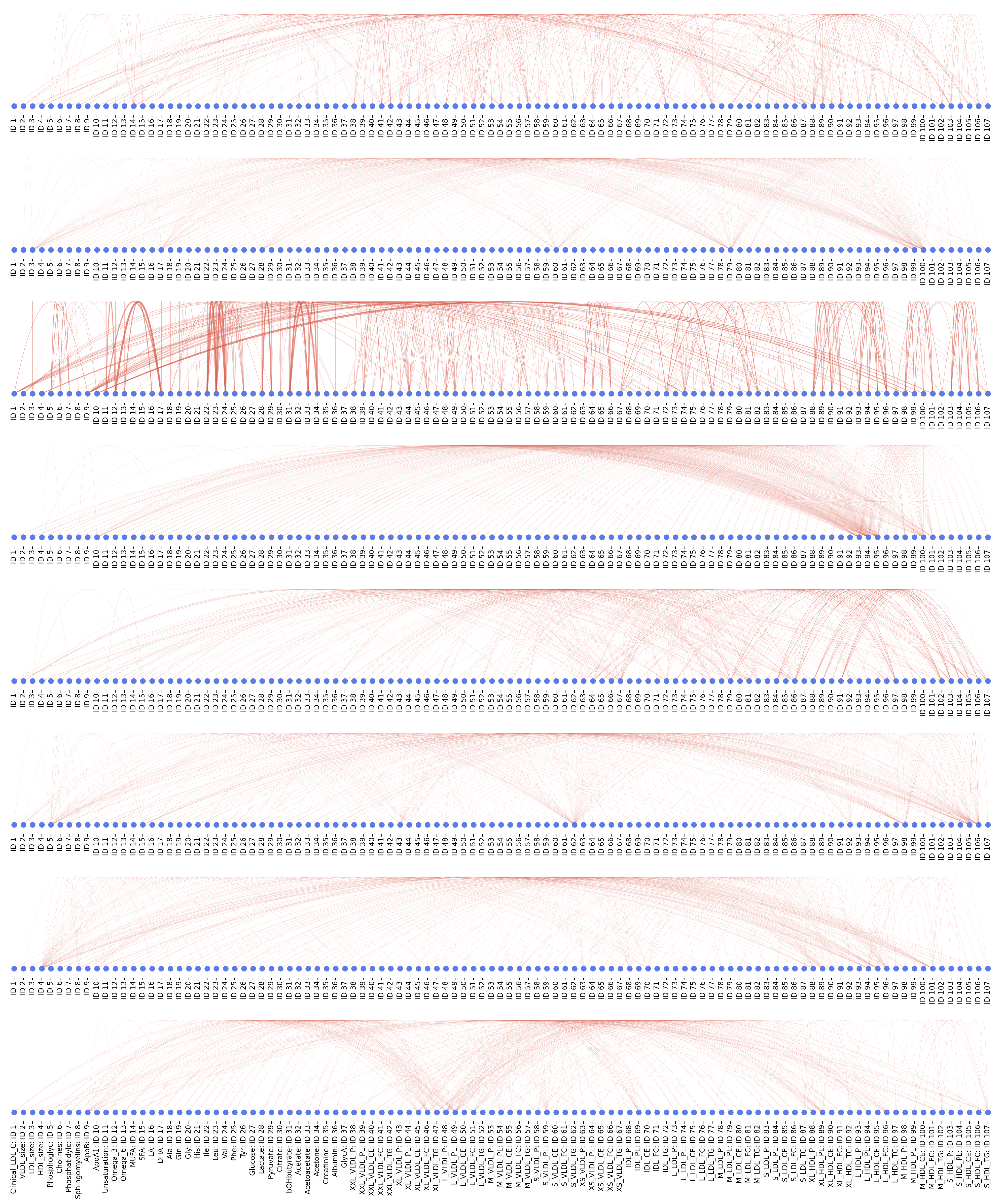

*Layer 1. Attention connection diagrams for heads 1–8 (top to bottom) in transformer-encoder layer 1 of MetAgeFormer. See Supplementary Data 1 legend for details.*

**Layer 2**

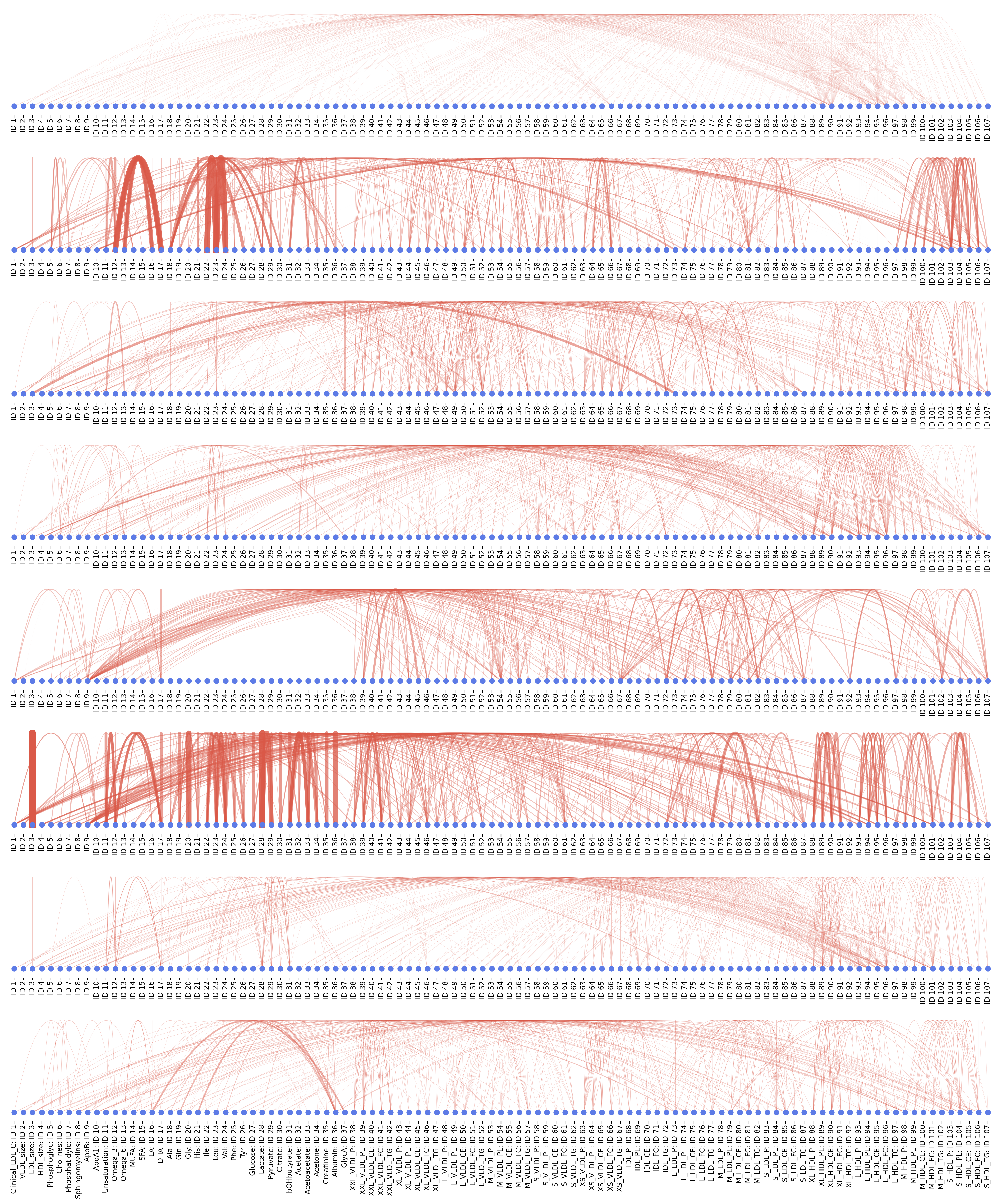

*Layer 2. Attention connection diagrams for heads 1–8 (top to bottom) in transformer-encoder layer 2 of MetAgeFormer. See Supplementary Data 1 legend for details.*

**Layer 3**

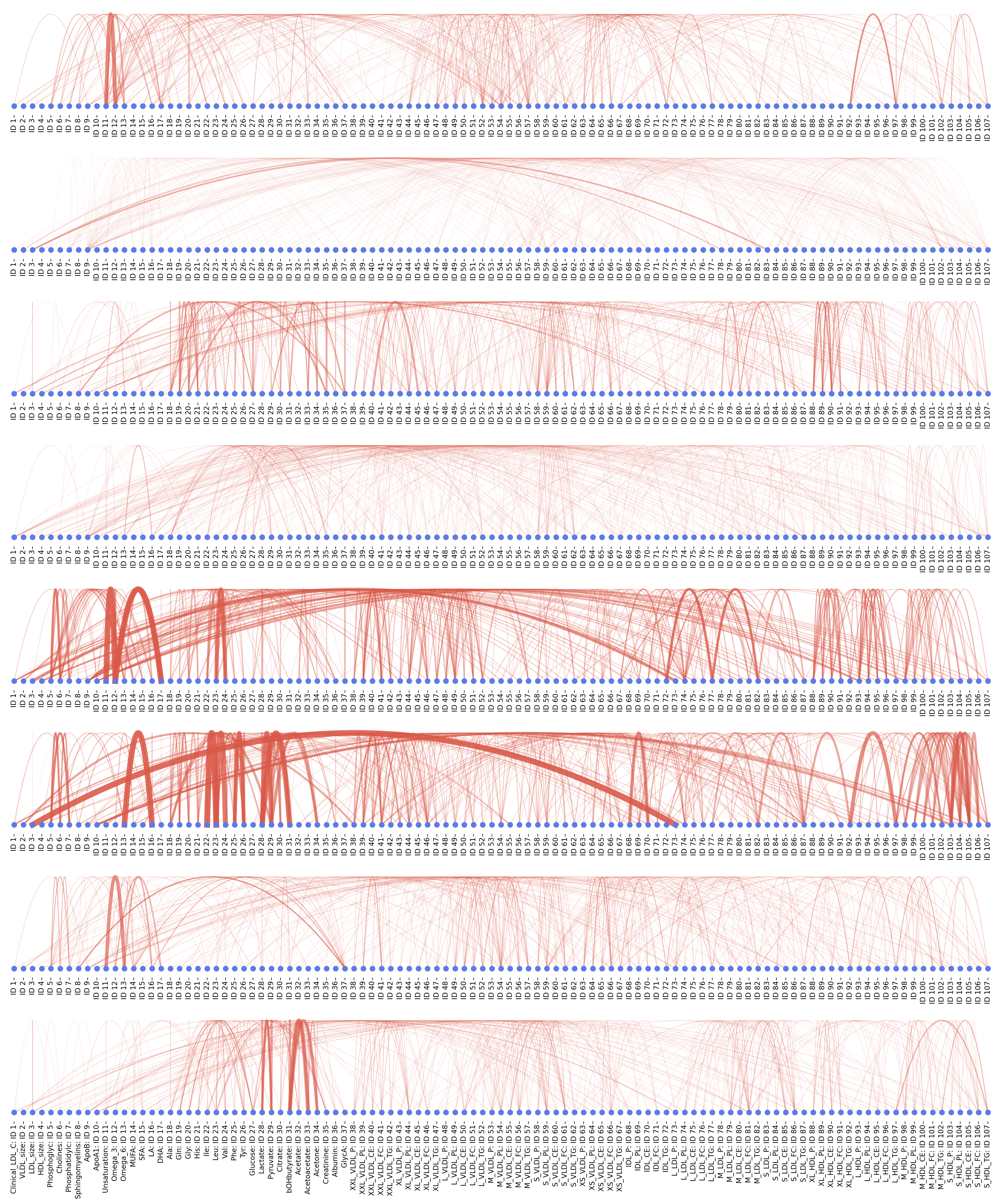

*Layer 3. Attention connection diagrams for heads 1–8 (top to bottom) in transformer-encoder layer 3 of MetAgeFormer. See Supplementary Data 1 legend for details.*

**Layer 4**

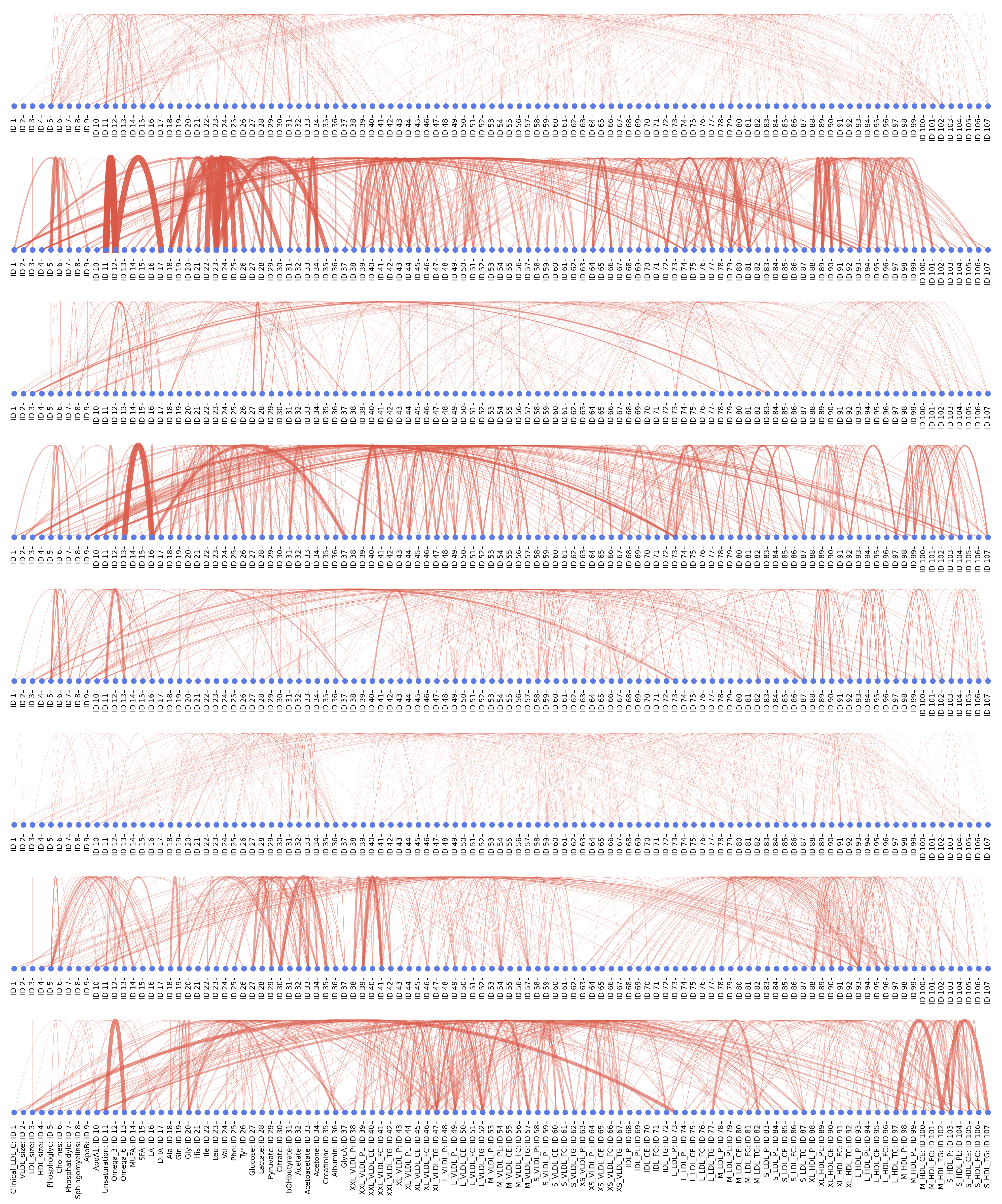

*Layer 4. Attention connection diagrams for heads 1–8 (top to bottom) in transformer-encoder layer 4 of MetAgeFormer. See Supplementary Data 1 legend for details.*

**Layer 5**

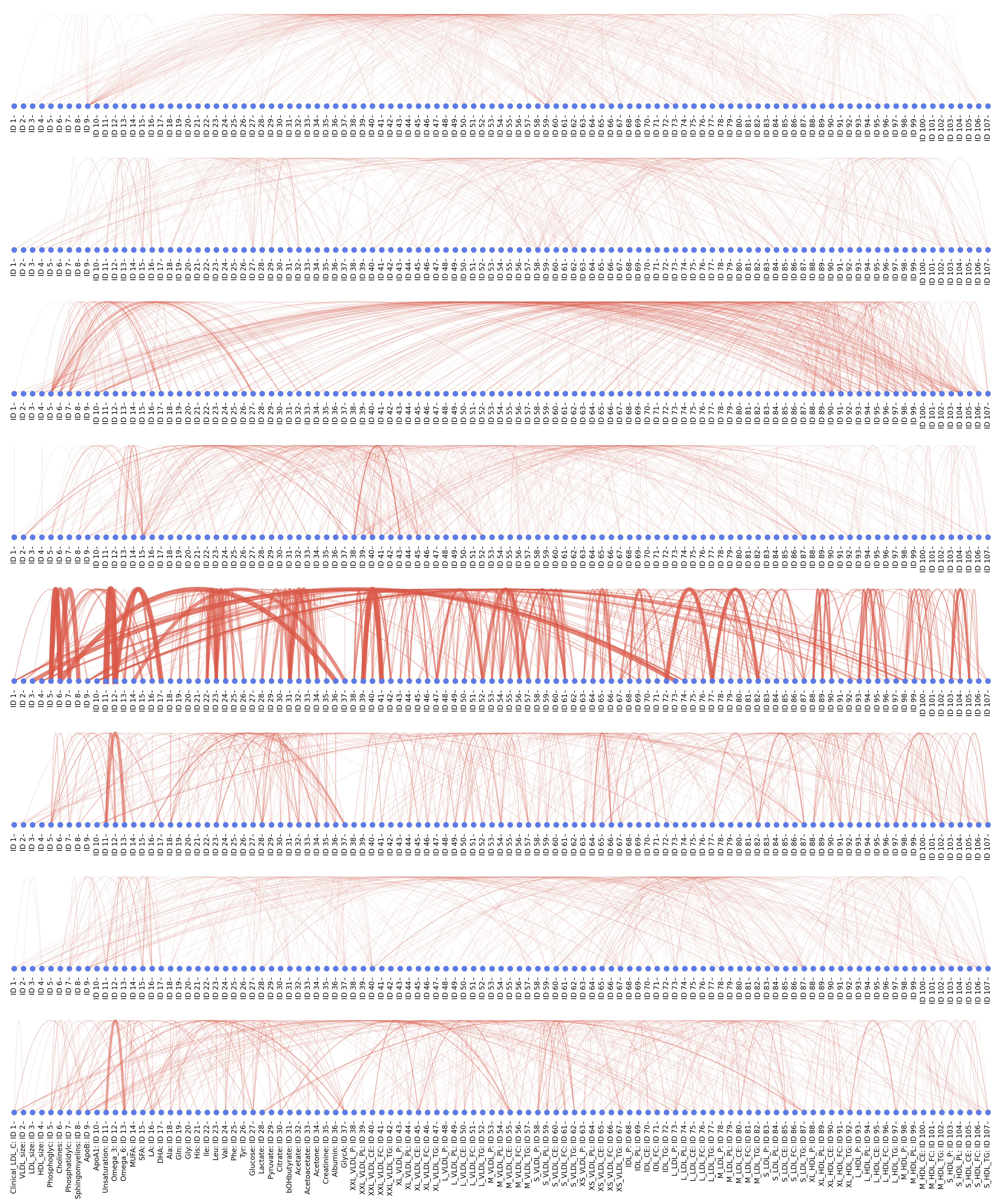

*Layer 5. Attention connection diagrams for heads 1–8 (top to bottom) in transformer-encoder layer 5 of MetAgeFormer. See Supplementary Data 1 legend for details.*

**Layer 6**

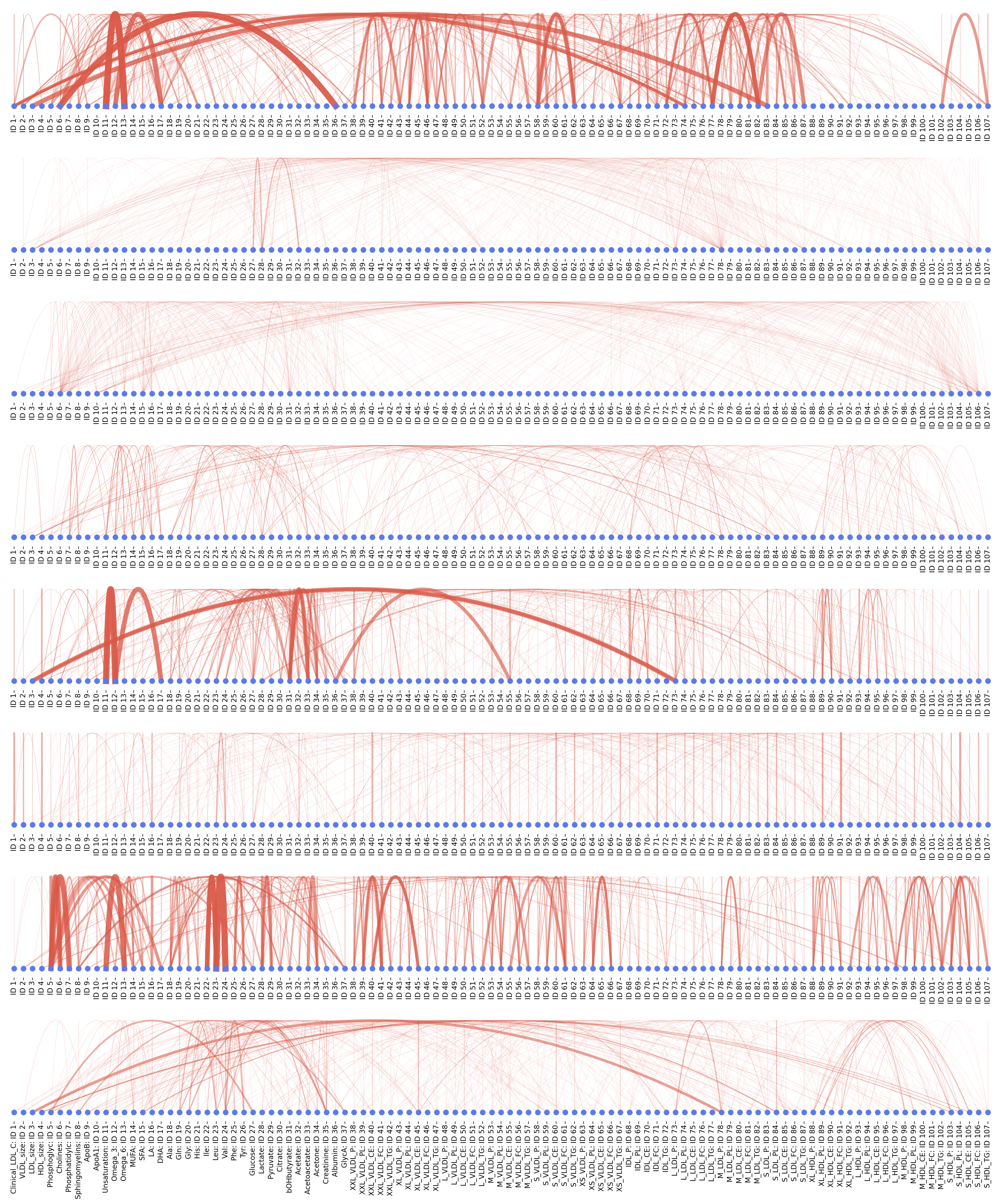

*Layer 6. Attention connection diagrams for heads 1–8 (top to bottom) in transformer-encoder layer 6 of MetAgeFormer. See Supplementary Data 1 legend for details.*
