## Supplementary Data 2 for "Decoding heterogeneous aging clocks and disease risk stratification using MetAgeFormer"

Concentration distribution comparisons for clinical blood biomarkers, NMR metabolites across UK Biobank and ADNI, and overlapping metabolites measured by NMR and LC-MS within ADNI.

### Clinical blood biomarkers · UK Biobank vs CHARLS

Distribution of blood biomarker concentrations across UK Biobank and CHARLS cohorts. Density-normalized histograms showing the distribution of 14 blood biomarkers measured in three cohorts: UK Biobank (blue, n=434,366), CHARLS 2011 (tan, n=11,823), and CHARLS 2015 (green, n=13,257). Each subplot represents one biomarker with measurement units indicated in parentheses on the x-axis. Statistical comparisons between cohorts were performed using two-sided Wilcoxon rank sum tests, with Bonferroni correction applied for three pairwise comparisons per biomarker. P-values are displayed in the top-right corner of each subplot: p<0.001, p<0.01, and p<0.05 denote statistical significance after correction. Otherwise, exact adjusted p-values are shown.

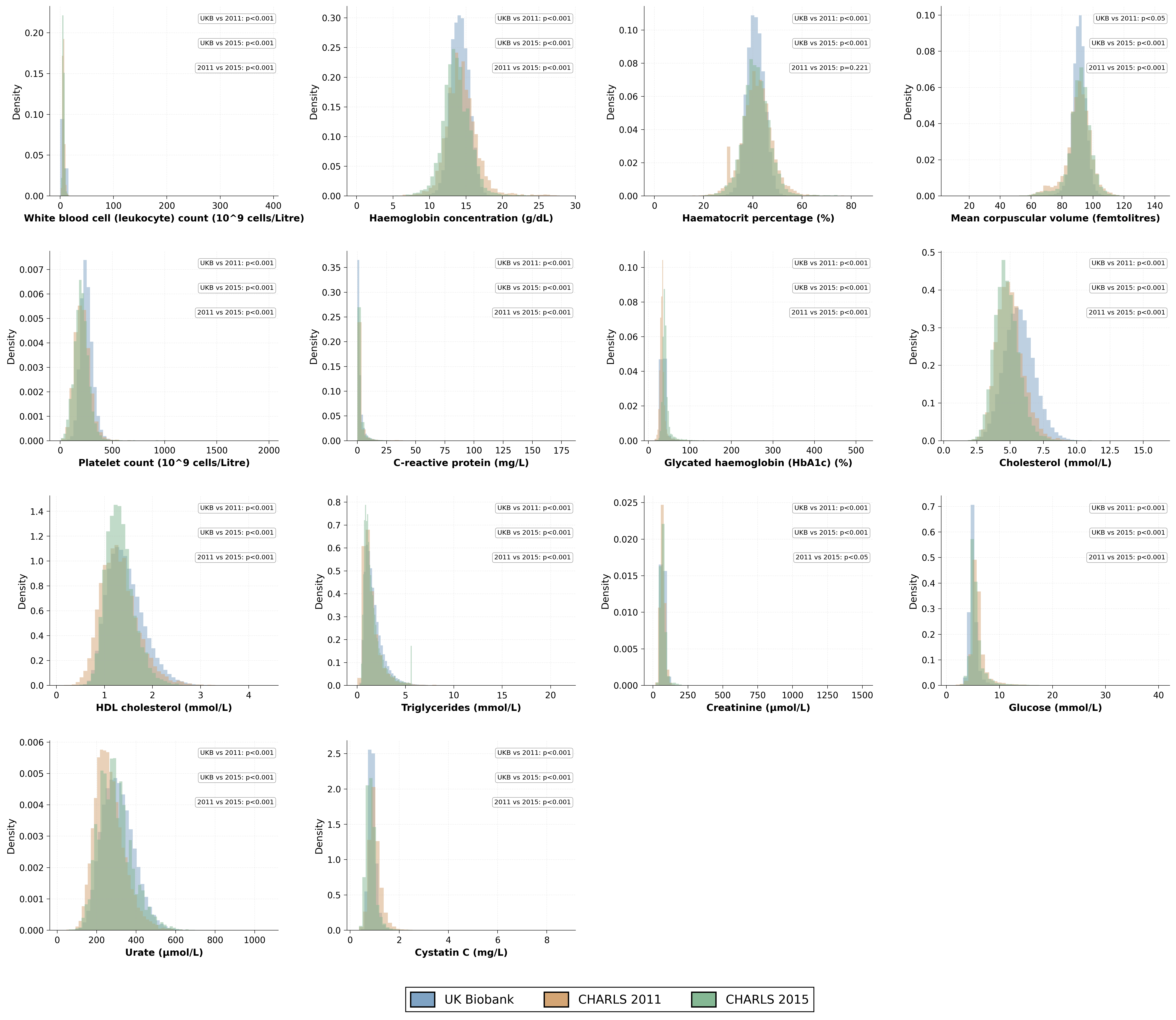

### NMR metabolites · UK Biobank vs ADNI (part 1 of 5)

Distribution of NMR metabolite concentrations across UK Biobank and ADNI cohorts (part 1 of 5). Density-normalized histograms showing 24 non-derived Nightingale NMR metabolites measured in two cohorts: UK Biobank (blue, n=507,783) and ADNI (orange, n=4,851). Each subplot represents one metabolite (metabolites 1–24 of 107) with concentration in mmol/L on the x-axis. Metabolites are shown in UK Biobank panel order. Statistical comparisons were performed using two-sided Wilcoxon rank sum tests, with Bonferroni correction applied across all 107 metabolites. P-values are displayed in the top-right corner of each subplot: p<0.001, p<0.01, and p<0.05 denote statistical significance after correction. Otherwise, exact adjusted p-values are shown.

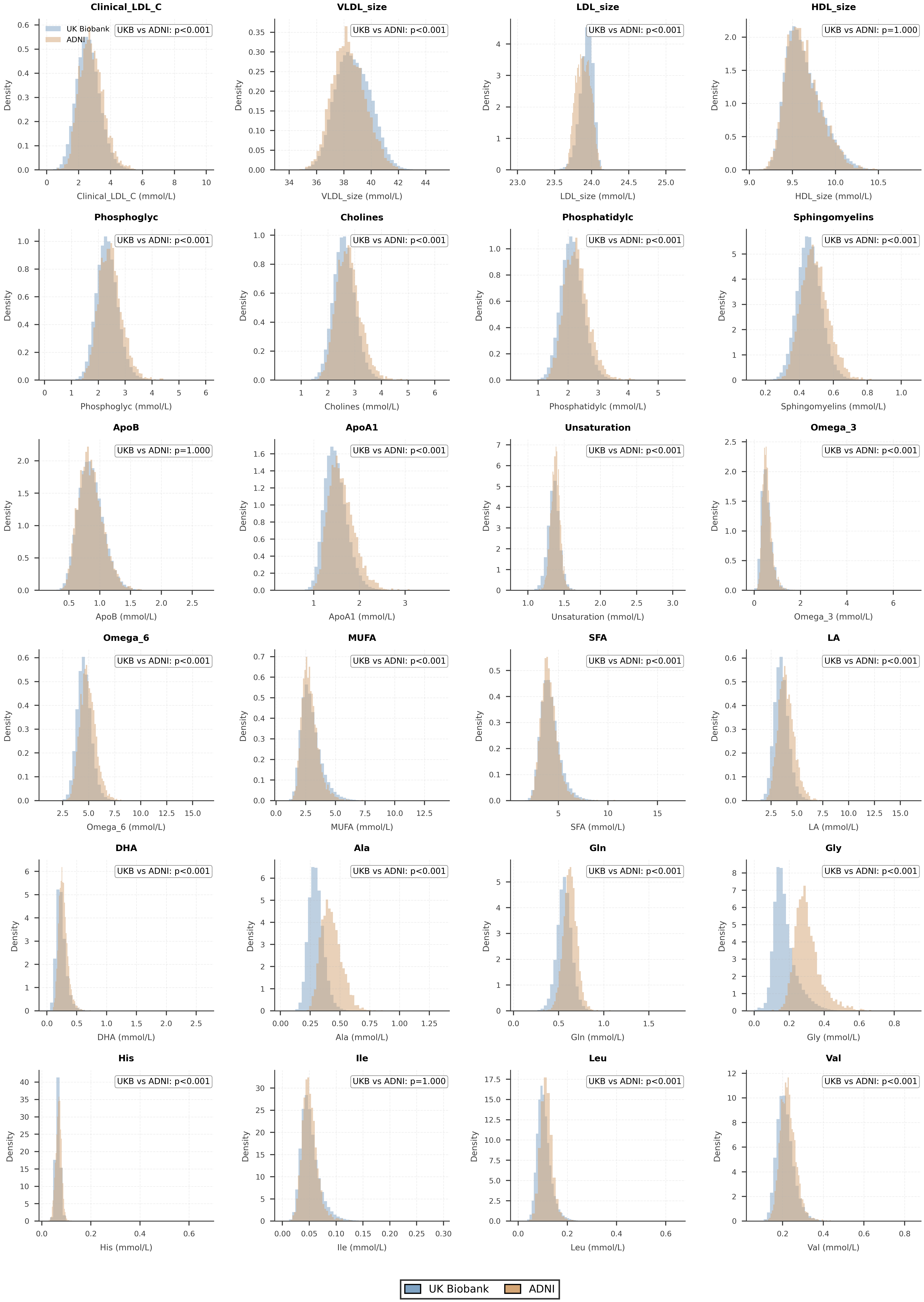

### NMR metabolites · UK Biobank vs ADNI (part 2 of 5)

Distribution of NMR metabolite concentrations across UK Biobank and ADNI cohorts (part 2 of 5). Density-normalized histograms showing 24 non-derived Nightingale NMR metabolites measured in two cohorts: UK Biobank (blue, n=507,783) and ADNI (orange, n=4,851). Each subplot represents one metabolite (metabolites 25–48 of 107) with concentration in mmol/L on the x-axis. Metabolites are shown in UK Biobank panel order. Statistical comparisons were performed using two-sided Wilcoxon rank sum tests, with Bonferroni correction applied across all 107 metabolites. P-values are displayed in the top-right corner of each subplot: p<0.001, p<0.01, and p<0.05 denote statistical significance after correction. Otherwise, exact adjusted p-values are shown.

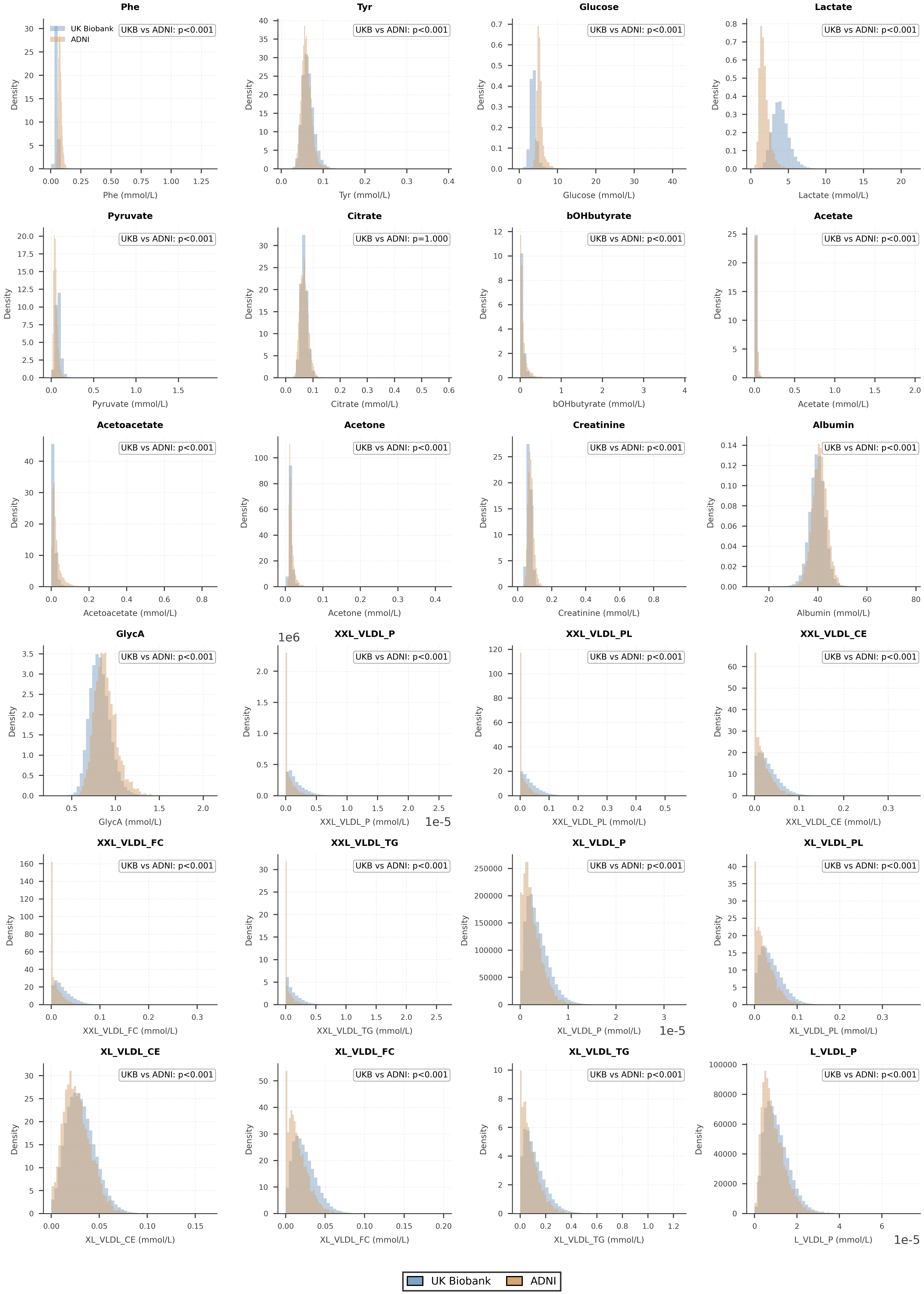

### NMR metabolites · UK Biobank vs ADNI (part 3 of 5)

Distribution of NMR metabolite concentrations across UK Biobank and ADNI cohorts (part 3 of 5). Density-normalized histograms showing 24 non-derived Nightingale NMR metabolites measured in two cohorts: UK Biobank (blue, n=507,783) and ADNI (orange, n=4,851). Each subplot represents one metabolite (metabolites 49–72 of 107) with concentration in mmol/L on the x-axis. Metabolites are shown in UK Biobank panel order. Statistical comparisons were performed using two-sided Wilcoxon rank sum tests, with Bonferroni correction applied across all 107 metabolites. P-values are displayed in the top-right corner of each subplot: p<0.001, p<0.01, and p<0.05 denote statistical significance after correction. Otherwise, exact adjusted p-values are shown.

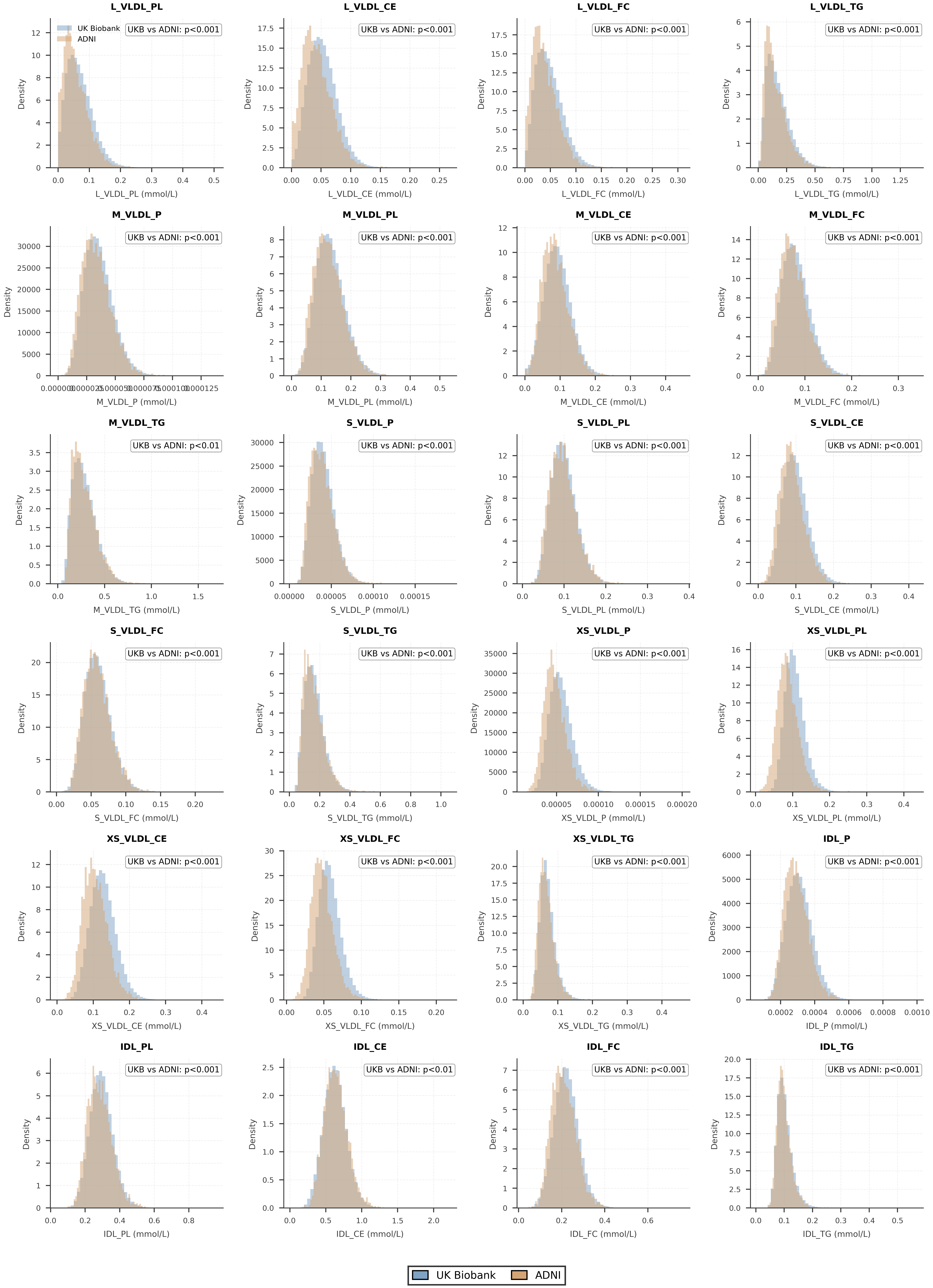

### NMR metabolites · UK Biobank vs ADNI (part 4 of 5)

Distribution of NMR metabolite concentrations across UK Biobank and ADNI cohorts (part 4 of 5). Density-normalized histograms showing 24 non-derived Nightingale NMR metabolites measured in two cohorts: UK Biobank (blue, n=507,783) and ADNI (orange, n=4,851). Each subplot represents one metabolite (metabolites 73–96 of 107) with concentration in mmol/L on the x-axis. Metabolites are shown in UK Biobank panel order. Statistical comparisons were performed using two-sided Wilcoxon rank sum tests, with Bonferroni correction applied across all 107 metabolites. P-values are displayed in the top-right corner of each subplot: p<0.001, p<0.01, and p<0.05 denote statistical significance after correction. Otherwise, exact adjusted p-values are shown.

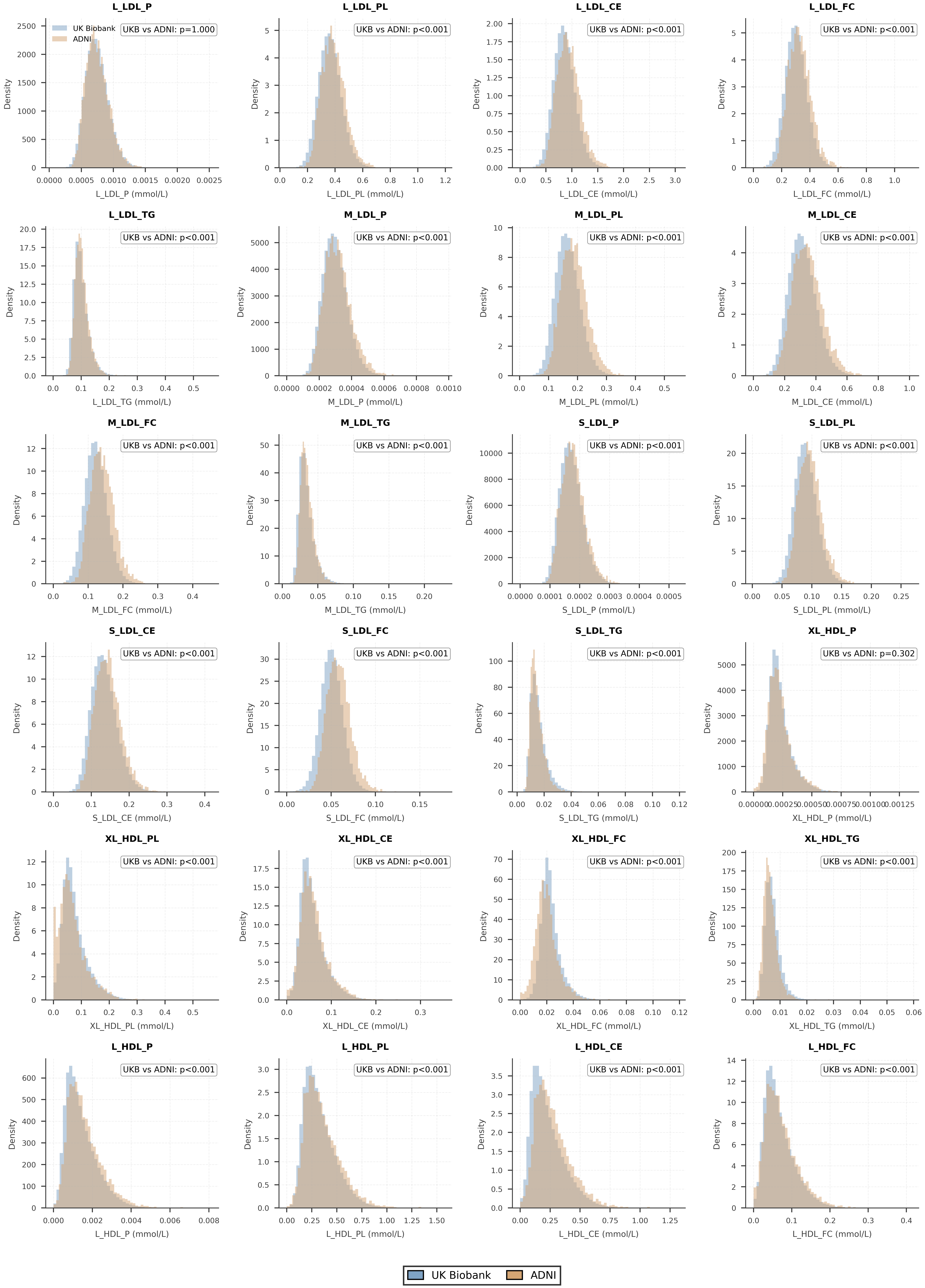

### NMR metabolites · UK Biobank vs ADNI (part 5 of 5)

Distribution of NMR metabolite concentrations across UK Biobank and ADNI cohorts (part 5 of 5). Density-normalized histograms showing 11 non-derived Nightingale NMR metabolites measured in two cohorts: UK Biobank (blue, n=507,783) and ADNI (orange, n=4,851). Each subplot represents one metabolite (metabolites 97–107 of 107) with concentration in mmol/L on the x-axis. Metabolites are shown in UK Biobank panel order. Statistical comparisons were performed using two-sided Wilcoxon rank sum tests, with Bonferroni correction applied across all 107 metabolites. P-values are displayed in the top-right corner of each subplot: p<0.001, p<0.01, and p<0.05 denote statistical significance after correction. Otherwise, exact adjusted p-values are shown.

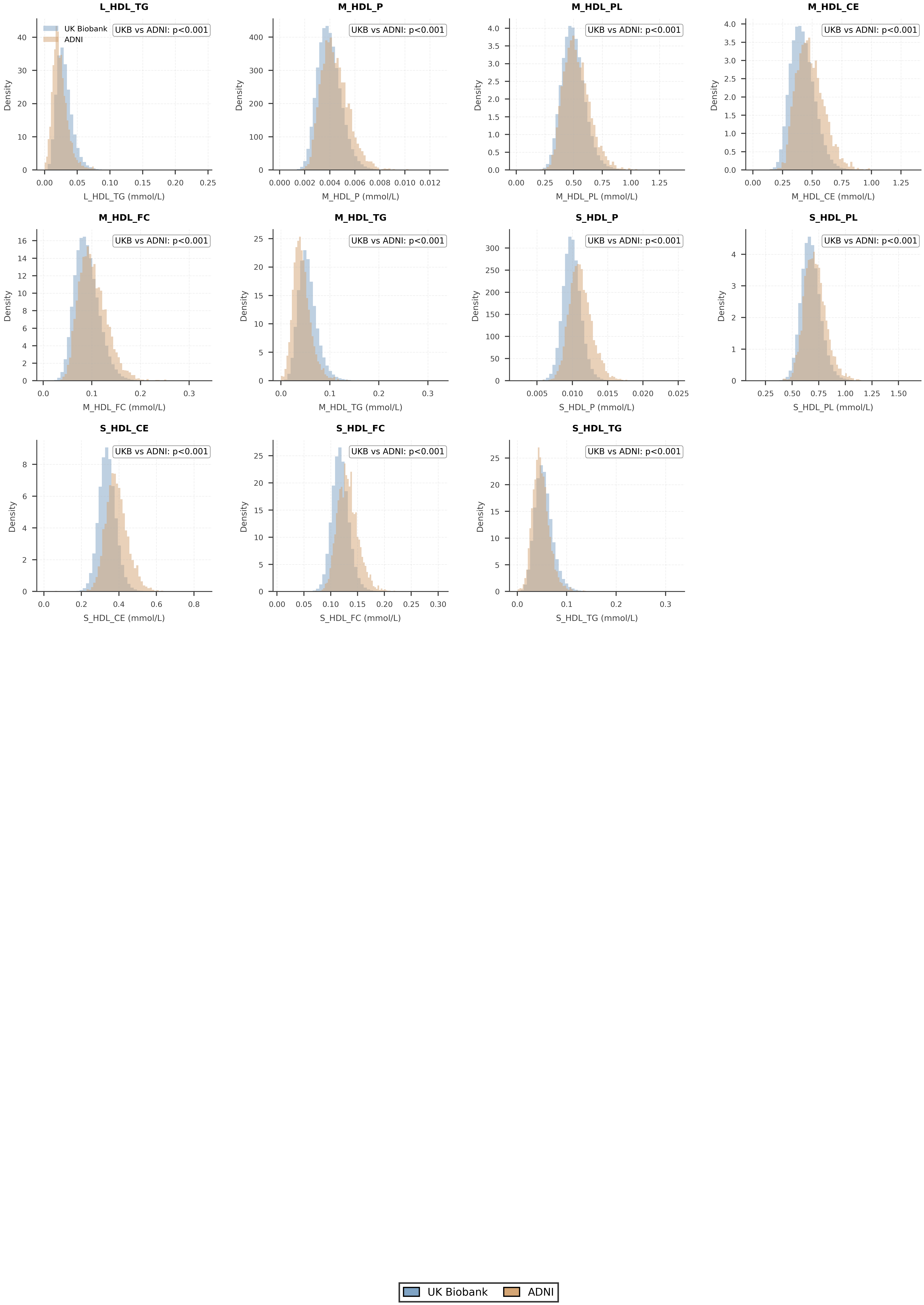

### ADNI NMR vs LC-MS · nine overlapping metabolites

Distribution of overlapping metabolite concentrations measured by NMR and LC-MS within ADNI. Density-normalized histograms showing nine metabolites with direct semantic overlap between Nightingale NMR and Biocrates Q300 LC-MS platforms in ADNI participants with both measurements (n=4,142): NMR (orange) and LC-MS (red, concentrations converted from µM to mmol/L by dividing by 1,000). Each subplot shows one metabolite pair (NMR token / LC-MS name) with concentration in mmol/L on the x-axis. Statistical comparisons between platforms were performed using two-sided Wilcoxon rank sum tests, with Bonferroni correction applied across all nine metabolites. P-values are displayed in the top-right corner of each subplot: p<0.001, p<0.01, and p<0.05 denote statistical significance after correction. Otherwise, exact adjusted p-values are shown.

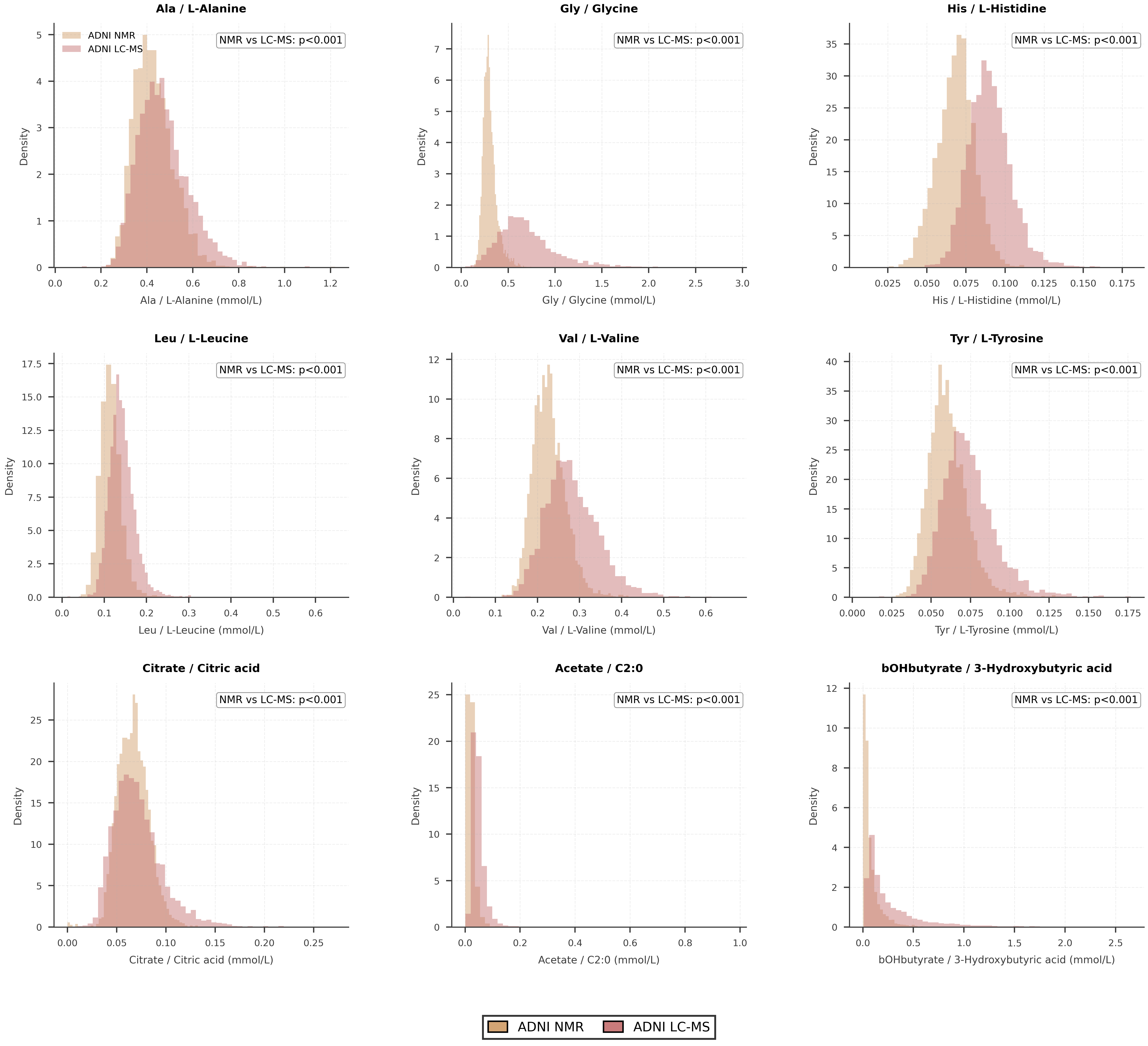
