## Supplementary Figures for "Decoding heterogeneous aging clocks and disease risk stratification using MetAgeFormer"

MetAgeFormer

Biological Aging Analysis

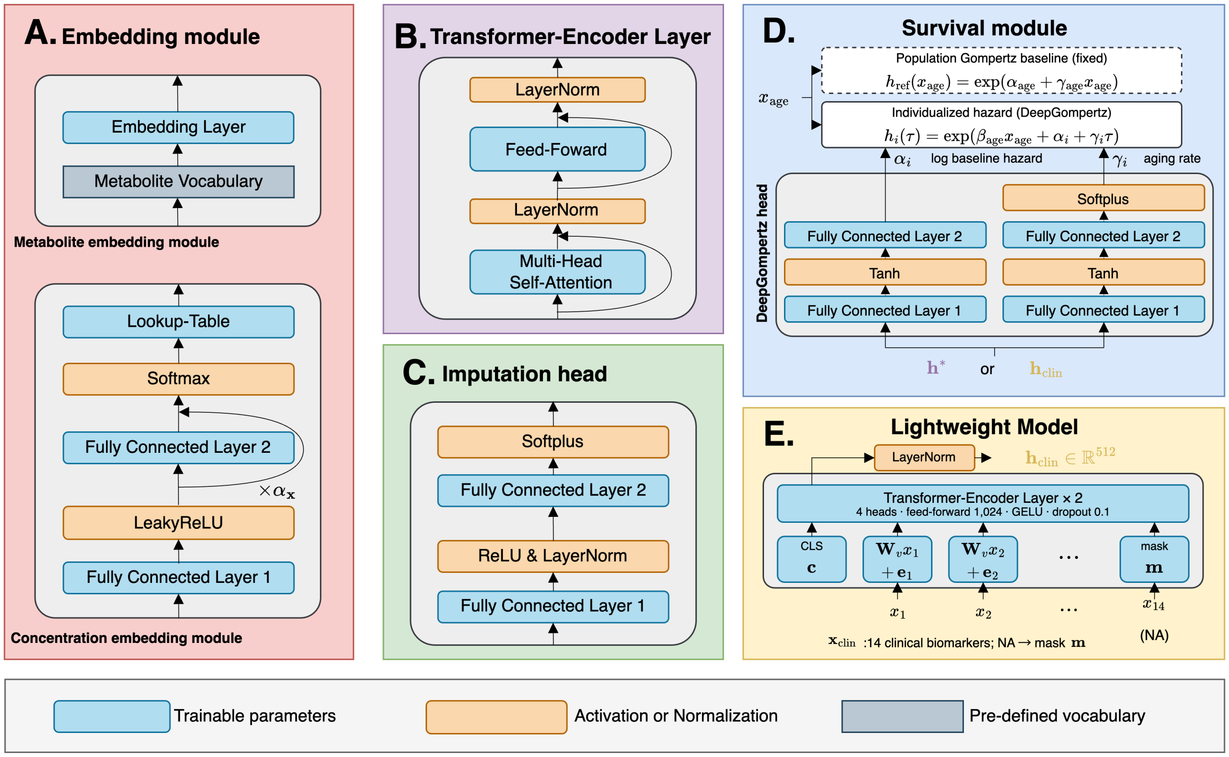


**Supplementary Figure S1.** Module architectures. **(A)** Embedding module. Two sub-modules are designed for encoding metabolite symbols and concentrations, respectively. **(B)** Transformer-encoder layer. The backbone of MetAgeFormer is a stack of six transformer-encoder layers. **(C)** Imputation head. This module is only used during the pre-training stage for self-supervised learning. **(D)** Survival module. This module takes metabolomic embeddings generated from MetAgeFormer or the lightweight model as inputs. Two parallel feed-forward branches of the DeepGompertz head predict the individual log baseline hazard ($\alpha_i$) and the individual Gompertz aging rate ($\gamma_i$), whereas chronological age enters the hazard only through the population-shared coefficient $\beta_{\text{age}}$. The population-level Gompertz baseline is fitted on the training set before fine-tuning and is kept fixed thereafter. **(E)** The lightweight model. Each clinical biomarker value is linearly projected by a shared weight ($\mathbf{W}_v$) and added to a learnable positional embedding ($\mathbf{e}_j$); a missing value is instead represented by a learnable mask embedding ($\mathbf{m}$), which carries no positional embedding and is masked out as an attention key, so that it makes no contribution to the CLS embedding. The missing pattern is therefore conveyed implicitly by which positional embeddings are present. A CLS embedding ($\mathbf{c}$) is prepended to the biomarker tokens and updated by two transformer-encoder layers, and its layer-normalized output serves as the clinical embedding ($\mathbf{h}_{\text{clin}}$).

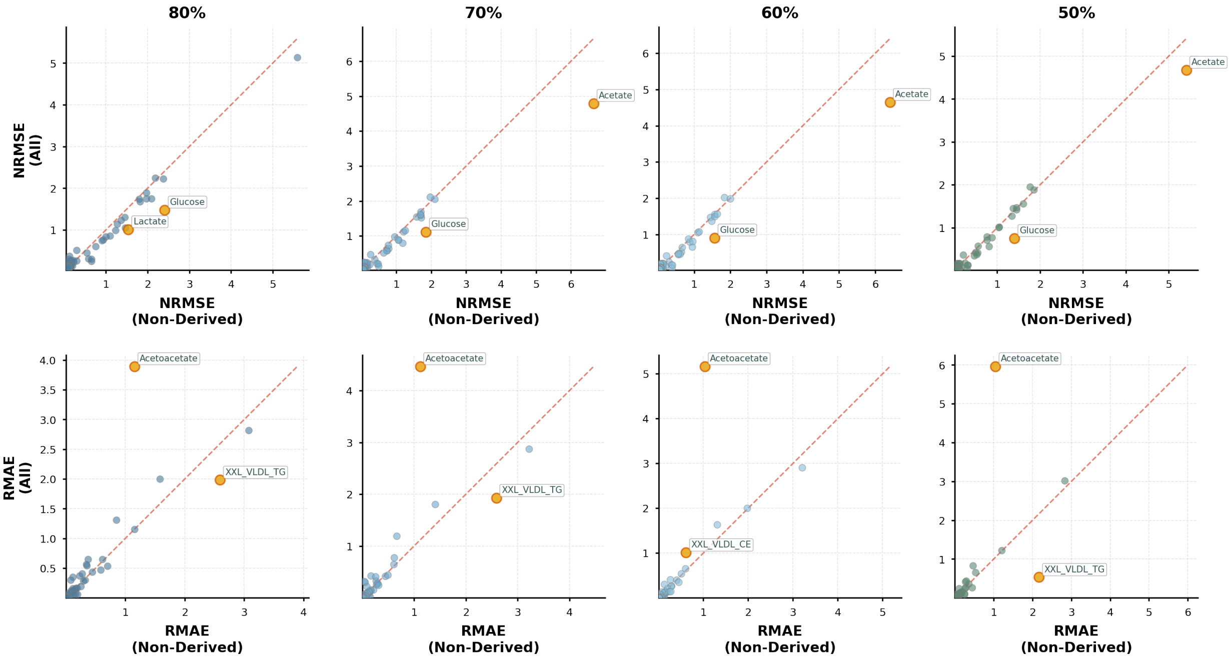


**Supplementary Figure S2.** Imputation performance of models trained on 107 non-derived metabolites versus all 326 metabolite measures. Scatter plots comparing imputation accuracy at varying sparsity levels (80%, 70%, 60%, and 50% data retention). Top row: NRMSE (Normalized Root Mean Square Error), calculated as RMSE divided by the range of observed values (maximum - minimum), implemented using the Permetrics package (`https://permetrics.readthedocs.io/en/latest/pages/regression/NRMSE.html`). Bottom row: RMAE (Relative Mean Absolute Error), calculated as $|\hat{y} - y|/y$ to normalize for scale differences across metabolites. Each point represents one of the 107 non-derived metabolites tested in both models (n=107). Outlier metabolites (>99th percentile distance from diagonal, orange) are annotated with abbreviations. Both models demonstrate comparable imputation accuracy, indicating that training on non-derived metabolites alone captures sufficient information for accurate prediction despite using fewer features.

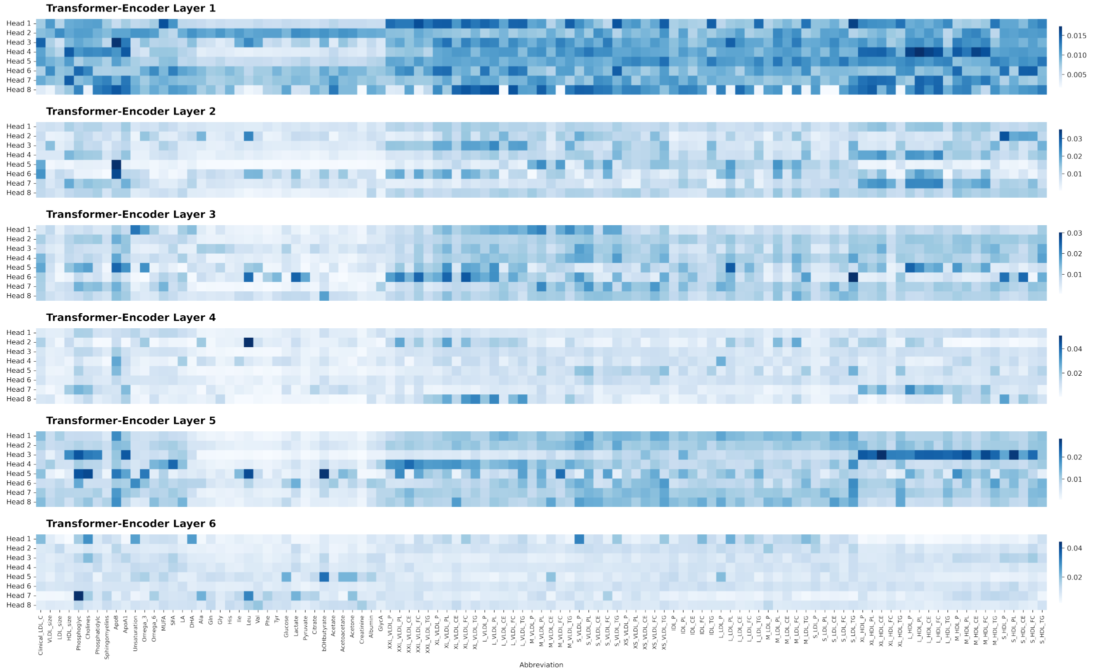


**Supplementary Figure S3.** Average attention scores across heads within each transformer-encoder layer. Attention matrices were extracted from the pre-trained MetAgeFormer on the UKB hold-out test set (n = 73,417) and averaged over samples, yielding 48 matrices (8 heads $\times$ 6 layers). Within each panel, rows correspond to the eight attention heads of that layer and columns to the 107 non-derived metabolites; colour indicates the mean attention each metabolite received from all positions, including itself. Attention was broadly distributed in early layers and became more concentrated on subsets of metabolites in deeper layers. These aggregates are descriptive and are not interpreted as causal biological mechanisms.

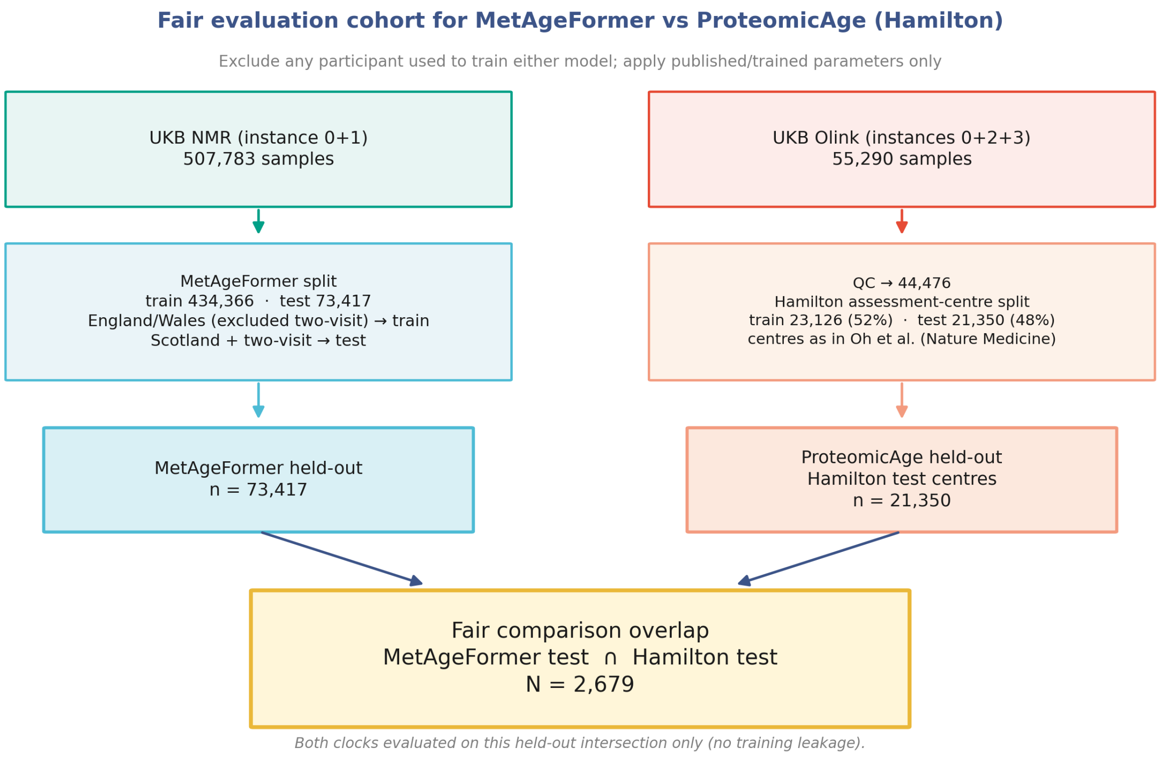


**Supplementary Figure S4.** Schematic illustration of the comparison setting between MetAgeFormer and ProteomicAge (Hamilton).

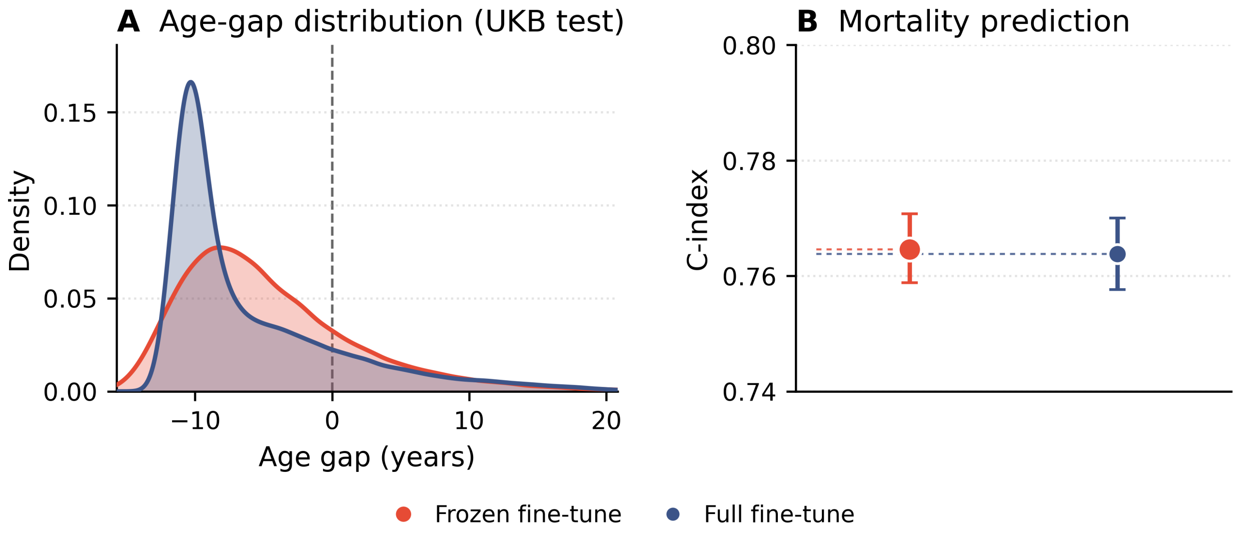


**Supplementary Figure S5.** Ablation of frozen versus full fine-tuning of the survival module. **(A)** Kernel density estimates of ΔAge in the UKB hold-out test set under frozen fine-tuning (default; the pre-trained transformer weights are frozen) and full fine-tuning (all weights unlocked). The dashed vertical line denotes ΔAge = 0. **(B)** Mortality C-index for the two strategies, with 95% bootstrap confidence intervals. The two strategies yielded nearly identical discrimination (0.765 [0.759, 0.771] versus 0.764 [0.758, 0.770], p = 0.36) despite modest shifts in the ΔAge distribution.

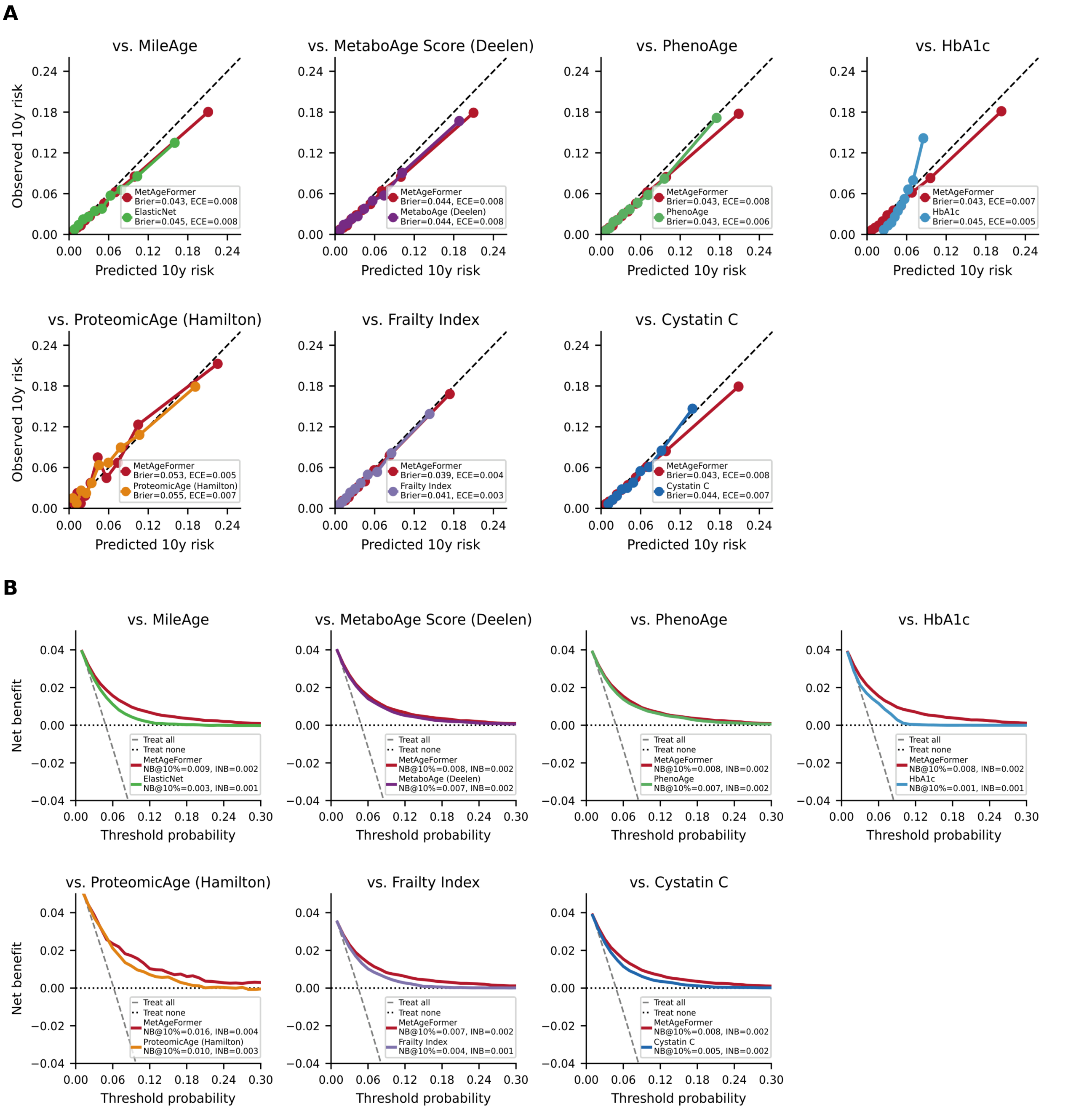


**Supplementary Figure S6.** Calibration and decision-curve analysis for 10-year mortality prediction. **(A)** For each head-to-head comparison, Cox proportional hazards models were fit on the discovery (training) set using age acceleration ($\Delta$age), chronological age, and sex, and 10-year mortality risk was derived from the predicted survival function. Points show quantile-binned calibration curves; the dashed diagonal denotes perfect calibration. Legend entries report the Brier score and expected calibration error (ECE). **(B)** Decision curve analysis (DCA) of the same models and cohort subsets. Net benefit is plotted against the threshold probability of treating an individual as high risk; dashed grey and dotted black lines indicate the "treat all" and "treat none" strategies, respectively. Legend entries report net benefit at a 10% threshold (NB@10%) and integrated net benefit (INB). Each column compares MetAgeFormer (red) with one benchmark: MileAge (ElasticNet), MetaboAge (Deelen), PhenoAge, glycated haemoglobin (HbA1c), ProteomicAge (Hamilton), Frailty Index, or Cystatin C. Evaluation sample sizes depend on biomarker availability (n = 73,417 for MileAge and PhenoAge; 68,211 for MetaboAge; 69,030 for Cystatin C; 65,091 for HbA1c; 53,843 for Frailty Index; 2,679 for ProteomicAge (Hamilton)).

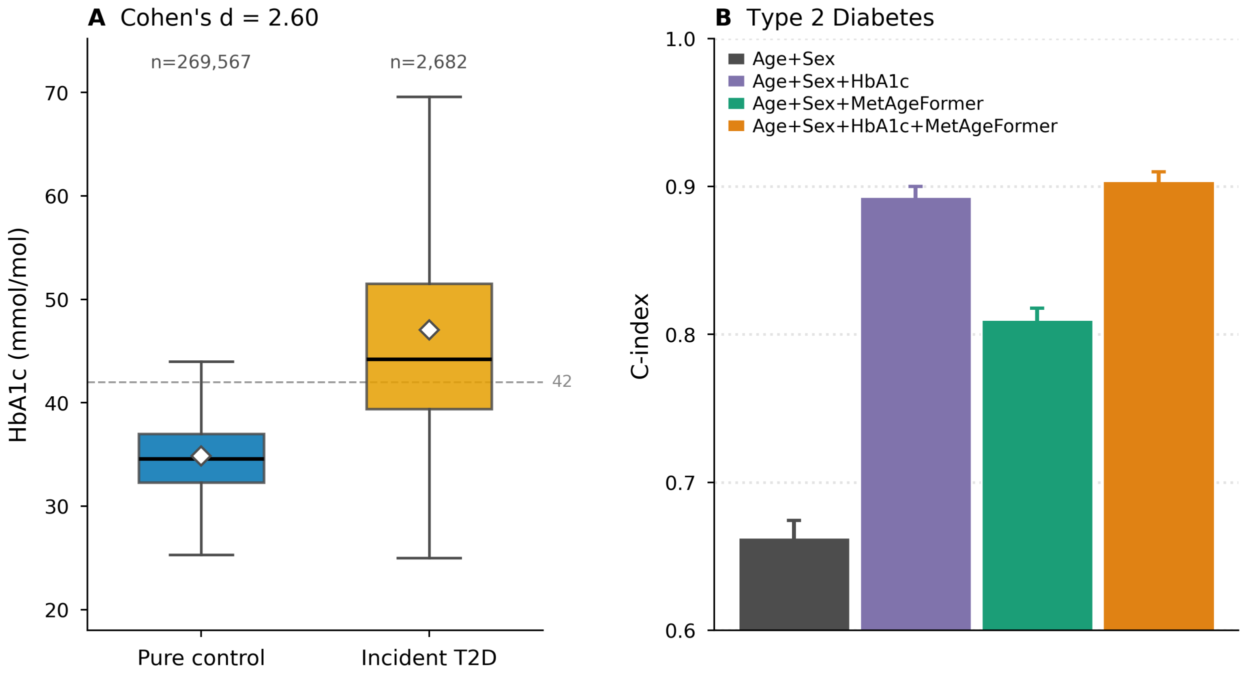


**Supplementary Figure S7.** HbA1c in incident type 2 diabetes and the incremental value of MetAgeFormer. **(A)** Distribution of baseline HbA1c in participants who developed incident type 2 diabetes (n = 2,682) versus pure controls (n = 269,567), defined as participants free of any of the 12 disease endpoints at baseline and during the subsequent 10 years (**Methods**). Boxes show the interquartile range, black lines the medians, and white diamonds the means; the dashed horizontal line marks the diagnostic threshold of 42 mmol/mol. Baseline HbA1c was substantially higher in incident cases (Cohen's d = 2.60). **(B)** C-index for incident type 2 diabetes from Cox models including chronological age and sex, with HbA1c, MetAgeFormer $\Delta$Age, or both added. Bars show the mean C-index and error bars the standard deviation across 1,000 bootstrap iterations. Adding MetAgeFormer $\Delta$Age to HbA1c further improved discrimination (0.904 versus 0.893 for HbA1c alone).

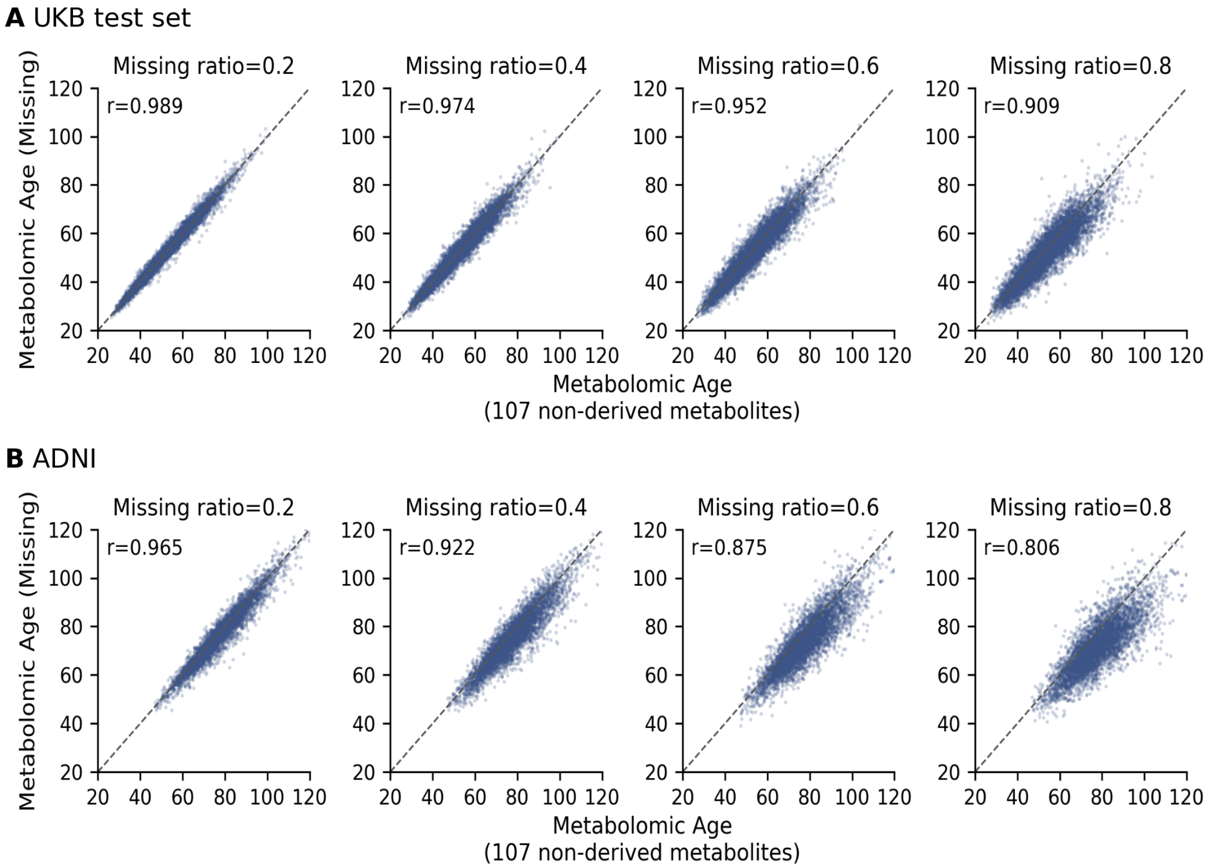


**Supplementary Figure S8.** Concordance of predicted metabolomic age under simulated missing metabolites. Metabolites were randomly masked at ratios of 0.2, 0.4, 0.6, and 0.8 (columns), and the resulting metabolomic age (y-axis) was compared with that predicted from the full panel of 107 non-derived metabolites (x-axis). **(A)** UKB hold-out test set (n = 73,417). **(B)** ADNI (n = 4,851 samples). Each point is one sample, the dashed line is the identity line, and the inset reports the Pearson correlation.

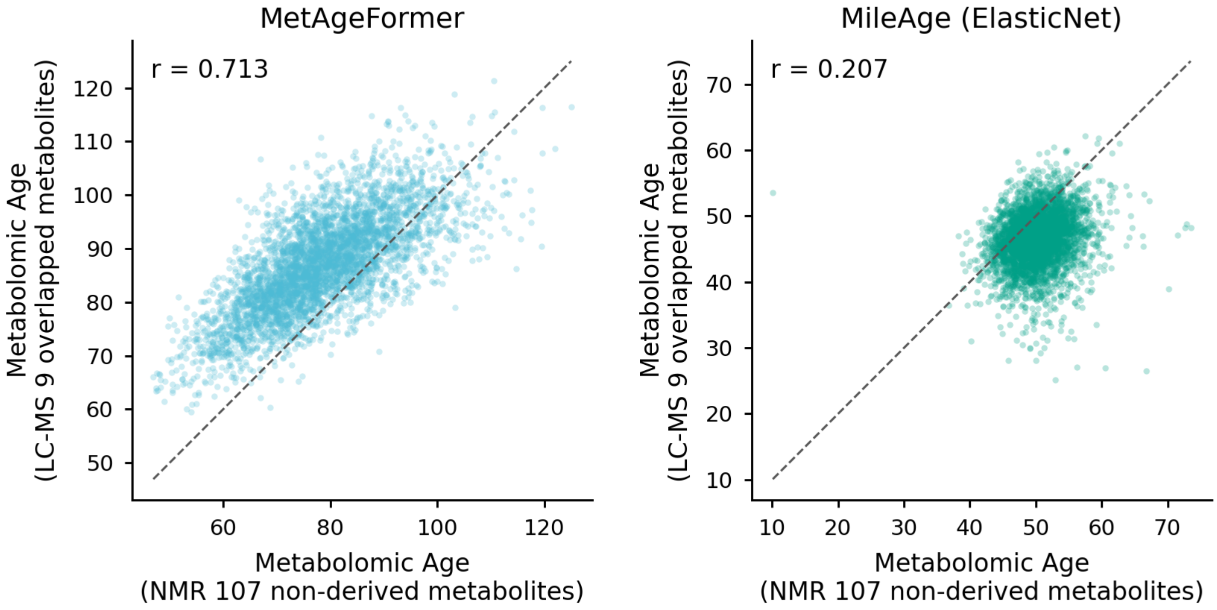


**Supplementary Figure S9.** Cross-platform evaluation of MetAgeFormer in metabolomic age estimation using nine LC–MS–NMR overlapping metabolites in ADNI.

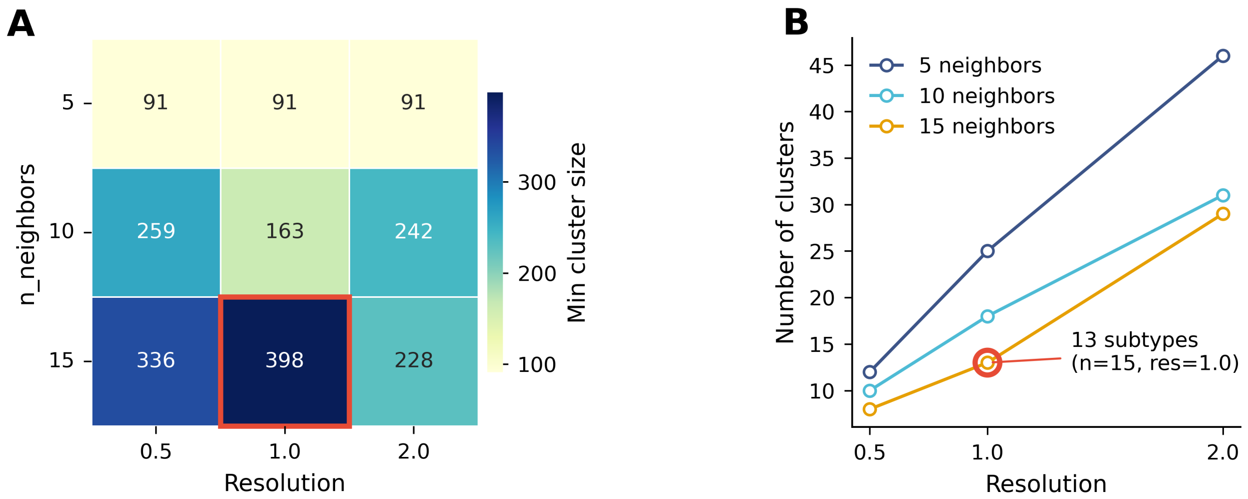


**Supplementary Figure S10.** Grid search of Leiden clustering parameters. Leiden clustering was applied to metabolomic embeddings of the UKB training set over a grid of the number of neighbours (5, 10, 15) and resolution (0.5, 1.0, 2.0), using cosine distance. **(A)** Size of the smallest resulting cluster for each parameter combination; the red box marks the selected setting (n_neighbors = 15, resolution = 1.0), which maximised the minimum cluster size (n = 398). **(B)** Number of clusters as a function of resolution for each neighbourhood size. The selected setting is circled and yielded 13 metabolic subtypes.

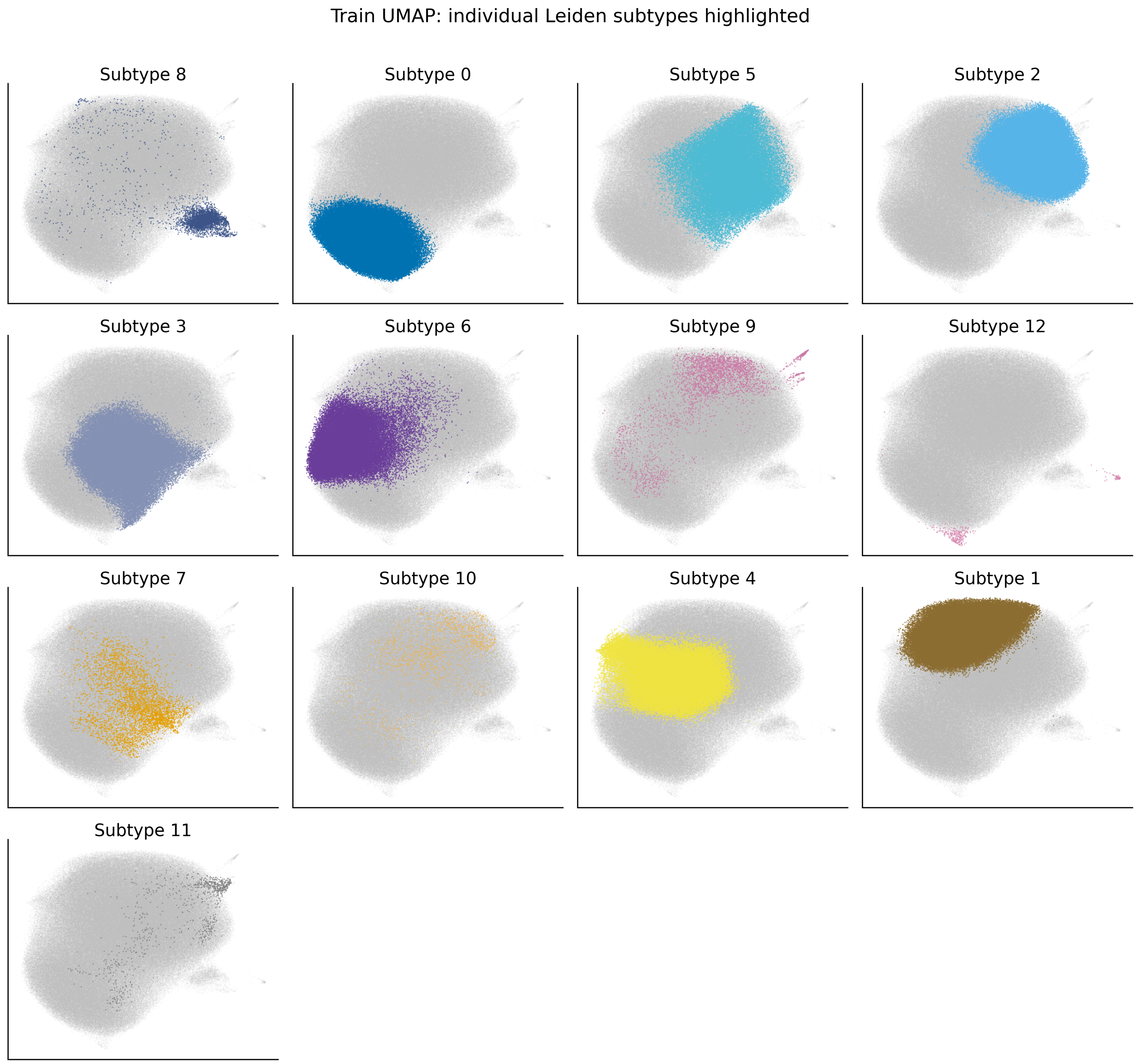


**Supplementary Figure S11.** Visualization of each metabolic subtype. UMAP projection of metabolomic embeddings from the UKB training set, with each of the 13 Leiden subtypes highlighted individually in colour; all remaining participants are shown in grey.

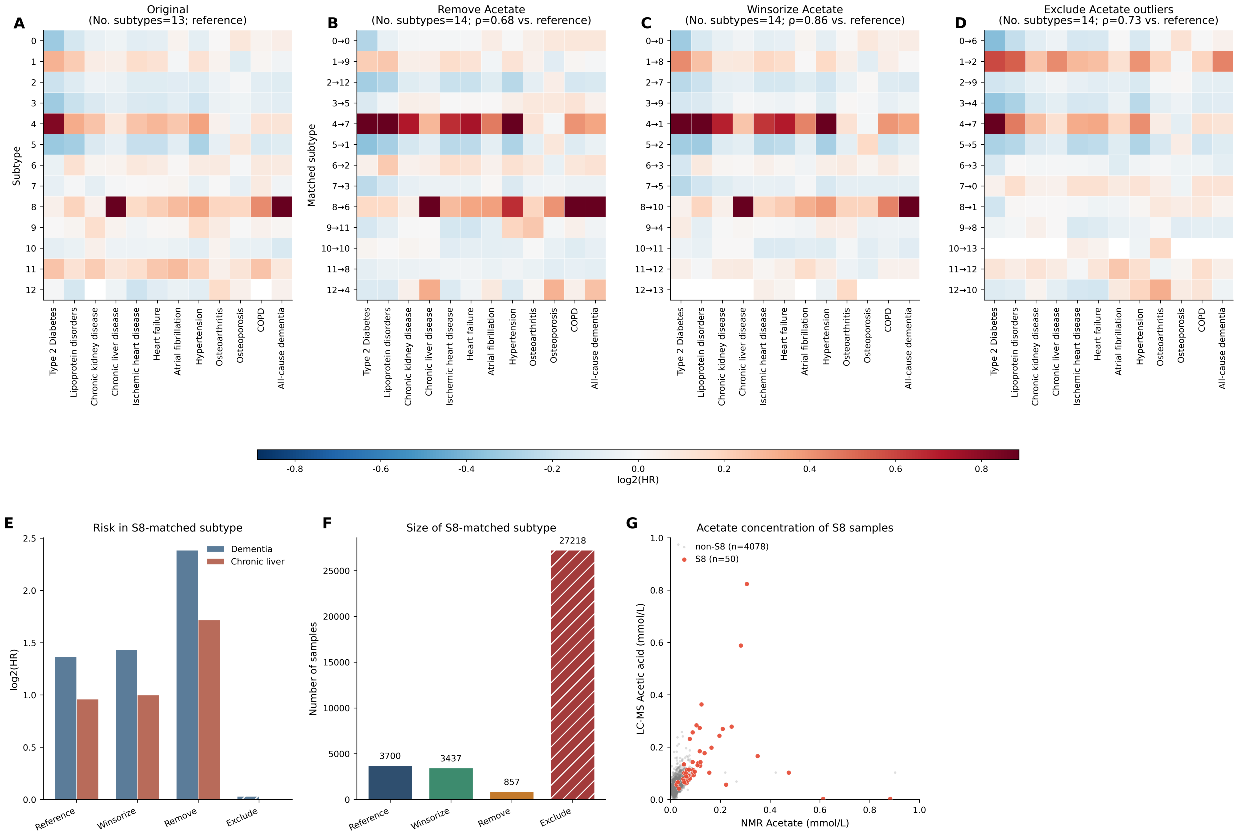


**Supplementary Figure S12.** Acetate sensitivity. Acetate ablation leaves disease-risk structure largely recoverable **(A-C)**, except when extreme Acetate carriers are removed **(D)**. **(E-F)** Disease risk and size in the subtype 8 (S8)–matched subtypes. **(G)** Cross-platform acetate concordance in the ADNI cohort with S8 highlighted (n=4,128 samples with non-missing Acetate on both the Nightingale NMR and targeted LC–MS platforms).

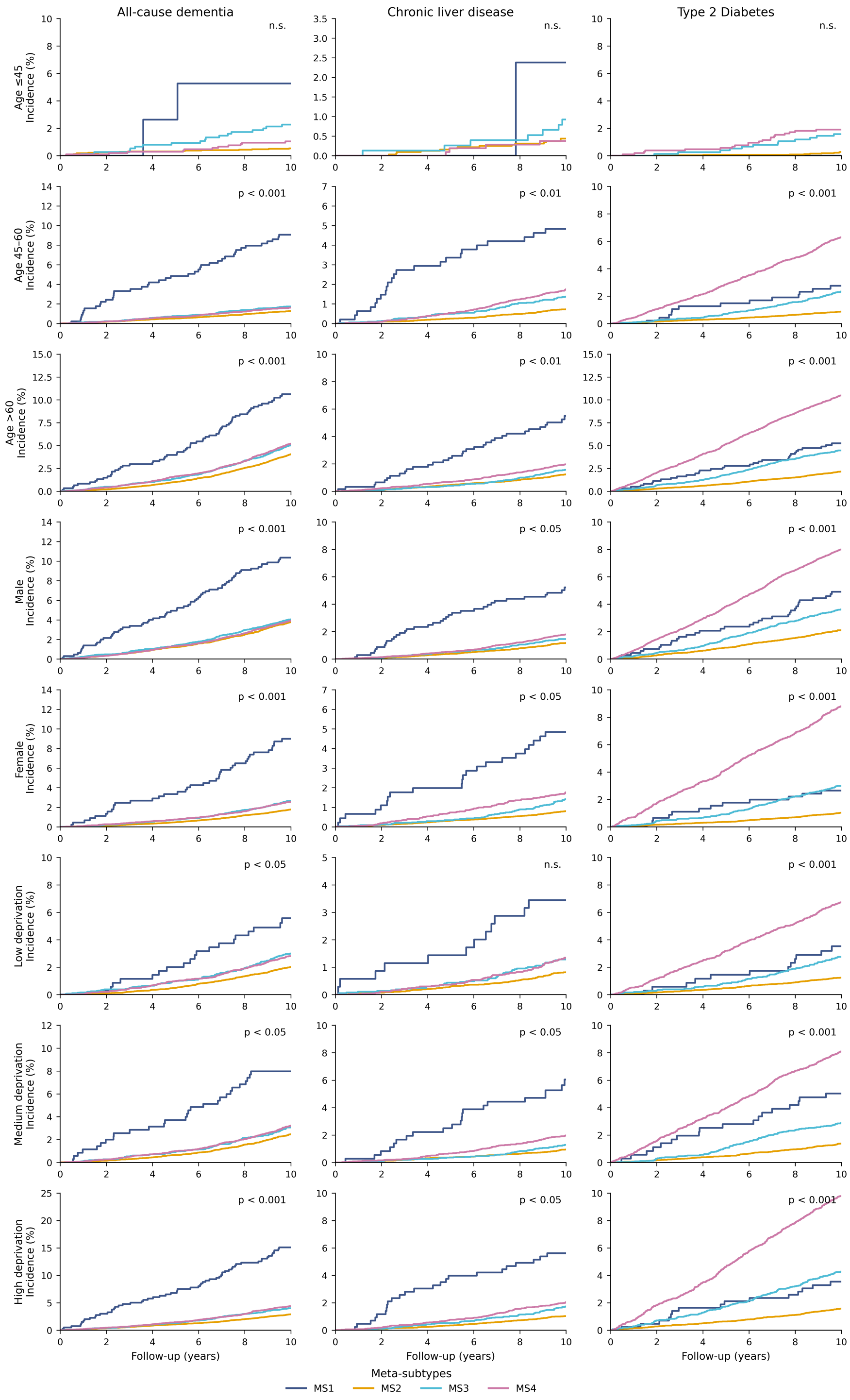


**Supplementary Figure S13.** Stratification analysis of subtype demographic sensitivity. Cumulative incidence curves for the age-related diseases within the UKB hold-out test set, partitioned by age intervals, sex, and Townsend Deprivation Index, respectively. Statistical significance across strata was determined via the log-rank test.

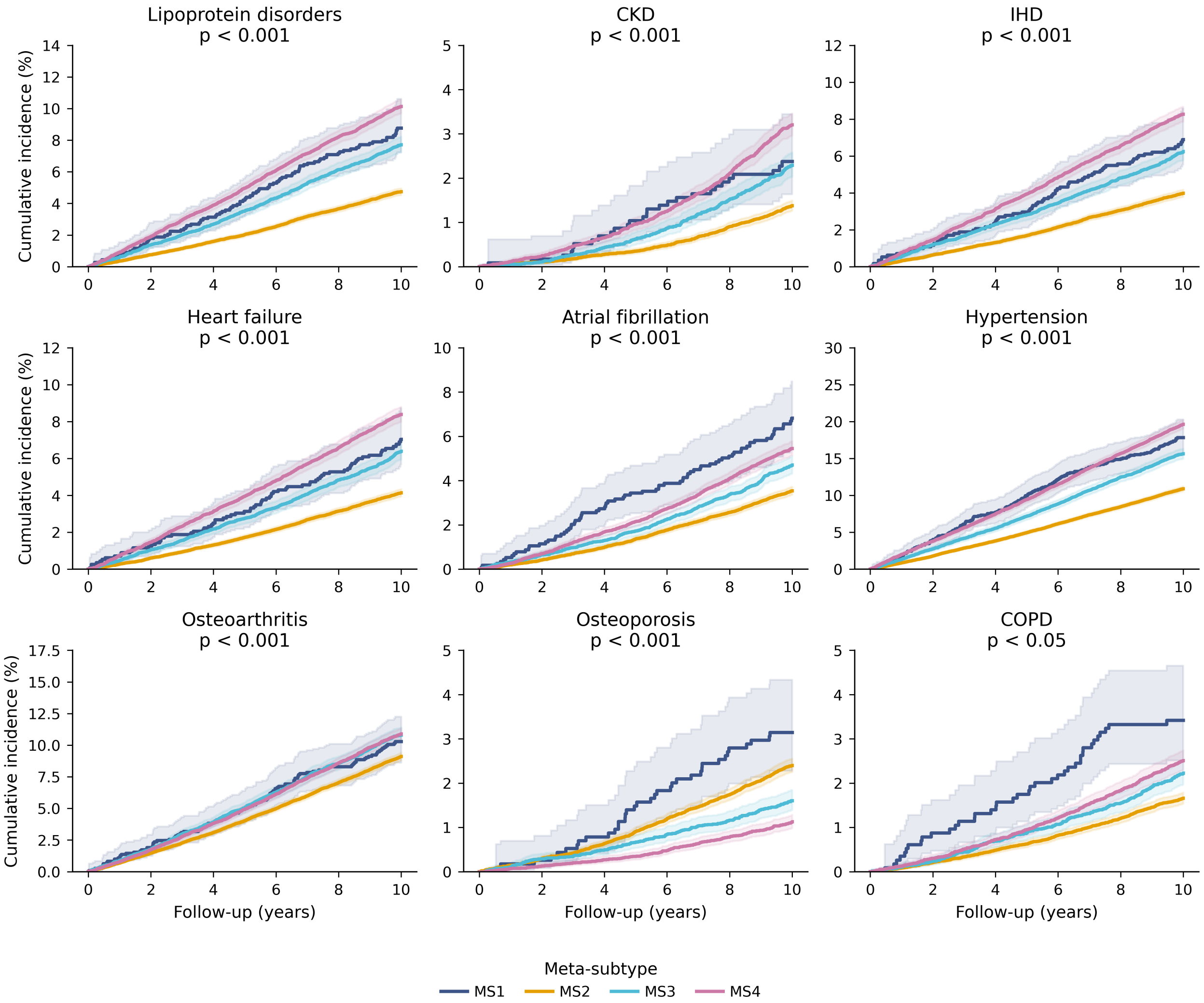


**Supplementary Figure S14.** Cumulative incidence for age-related diseases in the UKB hold-out test set. P-values were calculated using the log-rank test.

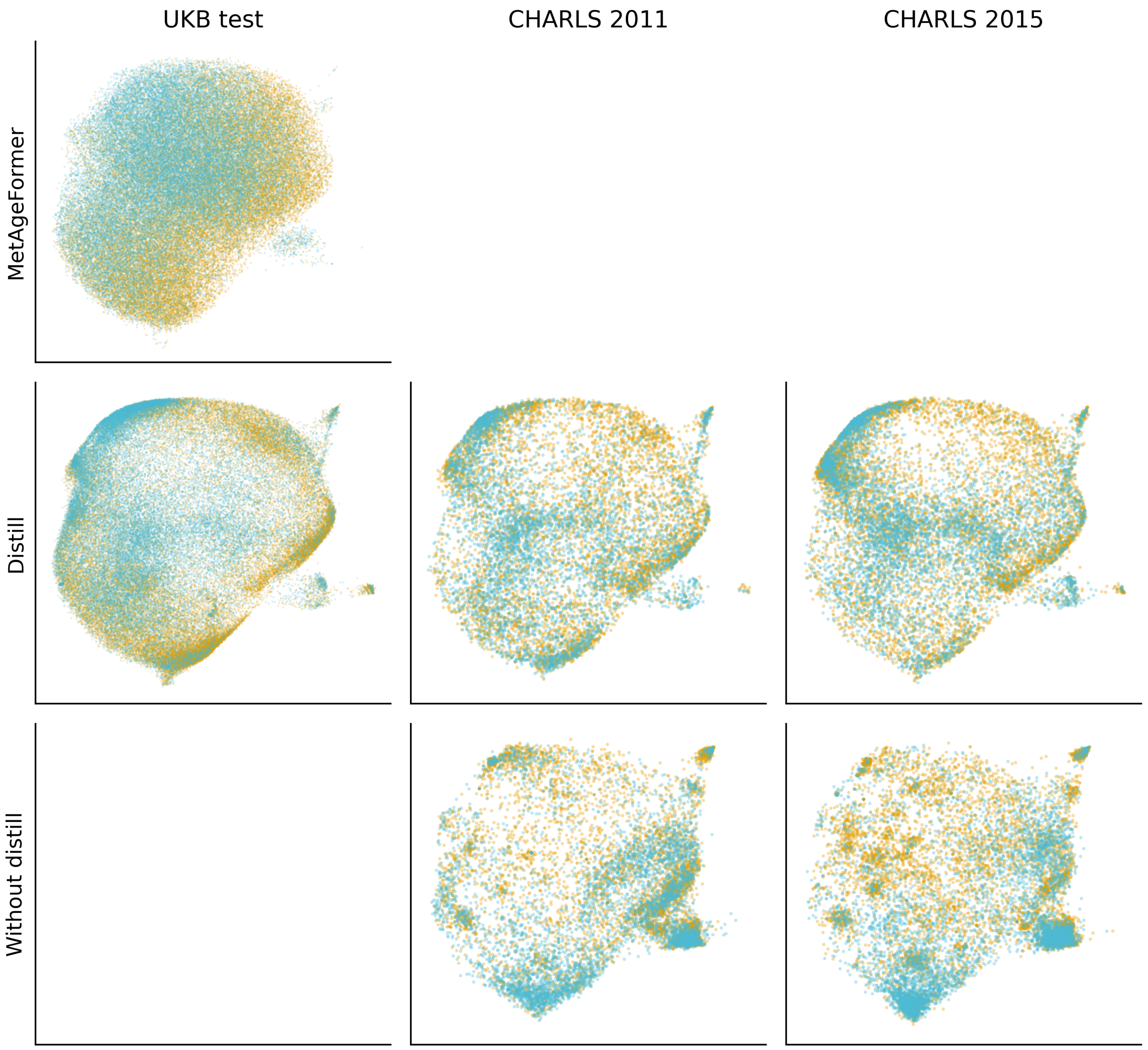


**Supplementary Figure S15.** Distillation ablation of hidden representation space. Two-dimensional UMAP projections of hidden representations colored by sex (female, orange; male, blue). MetAgeFormer row: teacher model embeddings on the UKB test set. Distill row: contrastive-distillation lightweight student (14 routine blood biomarkers) on the UKB test set and CHARLS 2011/2015 datasets. Without distill row: ablation student trained without distillation on CHARLS. Distillation preserved sex-structured geometry aligned with the teacher space on external CHARLS cohorts. The ablation model without distillation showed fragmented, cohort-specific clustering.

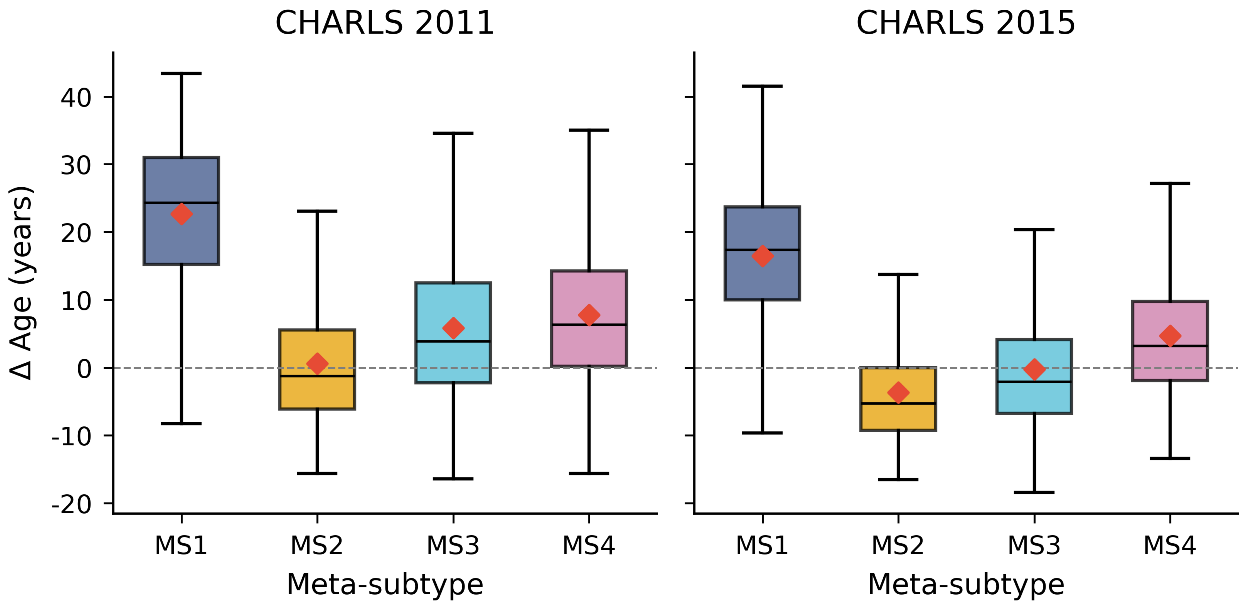


**Supplementary Figure S16.** ΔAge by meta-subtype in CHARLS. Distribution of ΔAge across four meta-subtypes in CHARLS at the 2011 (left) and 2015 (right) waves. Box plots show the interquartile range; horizontal black lines and red diamonds indicate medians and means, respectively. The dashed horizontal line denotes ΔAge = 0. MS1 showed the highest mean ΔAge in both waves (23 and 16 years), whereas MS2 remained near or below zero.

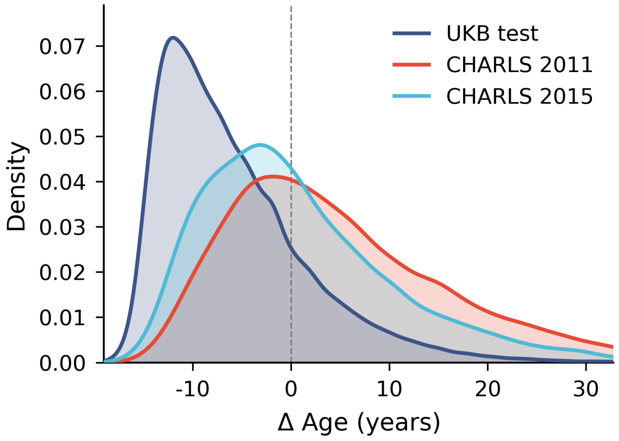


**Supplementary Figure S17.** ΔAge distribution in UK Biobank and CHARLS. Kernel density estimates of ΔAge from the distilled lightweight model in the UKB test set (n = 73,417) and CHARLS 2011 (n = 11,823) and 2015 (n = 13,257) cohorts. The UKB test distribution was centred below zero (mean -5.9 years), whereas CHARLS distributions were shifted toward positive ΔAge (2011 mean +4.6 years; 2015 mean +0.7 years). The vertical dashed line denotes ΔAge = 0.

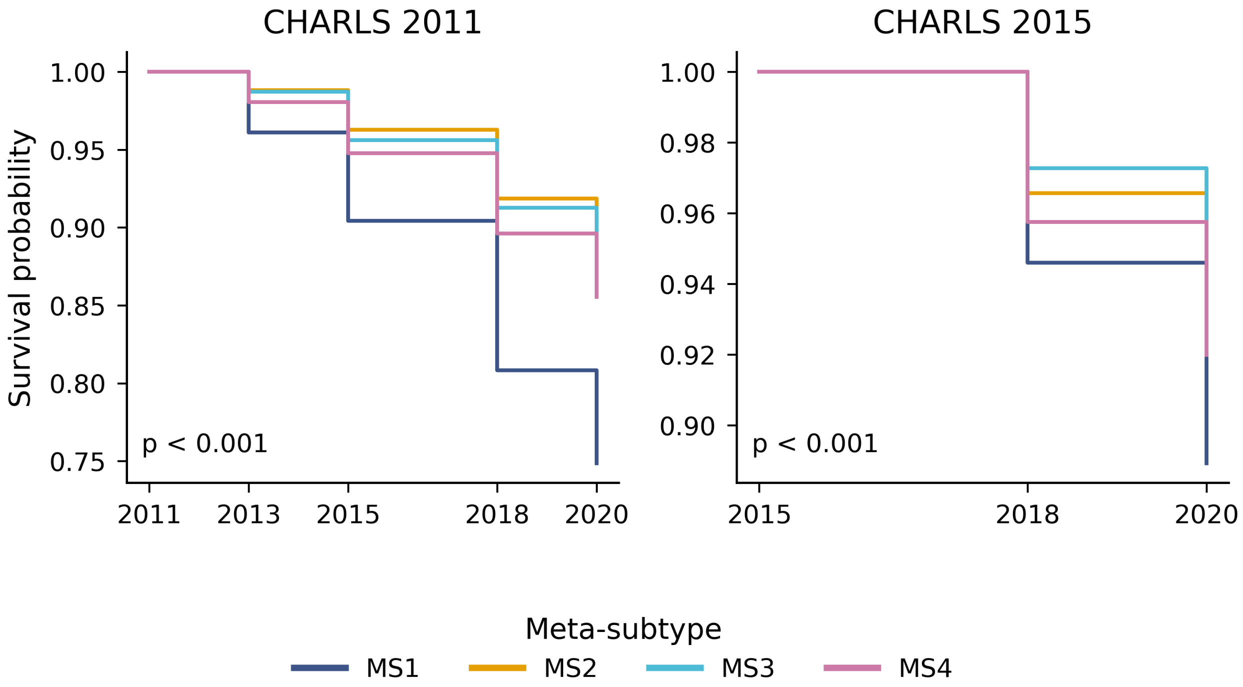


**Supplementary Figure S18.** Survival curves by meta-subtypes in CHARLS. Kaplan–Meier survival curves for mortality stratified by meta-subtypes (MS1–MS4) in CHARLS 2011 (left) and 2015 (right). Meta-subtypes were assigned by the UKB-trained subtype classifier transferred to distilled lightweight embeddings. Multivariate log-rank tests indicated significant differences across subtypes in both waves ($p < 0.001$). MS1 consistently showed the lowest survival probability over follow-up.

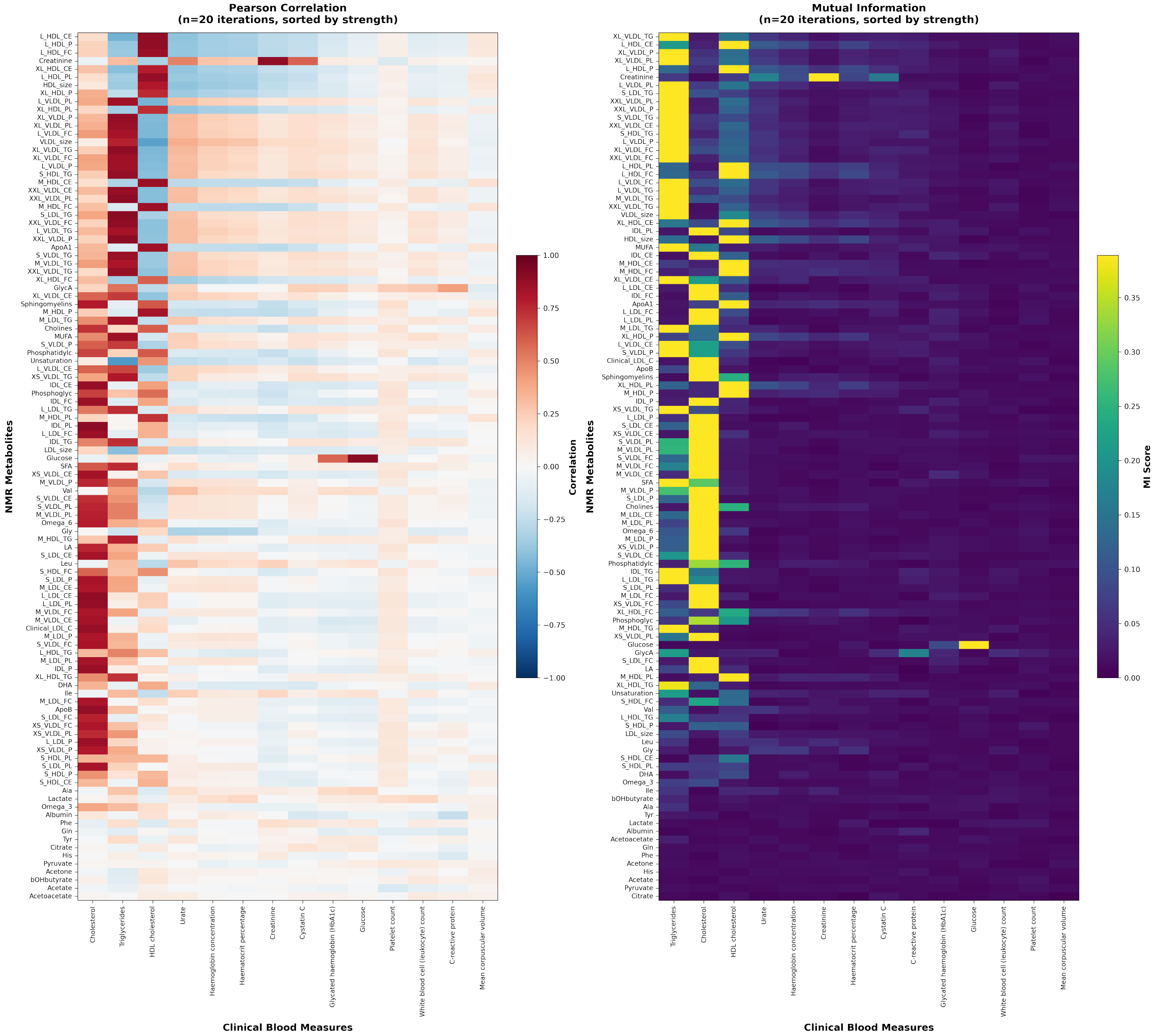


**Supplementary Figure S19.** Correlation and mutual information (MI) between clinical blood biomarkers and NMR-derived metabolites. **(Left panel)** Pearson correlation coefficients between each clinical blood measure (x-axis) and NMR metabolite (y-axis). **(Right panel)** MI scores quantifying non-linear associations between the same feature pairs. Both measures were computed using iterative subsampling (n=20 iterations, 1,000 randomly selected samples per iteration) to ensure robust estimates. Features are sorted by mean association strength (strongest to weakest) along both axes to reveal hierarchical patterns.

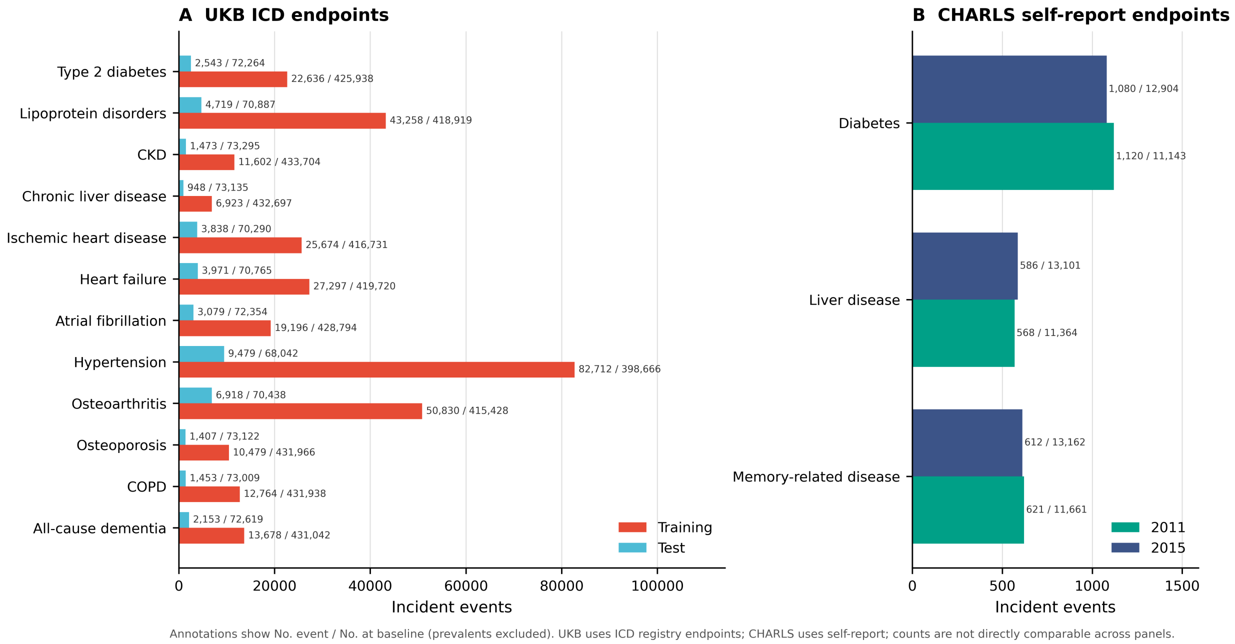


**Supplementary Figure S20.** Incident disease events across cohorts. Bars show the number of incident events for each disease endpoint, with prevalent cases excluded. Annotations give the number of incident events and the number of participants at risk at baseline (events / at risk). **(A)** UK Biobank, with ICD-10-defined endpoints (**Methods**) shown separately for the training set and the hold-out test set. **(B)** CHARLS, with self-reported endpoints shown separately for the 2011 and 2015 waves. Because UKB uses hospital ICD registry records whereas CHARLS uses self-reported physician diagnoses, event counts are not directly comparable between panels.

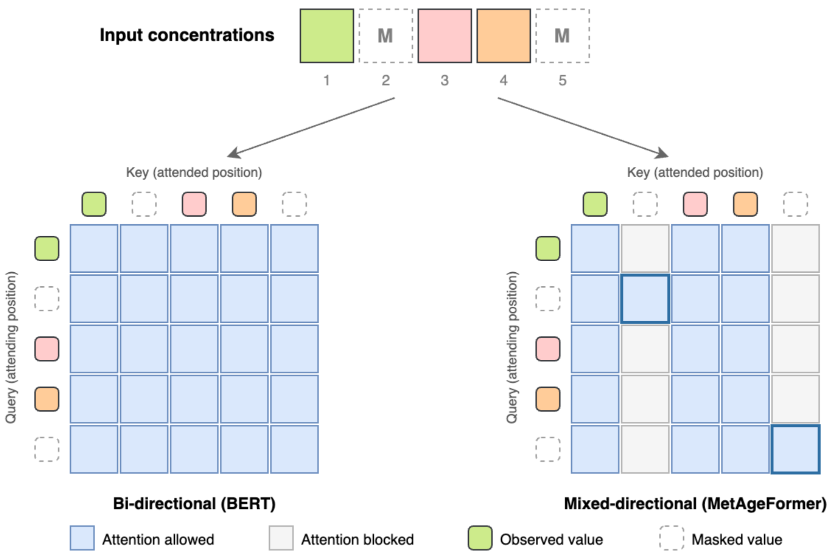


**Supplementary Figure S21.** Mixed-directional attention mask used during masked concentration imputation. Rows denote query positions and columns denote key positions; shaded cells indicate permitted attention. Under the bi-directional mask (left, as in BERT) every position attends to every other position. Under the mixed-directional mask (right), the columns of masked positions are blocked, so masked concentrations are never attended to by other metabolites; each masked position instead attends to the unmasked concentrations and to itself only (bold-outlined diagonal cells). Positions 2 and 5 are masked in this illustration.
