## Supplementary Methods for "Decoding heterogeneous aging clocks and disease risk stratification using MetAgeFormer"

**1. Acetate sensitivity analyses**

**1.1 Overview**

In the main text, we assessed whether the extreme acetate elevation in subtype 8 and its associated disease-risk signals were sensitive to acetate perturbation. Perturbed subtypes that were aligned to a baseline reference subtype on the basis of disease-risk profiles are described in the Results as matched clusters (for example, subtype 8–matched clusters after acetate removal or winsorization). The procedures below provide full methodological details for these analyses, including the assignment algorithm used to define matched clusters.

**1.2 Acetate perturbations**

Analyses were performed on the combined UK Biobank training and validation sets (n = 434,366). Using the fixed pretrained MetAgeFormer checkpoint (Methods), sample-level CLS embeddings were regenerated under three acetate perturbations applied before inference: (i) remove acetate—Acetate was set to missing and masked during embedding extraction; (ii) winsorize acetate—Acetate values were clipped to the training-set 1st–99th percentiles; and (iii) exclude outliers—participants with acetate |z| > 5 relative to the training-set mean and standard deviation were removed (n = 432,299 after exclusion; 2,067 participants removed).

**1.3 Clustering and disease-risk modelling**

For each perturbation, Leiden clustering was applied to the perturbed embeddings using the selected parameters described in the main Methods (n_neighbors = 15, resolution = 1.0, cosine distance). Subtype-versus-rest Cox proportional hazards models adjusted for chronological age, sex, and BMI, matching the specification used for the baseline subtypes in the main analysis, were then fitted for the same 12 ICD disease groups, yielding subtype-specific log hazard ratio (log-HR) matrices. The locked baseline partition (13 subtypes) served as the reference disease-risk structure.

**1.4 Definition of matched clusters**

Because perturbed analyses can alter cluster number and labelling, subtype log-HR profiles were aligned between the baseline reference and each perturbed analysis before comparing disease-risk structures. For each baseline subtype i and perturbed subtype j, we computed the Pearson correlation rij between their log-HR vectors across shared diseases (missing entries set to zero before correlation) and defined an assignment cost 1 − rij. Optimal one-to-one assignment between baseline and perturbed subtypes was obtained using the Hungarian algorithm (also known as the Kuhn–Munkres assignment algorithm)¹, implemented in SciPy (scipy.optimize.linear_sum_assignment)². In the main text, the perturbed subtype assigned to a given baseline subtype under this procedure is referred to as its matched cluster; the corresponding profile correlation (1 minus the assignment cost) quantifies how closely the matched cluster recapitulates the reference subtype’s disease-risk profile.

**1.5 Concordance metrics**

Overall concordance between the baseline and perturbed disease-risk structures was quantified as the Spearman correlation of matched log-HR cells after reordering perturbed subtypes according to the assignment above. Subtype 8 was evaluated by identifying the perturbed subtype assigned to baseline subtype 8 and reporting its profile correlation, cluster size, and log-HRs for all-cause dementia and chronic liver disease. Perturbed communities with low profile correspondence to baseline subtype 8 were not interpreted as recovery of subtype 8 in the main text (for example, after exclusion of extreme acetate samples).

**1.6 ADNI orthogonal acetate validation**

In the Alzheimer’s Disease Neuroimaging Initiative (ADNI) cohort, Nightingale NMR Acetate was compared with targeted LC-MS acetic acid (Q300 C2:0; µM converted to mmol/L) in overlapping samples with non-missing values on both platforms (n = 4,128 of the 4,142 samples with paired NMR and LC-MS profiles; 14 samples lacked an NMR Acetate measurement and were excluded from this comparison. All 14 were assigned to subtype 8 and had non-missing Q300 acetic acid values). UK Biobank-trained subtype labels were transferred to ADNI, and acetate levels were compared between subtype 8 and non–subtype 8 on both platforms.

**2. Cognitive scores for word recall and recognition from the CHARLS cohort**

**2.1 Word recall**

The number of correctly recalled words was utilized as the metric for immediate recall (maximum: 20) and delayed recall (maximum: 10).

**2.2 Word recognition and d-prime**

Recognition performance was quantified using the signal detection theory metric d-prime, which accounts for both sensitivity and response bias. Recognition responses were classified according to the following confusion matrix: True Positives (TP) represented correctly identified target words, reflecting the core indicator of recognition memory ability. True Negatives (TN) represented correctly identified novel words, reflecting the ability to discriminate new stimuli. False Positives (FP) indicated a liberal or impulsive response tendency. False Negatives (FN) represented failures in memory retrieval.

The metric of d-prime was calculated as:

$$d' = z(hit rate) - z(false alarm rate)$$

$$hit rate = \frac{TP}{TP+FN}$$

$$false alarm rate = \frac{FP}{FP+TN}$$

where z(·) denotes the inverse cumulative distribution function of the standard normal distribution.

**3. Calibration curves and decision-curve analyses**

**3.1 Ten-year mortality risk predictions**

Mortality models were evaluated on the held-out UK Biobank test set. For each candidate predictor, a Cox proportional hazards model was fitted on the training set using time to death as the outcome, with chronological age and sex included as covariates. Individual-specific 10-year mortality risks on the test set were obtained as one minus the model-predicted survival probability at 10 years. The corresponding binary outcome was defined as death from any cause within 10 years of follow-up among participants with sufficient observation time.

**3.2 Calibration assessment**

Agreement between predicted and observed 10-year risks was summarized using calibration curves, scalar calibration metrics, and logistic recalibration parameters. Calibration curves were constructed with scikit-learn (sklearn.calibration.calibration_curve): predicted risks were grouped into 10 bins of equal sample size (quantile binning), and within each bin the mean predicted risk was plotted against the observed event proportion.

The Brier score was computed as the mean squared error between the binary 10-year outcome and the predicted risk (sklearn.metrics.brier_score_loss), with lower values indicating better probabilistic accuracy. Expected calibration error (ECE) was calculated by partitioning predicted risks into 10 equal-width intervals on the unit interval. For each non-empty bin, we took the absolute difference between the observed event rate and the mean predicted risk, multiplied this difference by the proportion of individuals in the bin, and summed across bins.

Calibration-in-the-large (intercept) and calibration slope were estimated by logistic recalibration in statsmodels. The binary 10-year outcome was regressed on an intercept and the logit of the predicted risk using a generalized linear model with binomial family (statsmodels.api.GLM with statsmodels.api.families.Binomial), after adding a constant term with statsmodels.api.add_constant. Predicted probabilities were bounded away from 0 and 1 before logit transformation. The fitted intercept and slope quantify systematic over- or under-estimation of risk and the degree of shrinkage or expansion of predicted probabilities, respectively; values of 0 and 1 indicate perfect calibration.

**3.3 Decision-curve analysis**

Clinical utility of the predicted 10-year mortality risks was assessed with decision-curve analysis using the net-benefit framework of Vickers et al. For each threshold probability pt, net benefit was computed as the proportion of true positives minus the proportion of false positives weighted by the odds at the threshold, pt / (1 − pt). Model net benefit was compared with treat-all and treat-none strategies across a grid of threshold probabilities from 0.01 to 0.30. Treat-all net benefit was calculated from the outcome prevalence, and treat-none net benefit was set to zero. Analyses were implemented directly in NumPy.
