## Supplementary Notes for "Decoding heterogeneous aging clocks and disease risk stratification using MetAgeFormer"

MetAgeFormer: supporting ablation, robustness and quality-control analyses

This document reports supporting analyses that were carried out during the development and evaluation of MetAgeFormer but are not required for the main conclusions of the manuscript. They are provided in full for completeness and reproducibility, and are referenced from the main text where relevant. All analyses use the same data, pre-processing and quality-control pipeline described in the Methods of the main manuscript. Unless stated otherwise, UK Biobank (UKB) analyses use the training set (n = 434,366) for model fitting and the held-out test set (n = 73,417) for evaluation; Cox models of subtype-level disease risk are adjusted for chronological age, sex and body mass index (BMI), and head-to-head comparisons of aging metrics are adjusted for chronological age and sex.

Contents:

Supplementary Note 1. Contribution of self-supervised pre-training

Supplementary Note 2. Comparison with non-linear Gompertz-based survival baselines

Supplementary Note 3. Robustness of the metabolic subtypes

Supplementary Note 4. Technical reproducibility and stability of subtype assignment

Supplementary Note 5. Cross-cohort reproducibility of subtype-defining metabolite effects

Supplementary Note 6. Missing data

Supplementary Note 7. Sensitivity of the head-to-head benchmarking to BMI adjustment

### **Supplementary Note 1. Contribution of self-supervised pre-training**

#### **1.1 Effect of pre-training on mortality prediction and on training stability**

To quantify the contribution of the self-supervised pre-training stage, we trained an otherwise identical model directly on the survival objective, without pre-training, and compared it with the pre-trained-then-fine-tuned MetAgeFormer on the UKB held-out test set.

Pre-training substantially improved mortality discrimination: the C-index was 0.765 (95% CI 0.759–0.771) with pre-training and 0.715 (95% CI 0.709–0.721) without pre-training (**Supplementary Note Figure 1A**). The training dynamics differed accordingly. With pre-training, validation loss decreased smoothly (≈0.485 to ≈0.477) and the validation C-index rose steadily (≈0.740 to ≈0.750) over 35 epochs (**Supplementary Note Figure 1B**), indicating stable fine-tuning from an already informative representation. Without pre-training, validation loss remained at ≈0.498 and the validation C-index at ≈0.694, with essentially no improvement; under the default early-stopping criterion (monitoring validation C-index, patience = 5) training would halt by epoch 6, and extending the patience to 20 (31 epochs logged) still failed to produce meaningful learning (**Supplementary Note Figure 1C**).

These dynamics indicate that training a deep network from scratch with a single supervised survival objective is prone to instability, because mortality labels alone provide a supervisory signal that is too weak to shape a useful metabolomic representation. Self-supervised pre-training instead exploits the far richer signal contained in the metabolite profiles themselves, yielding a stable initialization for downstream survival fine-tuning.

The same conclusion holds for subtype discovery. Applying the Leiden algorithm with the settings used in the main analysis (`n_neighbors` = 15, `resolution` = 1.0) to embeddings learned from scratch under the survival objective yielded 85 clusters with an extremely unbalanced size distribution: three macro-clusters contained essentially the entire cohort (approximately 24%, 34% and 42% of the UKB training set), while the remaining 82 clusters were very small (median cluster size = 3). Supervised training from scratch therefore does not recover the structured metabolomic subtypes obtained after self-supervised pre-training.

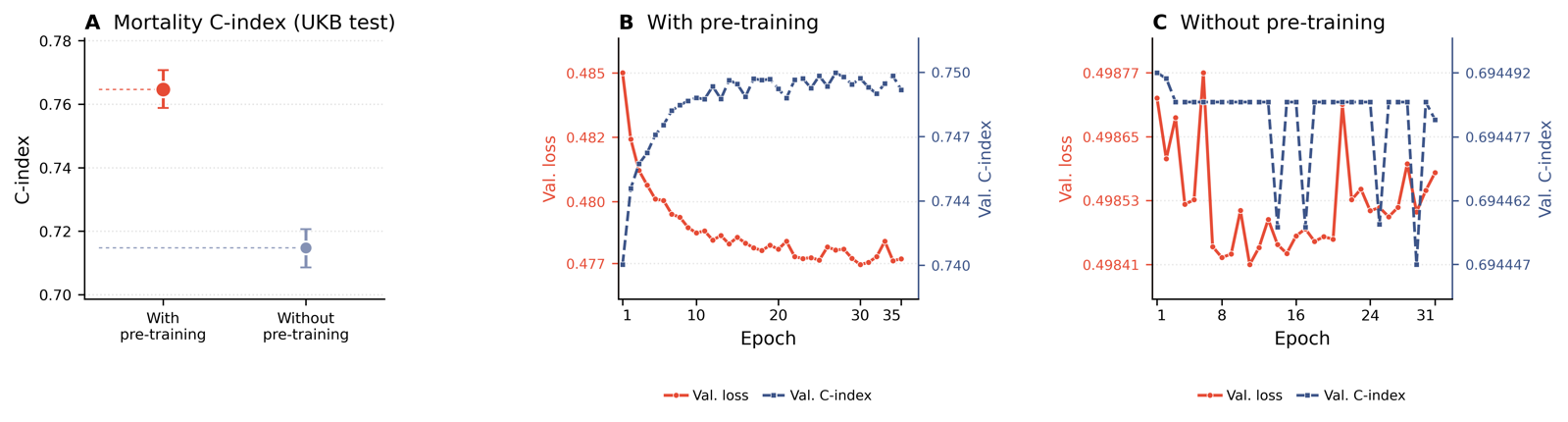

**Supplementary Note Figure 1.** Ablation of the self-supervised pre-training stage. (**A**) All-cause mortality C-index on the UKB test set with and without pre-training; error bars are bootstrap 95% confidence intervals. (**B**) Validation loss and validation C-index across fine-tuning epochs with pre-training. (**C**) The same metrics for a model trained from scratch on the survival objective.

#### **1.2 Subtype structure obtained without pre-training, from metabolite concentrations**

We next asked whether the subtype-level disease-risk structure reported in the main text depends specifically on the transformer representation, or whether it can also be recovered from the metabolite concentrations directly. We applied Leiden clustering with identical hyperparameters (`n_neighbors` = 15, `resolution` = 1.0) to the z-score-normalized matrix of the 107 non-derived NMR metabolite concentrations.

Because Leiden labels are not uniquely defined (both the number of clusters and the cluster identifiers differ between runs), we compared solutions at the level of disease risk rather than by label-overlap indices. For each clustering solution we (i) fitted Cox models for the 12 incident disease endpoints adjusted for chronological age, sex and BMI and assembled the subtype × disease log₂(HR) matrix, (ii) matched subtypes across solutions by Hungarian assignment (Supplementary Note 3.1), and (iii) computed Spearman's ρ between the matched risk vectors.

Clustering the raw concentrations produced 17 subtypes, compared with 13 from the pre-trained embeddings, and a broadly concordant disease-risk matrix (ρ = 0.79). A subtype with elevated risk of all-cause dementia and chronic liver disease was present under both representations (highest all-cause dementia log₂(HR) ≈ 0.95 for embedding-derived subtype 8; ≈ 1.20 for concentration-derived subtype 14; **Supplementary Note Figure 2**).

This concordance does not make the learned representation redundant. It shows that the risk structure is a property of the metabolome rather than an artefact of the model, which supports the biological validity of the embedding. The embedding remains necessary as a shared, transferable feature space across cohorts and tasks — in particular for distillation to a cohort without NMR measurements (CHARLS) and for coupling subtype discovery to the metabolomic aging clock. Concordance at the level of subtypes also does not imply that concentration-only models match embedding-based clocks: relative to a clock trained on the identical NMR panel (MileAge, main-text Figure 4A) and to an otherwise matched model trained from scratch (Supplementary Note 1.1), the pre-trained embedding gives stronger metabolomic age estimation.

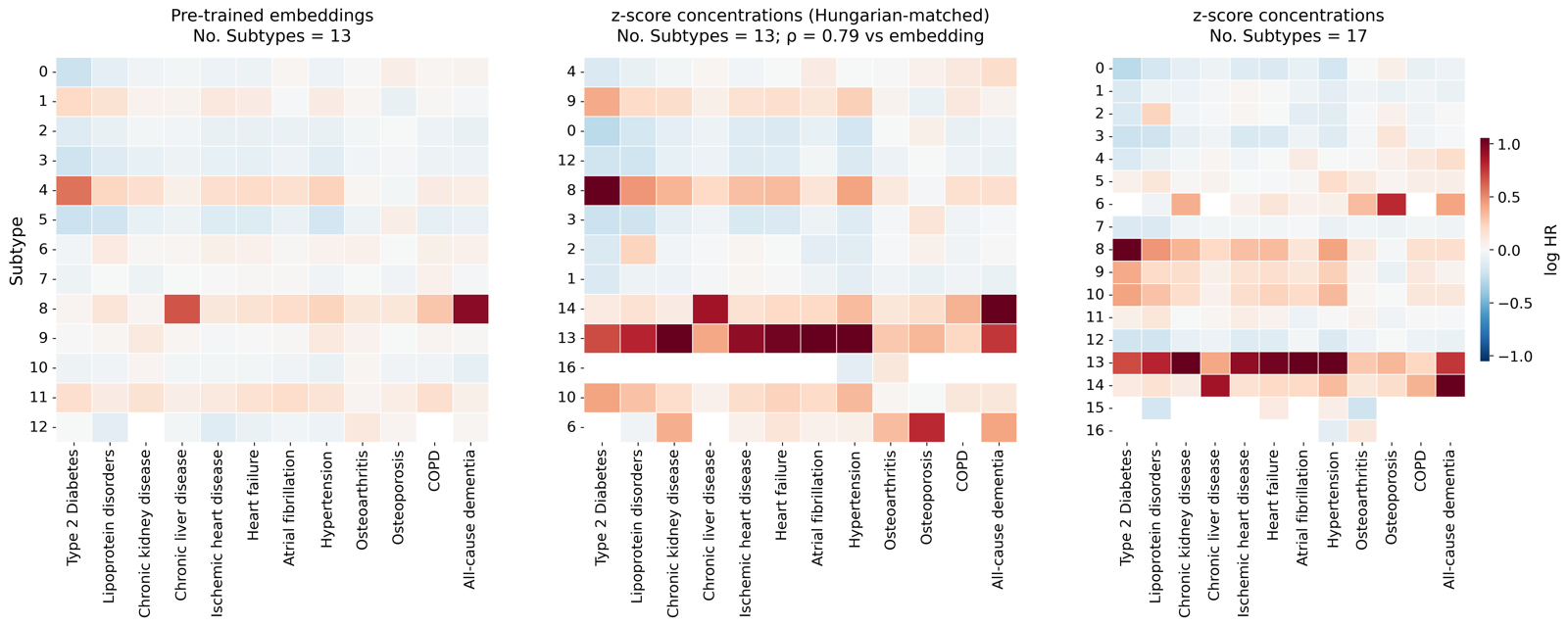

**Supplementary Note Figure 2.** Subtype-level disease risk derived from pre-trained embeddings versus from z-score-normalized metabolite concentrations. Left, log₂(HR) profile of the 13 embedding-derived subtypes; middle, the concentration-derived solution after Hungarian matching to the embedding solution; right, the full concentration-derived solution. Rows in the middle panel are ordered according to the matched embedding-derived subtypes.

### **Supplementary Note 2. Comparison with non-linear Gompertz-based survival baselines**

MetAgeFormer uses a standard transformer encoder; we do not claim a novel network architecture, and architectural novelty is not presented as a contribution of this work. To assess whether the transformer nevertheless provides benefit over strong non-linear tabular models under a shared survival objective, we implemented two additional Gompertz-based metabolomic clocks on the same 107 non-derived NMR metabolites and evaluated them on the same UKB test set (n = 73,417).

#### **2.1 Construction of the Gompertz-based baselines**

(i) *Gompertz baseline.* A parametric Gompertz model of right-censored mortality was fitted on the training set using chronological age (x_age) only, so that the hazard at follow-up time t is h(t) = exp(α_age + γ_age · x_age) × exp(γ_age · t). Maximum-likelihood estimation yields (α_age, γ_age). For each participant we then computed the cumulative hazard implied by chronological age alone over the observed follow-up time, H₀ = [exp(α_age + γ_age · x_age) / γ_age] × [exp(γ_age · t) − 1], that is, the mortality risk expected from age in the absence of metabolomic information.

(ii) *Excess-mortality target.* With event = 1 for an observed death during follow-up and 0 otherwise, the raw discrepancy between the observed outcome and the age expectation is (event − H₀). To obtain a numerically stable regression target we used y = clip[(event − H₀) / max(H₀, H_floor), −3, 3], where H_floor is the median H₀ among training-set participants who died during follow-up. Values y > 0 indicate higher mortality than expected for chronological age and y < 0 lower mortality than expected.

(iii) *Metabolomic models.* XGBoost and a multilayer perceptron (hidden layers of 256, 128 and 64 units) were trained by mean-squared error to predict y from the 107 non-derived metabolites; hyperparameters were selected by randomized search on a 10% training subsample and the models were then refitted on the full training set. The prediction ŷ is a metabolite-based estimate of excess mortality relative to the Gompertz age baseline.

(iv) *Mapping to metabolomic age.* Because γ_age converts a unit change on the linear predictor into years of chronological age, metabolomic age was defined as μ_age + ŷ / γ_age + c, where μ_age is the mean training-set chronological age and c = μ_age − mean(μ_age + ŷ / γ_age) is a single calibration constant, computed once on the training set and reused for all splits, that sets mean metabolomic age equal to mean chronological age.

#### **2.2 Results**

Metabolomic age derived from MetAgeFormer achieved a mortality C-index of 0.765 (95% CI 0.759–0.771), compared with 0.736 (0.730–0.742) for the XGBoost-Gompertz clock and 0.746 (0.740–0.752) for the MLP-Gompertz clock (**Supplementary Note Figure 3**). The absolute gains are approximately 0.029 and 0.019, respectively, and the MetAgeFormer confidence interval does not overlap that of either baseline. Pre-trained transformer representations therefore add predictive value beyond strong non-linear models applied to the same input panel under the same survival formulation.

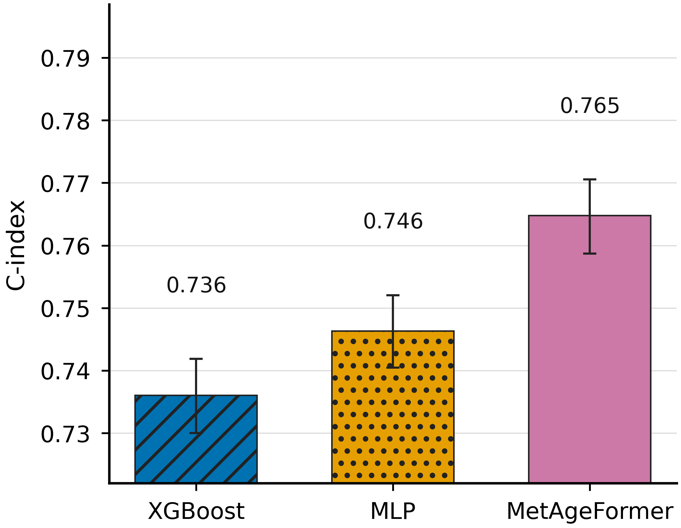

**Supplementary Note Figure 3.** Mortality prediction performance of Gompertz-based metabolomic aging clocks (XGBoost, MLP) and MetAgeFormer. Error bars are bootstrap 95% confidence intervals (1,000 resamples of the UKB test set).

### **Supplementary Note 3. Robustness of the metabolic subtypes**

The 13 metabolic subtypes reported in the main text were obtained by Leiden clustering of the pre-trained embeddings with `n_neighbors` = 15 and `resolution` = 1.0. Uniform manifold approximation and projection (UMAP) is used for visualization only; subtype labels are defined on the k-nearest-neighbour graph of the embeddings and do not depend on UMAP parameters. The analyses below examine whether the subtype-level risk structure is stable with respect to clustering parameters, embedding normalization, potential confounders and the label-transfer step used for external cohorts.

#### **3.1 Hungarian matching of clustering solutions**

Because clustering labels are arbitrary and the number of clusters differs between settings, solutions were compared through their disease-risk profiles using the following procedure.

Step 1. Given two log₂(HR) matrices A and B of subtype-specific disease risk (rows, subtypes; columns, diseases), each obtained from Cox models adjusted for chronological age, sex and BMI.

Step 2. Construct a cost matrix, cost[i, j] = 1 − PearsonCorr(A[i, :], B[j, :]).

Step 3. Apply Hungarian assignment (scipy.optimize.linear_sum_assignment) to obtain the one-to-one pairing with minimum total cost.

Step 4. Reorder matrix B according to the assignment and drop unmatched extra subtypes.

Step 5. Flatten the matched subtypes and compute Spearman's ρ between the two risk vectors.

#### **3.2 Sensitivity to Leiden clustering parameters**

We re-derived cluster labels under alternative settings (`nn15/res0.5`, `nn15/res2.0`, `nn10/res1.0`), which yielded 8, 29 and 18 subtypes, respectively. The number of subtypes therefore depends on the resolution and neighbourhood parameters, as expected. The disease-risk structure, however, was preserved: after Hungarian matching, all alternative settings recovered risk profiles that were highly concordant with the original solution (ρ = 0.90 for `nn15/res0.5`, 8 of 8 subtypes matched; ρ = 0.89 for `nn15/res2.0`, 13 of 29 matched; ρ = 0.85 for `nn10/res1.0`, 13 of 18 matched), and the high-risk subtypes were recovered under every setting (**Supplementary Note Figure 4**).

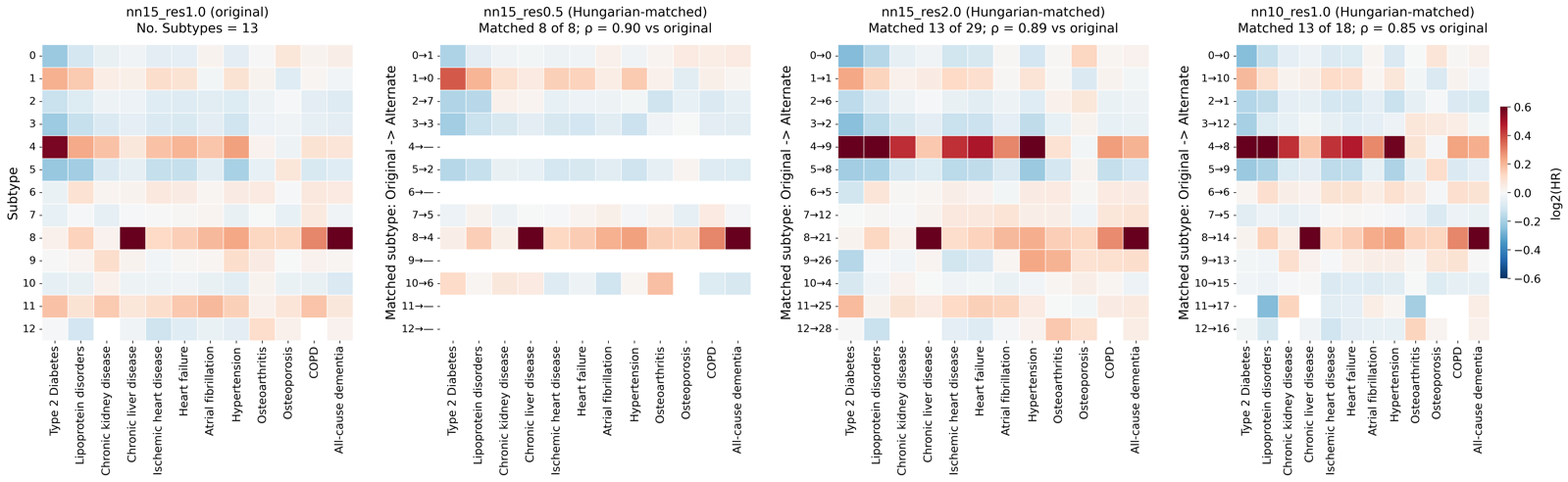

**Supplementary Note Figure 4.** Subtype-level disease-risk profiles under alternative Leiden clustering parameters, after Hungarian matching to the original solution (`n_neighbors` = 15, `resolution` = 1.0).

#### **3.3 Raw versus L2-normalized embeddings**

Leiden clustering of L2-normalized embeddings with the same parameters (`nn15/res1.0`) produced a different partition from the 13 subtypes obtained with raw embeddings, but a highly concordant disease-risk matrix (ρ = 0.82). A dementia-elevated subtype was preserved (subtype 8 with raw embeddings and subtype 10 with L2-normalized embeddings), and disease-onset patterns were consistent between the two representations (**Supplementary Note Figure 5**).

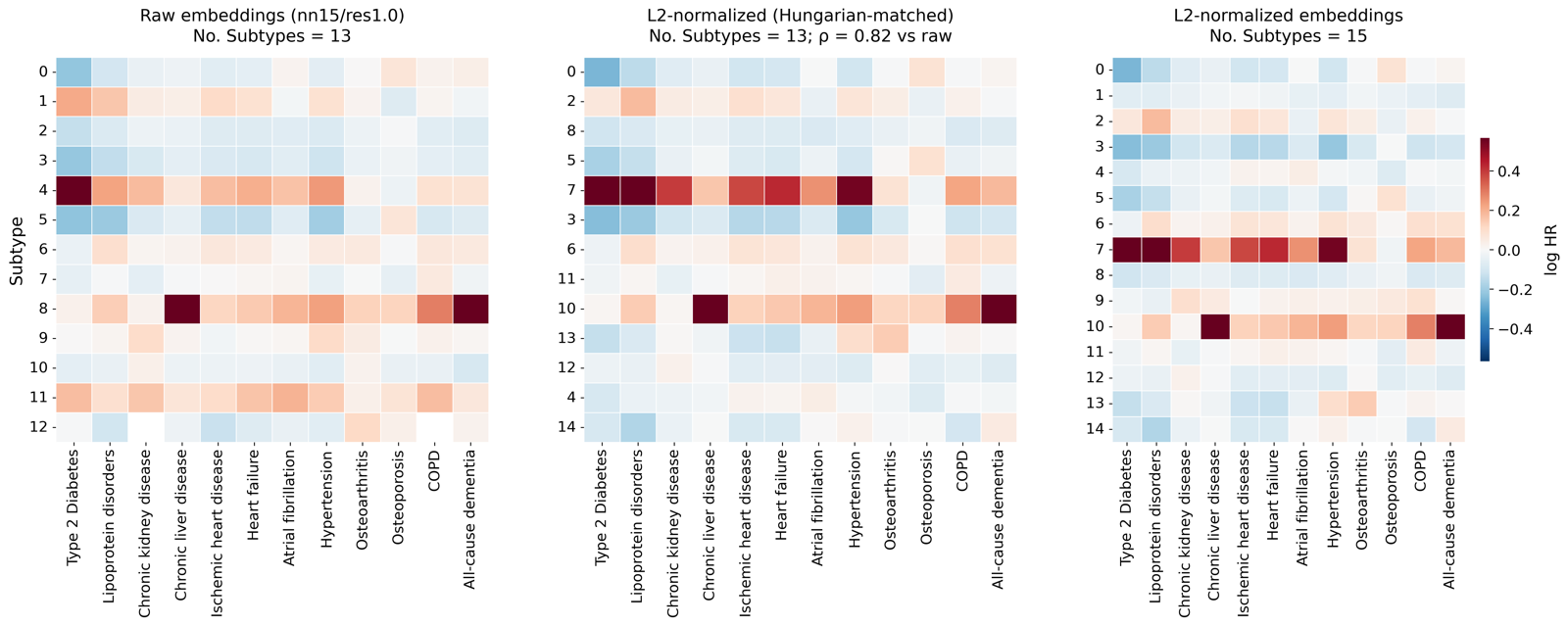

**Supplementary Note Figure 5.** Subtype-level disease risk derived from raw versus L2-normalized embeddings. Rows in the middle panel are ordered according to the Hungarian-matched subtypes from raw embeddings.

#### **3.4 Subtype membership after accounting for demographic, sampling and treatment covariates**

To examine whether subtype membership can be attributed to demographic, sampling or treatment-related covariates rather than to metabolic features, we trained a supervised classifier to recover the original Leiden labels from the embedding together with these covariates. The 512-dimensional pre-trained embedding was compressed by an MLP to a bottleneck representation (z ∈ R²⁷) whose dimensionality matches that of the covariate vector: four continuous variables (chronological age, BMI, fasting time [UKB Data-Field 74] and medication count [Data-Field 137]), two binary variables (sex and any medication use) and 21 assessment-centre indicators. Subtype labels were then predicted by a classification head trained end-to-end with cross-entropy loss.

On the UKB test set (n = 73,255), subtype membership was highly concordant between assignments made with and without the covariate block (**Supplementary Note Figure 6**), and subtype sizes were almost unchanged. Agreement was 93.1% at the subtype level and 95.6% at the meta-subtype level. We note that this analysis demonstrates that subtype labels are recoverable predominantly from the embedding, with the covariates adding little; it does not by itself establish statistical independence of the subtypes from these covariates. Consistent with this, all subtype-level, meta-subtype-level and ∆Age-trajectory analyses of disease risk in the main text are additionally adjusted for BMI, because BMI differs systematically across metabolic subtypes and is therefore a potential confounder of subtype–outcome associations.

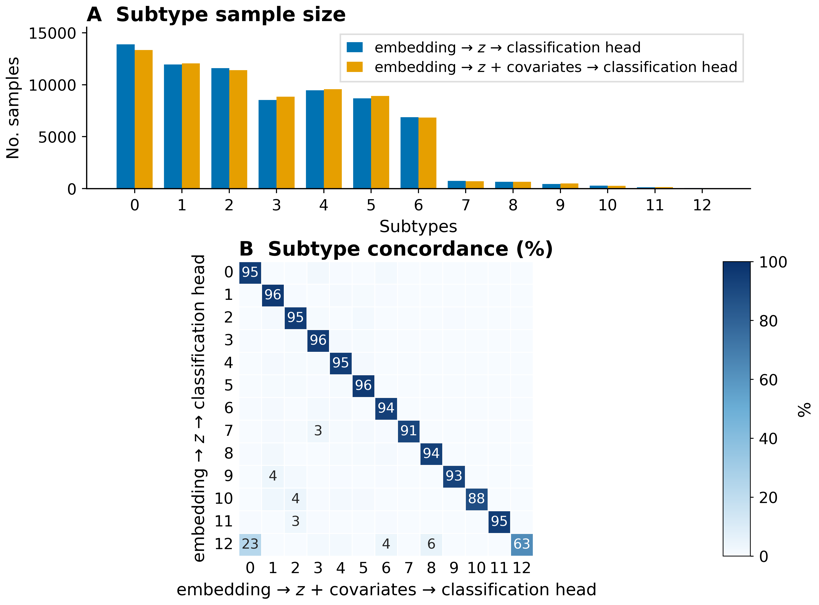

**Supplementary Note Figure 6.** Concordance of subtype membership assigned from the embedding alone versus from the embedding plus chronological age, sex, BMI, medication use, fasting time and assessment centre (UKB test set, n = 73,255).

#### **3.5 De novo clustering versus label transfer in CHARLS**

In the main text, CHARLS participants are assigned to subtypes using the MLP classifier fitted on the UKB training set. To verify that the external subtype structure does not rest solely on transferred labels, we additionally clustered the CHARLS 2011 embeddings de novo with the same Leiden parameters (`nn` = 15, `res` = 1.0) and repeated the Cox and Hungarian workflow of Supplementary Note 3.1, using 12 self-reported chronic conditions from the CHARLS health questionnaire (field `da007_*` of `health_status_and_functioning.dta`; see Methods).

Disease definitions differ between the two cohorts: CHARLS conditions are physician-diagnosed self-reports, whereas UKB endpoints are ICD-coded from hospital records. In CHARLS, memory-related disease, diabetes and liver disease are each ascertained from a single self-report item, and the memory-related item covers several disorders (for example dementia, brain atrophy and Parkinson's disease). We therefore assessed directional consistency rather than exact endpoint matching.

Matched log₂(HR) concordance between the MLP-transferred and de novo solutions was ρ = 0.63 at the subtype level (**Supplementary Note Figure 7**). The UKB-trained classifier assigned no CHARLS 2011 participant to subtype 10 and only two participants to subtype 12; both subtypes were excluded from the CHARLS log₂(HR) heat map. After merging the 13 subtypes into the four meta-subtypes defined in UKB — with each de novo cluster first matched to an MLP subtype by risk-profile concordance and then assigned that subtype's meta-subtype — concordance rose to ρ = 0.79 (**Supplementary Note Figure 8**). Independent clustering in CHARLS therefore recovers a comparable risk structure, and agreement is stronger at the meta-subtype level at which the external validation in the main text is performed.

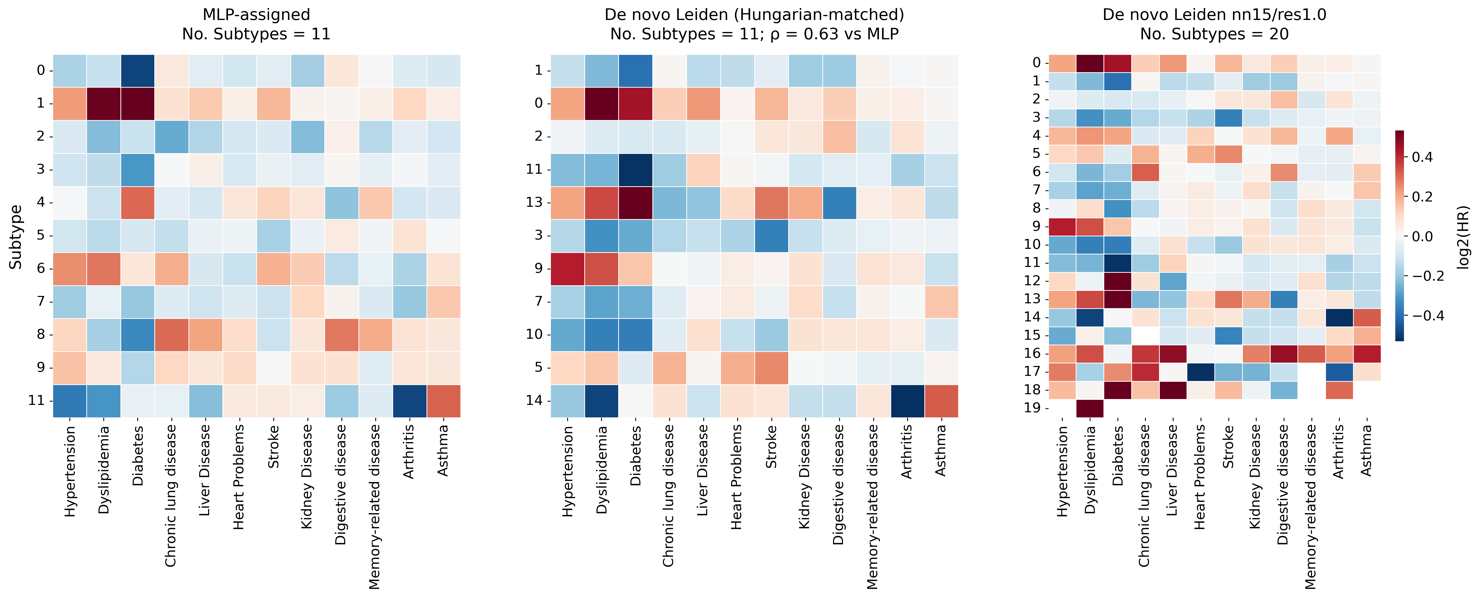

**Supplementary Note Figure 7.** Subtype-level disease risk in CHARLS 2011 under MLP label transfer versus de novo Leiden clustering. Rows in the middle panel are ordered according to the Hungarian-matched MLP-assigned subtypes.

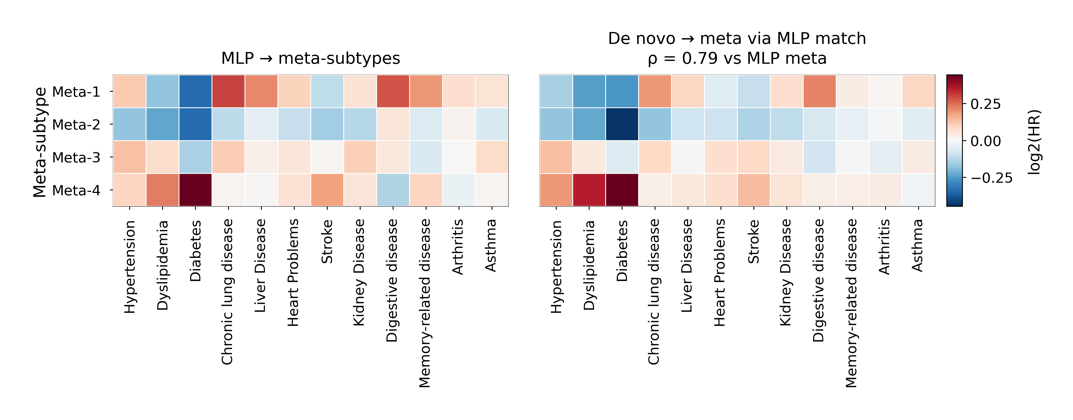

**Supplementary Note Figure 8.** Meta-subtype-level disease risk in CHARLS 2011 under MLP label transfer versus de novo Leiden clustering.

### **Supplementary Note 4. Technical reproducibility and stability of subtype assignment**

Longitudinal changes in discretized subtype membership should not be interpreted as metabolic-state transitions without evidence on assay precision, sampling-related covariates and label stability. We therefore assembled the following four lines of evidence (**Supplementary Note Figure 9**).

(i) *Technical reproducibility of the NMR platform.* All analyses use metabolite values after UK Biobank quality control and removal of technical variation, following the pipeline of Ritchie et al.¹ Platform-level metrics from Nightingale internal controls and UKB blind duplicates (Phase 3 companion documentation, UKB Category 220) show coefficients of variation (CV) below 5% for most biomarkers. Independently, Ritchie et al.¹ reported, across 3,169 UK Biobank blind duplicate pairs, a median CV of 4.55%, improving to 4.03% after removal of technical covariates (**Supplementary Note Figure 9A**). These figures address intra- and inter-batch CV and technical replicate reproducibility without requiring new assays.

(ii) *Metabolite intraclass correlation coefficients.* Among UKB participants with NMR measurements at instances 0 and 1 (n = 19,448 pairs), we computed the Shrout and Fleiss ICC(1,1)² for each of the 107 non-derived metabolites. The median visit-to-visit ICC was 0.61 (IQR 0.55–0.64) and 83% of metabolites had ICC ≥ 0.5. Low values were concentrated among short-half-life metabolites such as lactate, acetate and ketone bodies (**Supplementary Note Figure 9B**), as expected from biology rather than from global assay failure. Because the two visits are years apart, these coefficients combine genuine biological variation with residual technical noise and should be read as a lower bound on analytical reproducibility.

(iii) *Sampling-related covariates.* On the UKB test set with complete covariates (n = 73,255), subtype assignments from the embedding alone agreed with assignments from the embedding plus chronological age, sex, BMI, medication count, fasting time and assessment centre for 93.1% of individuals (95.6% at the meta-subtype level; **Supplementary Note Figure 9C**; see also Supplementary Note 3.4). Diet and inter-visit weight change are not available as repeated exposures; BMI and fasting time are included as the available proxies.

(iv) *Stability of subtype assignment.* Under alternative Leiden settings, subtype–disease log₂(HR) profiles remained highly concordant after Hungarian matching (ρ = 0.90, 0.89 and 0.85), and in CHARLS the MLP-transferred and de novo solutions agreed with ρ = 0.63 at the subtype level and ρ = 0.79 after merging to meta-subtypes (**Supplementary Note Figure 9D**). Among CHARLS participants with blood measurements in both 2011 and 2015 (n = 7,539), 34.2% retained the identical subtype and 54.5% the identical meta-subtype (**Supplementary Note Figure 9E**). Label turnover across years therefore partly reflects boundary effects in a continuous embedding space in addition to biological change, which is why the manuscript treats meta-subtypes as the appropriate granularity for longitudinal use and does not describe label changes as definitive metabolic-state transitions.

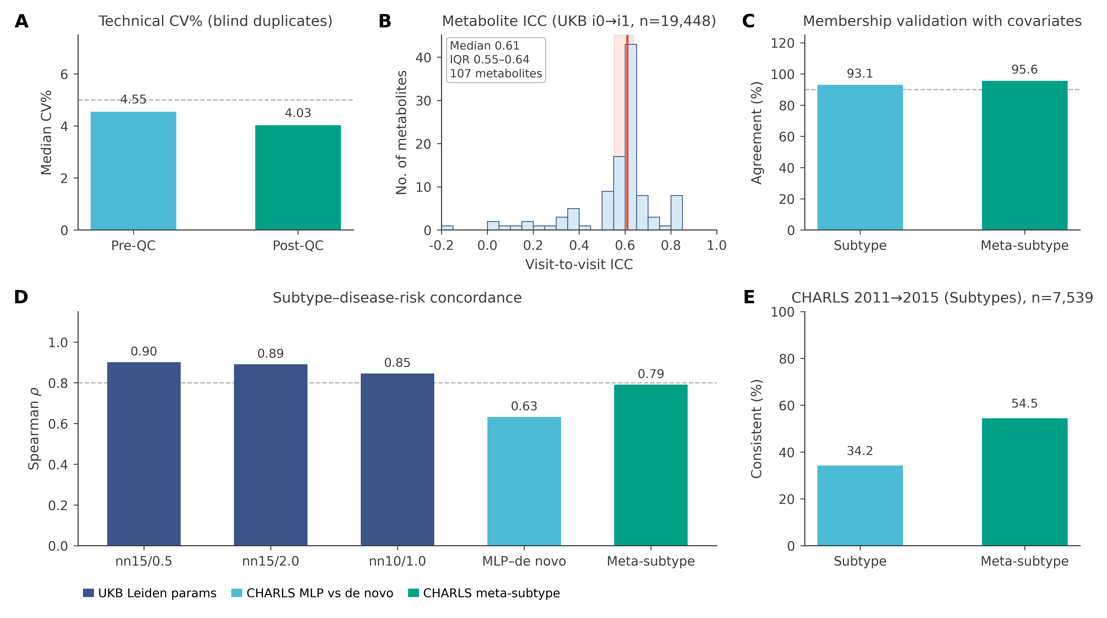

**Supplementary Note Figure 9.** Technical reproducibility and subtype-assignment stability. (**A**) Median CV of NMR metabolites in UKB blind duplicates before and after removal of technical variation (Ritchie et al.¹, n = 3,169 pairs); dashed line, 5%. (**B**) Visit-to-visit ICC(1,1) for the 107 non-derived metabolites in participants with NMR data at instances 0 and 1 (n = 19,448 pairs); orange line, median; shaded band, IQR. (**C**) Agreement of subtype and meta-subtype labels assigned from embeddings alone versus embeddings plus chronological age, sex, BMI, medication count, fasting time and assessment centre (UKB test set, n = 73,255). (**D**) Concordance of subtype–disease log₂(HR) profiles (Spearman ρ after Hungarian matching) under alternative UKB Leiden settings and for CHARLS MLP-transferred versus de novo clustering, at subtype and meta-subtype level. (**E**) Fraction of CHARLS participants (n = 7,539) retaining the identical subtype or meta-subtype between the 2011 and 2015 waves.

### **Supplementary Note 5. Cross-cohort reproducibility of subtype-defining metabolite effects**

To assess whether the metabolite effects that characterize particular subtypes in UK Biobank are reproduced in an independent cohort, we compared standardized effect sizes between UKB and ADNI. ADNI Nightingale NMR visits were assigned to UKB Leiden subtypes with the UKB-trained subtype classifier; for each highlighted metabolite we recomputed Cohen's *d* comparing the assigned subtype with all other subtypes within ADNI; and UKB and ADNI effect sizes were then plotted side by side for a curated set of subtype–metabolite pairs.

We did not screen all subtype–metabolite combinations. Instead we restricted the comparison to metabolites that are defining features of specific UKB subtypes: subtype 8, the acetate-elevated subtype, together with its related ketone, alanine and HDL-triglyceride signals; subtype 0, characterized by ketone-body (β-hydroxybutyrate) elevation; and subtype 1, characterized by elevated GlycA and alanine. This asks a focused question, namely whether these defining effects retain their direction and approximate magnitude in an independent cohort under transferred subtype labels.

In ADNI, subtype 8 remained strongly acetate-elevated (*d* = 4.71, versus *d* = 9.92 in UKB), with concordant direction for acetone, alanine and L_HDL_TG; the β-hydroxybutyrate elevation of subtype 0 and the GlycA and alanine elevation of subtype 1 also showed close agreement between cohorts (**Supplementary Note Figure 10**). These results indicate cross-cohort reproducibility of key subtype-associated metabolite effects. Consistent with the conservative wording used throughout the manuscript, they characterize the subtypes and do not establish any of these metabolites as a clinical biomarker.

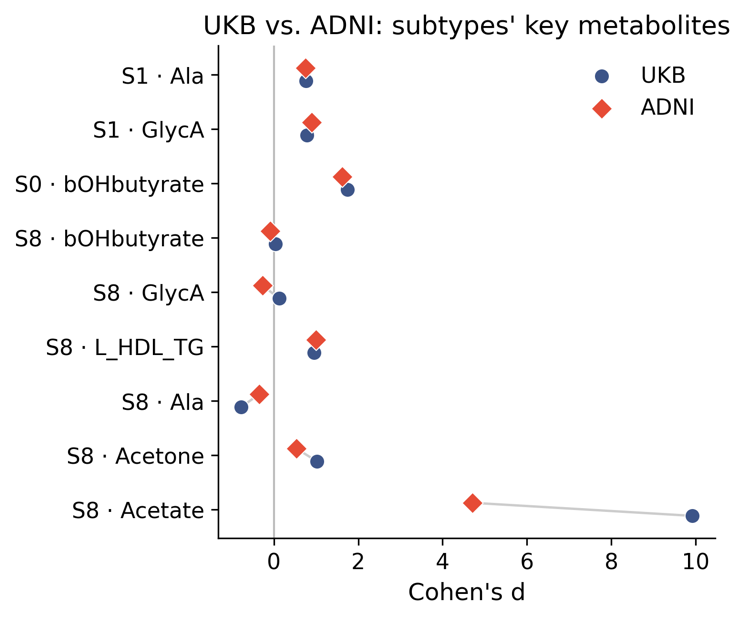

**Supplementary Note Figure 10.** Concordance between UK Biobank and ADNI of standardized effect sizes (Cohen's *d*) for subtype-defining metabolites, with ADNI subtypes assigned by the UKB-trained classifier.

### **Supplementary Note 6. Missing data**

#### **Missingness in the NMR panel and in routine blood biomarkers**

Missing or zero metabolite concentrations are not excluded from modelling. During pre-training, a subset of observed concentrations is replaced by a special mask token and the model is trained to reconstruct the held-out values from the remaining metabolite context. At inference, and wherever native missingness is present, missing or zero entries are represented by the same learnable mask embedding, so that incomplete profiles are retained in the input representation rather than removed by complete-case filtering.

Missingness in the NMR panel is low after quality control and removal of technical variation. The overall missing fraction across the 107 non-derived metabolites is 0.215% in UKB and 0.331% in ADNI. The highest per-metabolite rates occur among sparsely quantified lipoprotein subclasses and related analytes (UKB: `XXL_VLDL_PL` 3.78%, `XXL_VLDL_TG` 3.64%, β-hydroxybutyrate 2.08%; ADNI: β-hydroxybutyrate 1.36%; **Supplementary Note Figure 11**).

Missingness is more common among the routine blood biomarkers used by the lightweight distilled model. Values are z-scored using the mean and standard deviation of the UKB training set, with missingness preserved through scaling and encoded by a learnable mask embedding rather than imputed; the lightweight model additionally applies hidden-layer dropout (probability 0.1). In UKB (n = 507,783) the overall missing rate is 6%, the highest being glucose (13.6%) and HDL cholesterol (13.5%). In CHARLS the overall rates are 3.45% in 2011, driven largely by cystatin C (24.9%), and 0.71% in 2015.

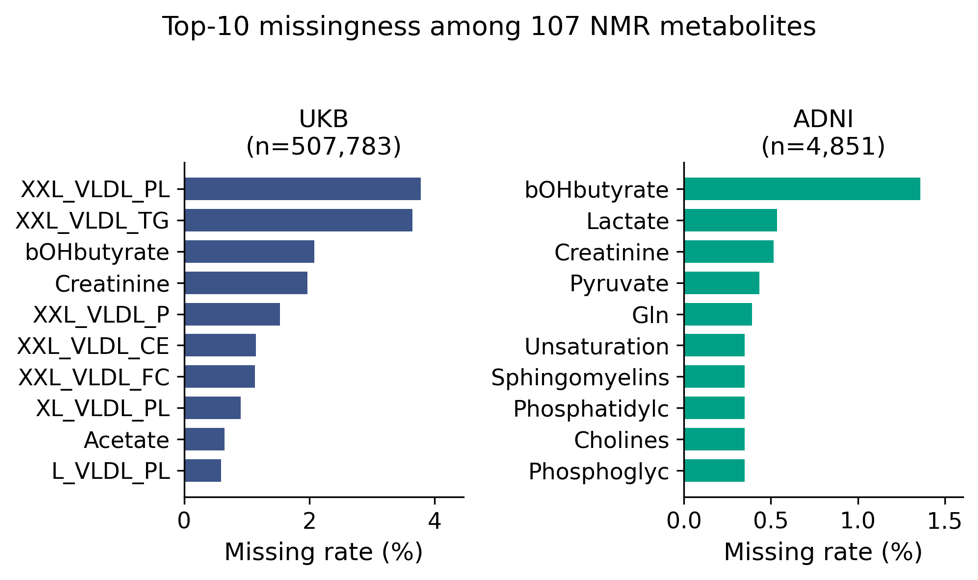

**Supplementary Note Figure 11.** The ten NMR metabolites with the highest missing rates in the UK Biobank (left) and ADNI (right) cohorts.

### **Supplementary Note 7. Sensitivity of the head-to-head benchmarking to BMI adjustment**

The covariate specification used in the main text differs by analysis type by design. Head-to-head comparisons of aging clocks and markers, both for all-cause mortality (**Figure 3**) and for the 12 incident disease endpoints (**Figure 4**), are adjusted for chronological age and sex only, following the evaluation protocol of Hamilton et al., whose published model weights and endpoint definitions we adopted, so that no measure is favoured by a covariate set different from the one used in its original report. All subtype-, meta-subtype- and ΔAge-trajectory-level analyses are additionally adjusted for body mass index (BMI), because subtypes differ systematically in BMI and BMI is therefore a plausible confounder of subtype–outcome associations. For a single continuous aging score the situation is different: BMI is itself a metabolic phenotype that is correlated with, and partly downstream of, the circulating metabolome, so adjusting a metabolomic aging clock for BMI removes part of the signal the clock is intended to capture (mediator over-adjustment) rather than controlling confounding.

To confirm that this choice does not drive any conclusion in the main text, we repeated the entire benchmarking analysis with z-scored BMI added as a further covariate, holding the analysis populations, endpoint definitions, bootstrap procedure and all other model terms fixed. Participants with missing BMI were dropped from the corresponding models, which reduced the effective sample size only marginally (at most 159 of 73,417 plasma samples in the largest comparison and 7 of 2,679 in the proteomics subset).

#### **7.1 All-cause mortality**

Adding BMI left mortality discrimination essentially unchanged. Across the 23 model–comparison rows the mean change in C-index was +0.001 (median +0.0003). MetAgeFormer ΔAge remained the best-performing measure in all seven comparison groups under both specifications, and its mean margin over the best competitor in each group was almost identical (0.029 without BMI versus 0.029 with BMI). With BMI in the model the MetAgeFormer C-index was 0.762–0.765 across comparison groups, against 0.721 for MileAge (ElasticNet), 0.747 for MetaboAge (Deelen), 0.747 for PhenoAge, 0.741 for Frailty Index, 0.731 for Cystatin C, 0.725 for HbA1c and 0.738 for ProteomicAge (Hamilton) (**Supplementary Note Figure 12**).

The hazard ratio per standard deviation of MetAgeFormer ΔAge attenuated from approximately 1.14 to approximately 1.08 (p < 0.001 under both specifications). This attenuation without loss of discrimination is the pattern expected when a covariate captures part of the same biological variation as the predictor rather than confounding it: the per-SD effect is shared with BMI, but the ranking of individuals by predicted risk, which is what the C-index measures, is preserved.

#### **7.2 Incident disease endpoints**

Across the 276 paired disease models the mean C-index rose by 0.016 when BMI was added. The increase was concentrated in BMI-proximal endpoints; for MetAgeFormer in the MileAge comparison, for example, the C-index rose from 0.807 to 0.837 for type 2 diabetes, from 0.691 to 0.709 for hypertension and from 0.720 to 0.730 for chronic liver disease, while it was unchanged for all-cause dementia (0.748 to 0.747) and COPD (0.761 to 0.761). Because BMI improved the fit of the competitor models by a similar or slightly larger amount, the differences between measures shifted only marginally.

In the Panel A comparison of ΔC-index (competitor minus MetAgeFormer), MetAgeFormer achieved the higher C-index in 68 of 84 measure–disease comparisons without BMI adjustment and in 66 of 84 with BMI adjustment; the mean shift of the plotted point was +0.004. Six comparisons changed sign, and all were small relative to their bootstrap confidence intervals: hypertension against ProteomicAge (Hamilton) (−0.010 to +0.002), lipoprotein disorders against MileAge (−0.002 to +0.001) and osteoarthritis against MetaboAge (Deelen) and MileAge (both approximately −0.002 to +0.0003) moved in favour of the competitor, whereas osteoarthritis against Cystatin C (+0.001 to −0.001) and osteoporosis against ProteomicAge (Hamilton) (+0.003 to −0.002) moved in favour of MetAgeFormer. Every sign change involved an absolute ΔC-index below 0.01. The large advantages reported in the main text were unaffected, including type 2 diabetes, chronic liver disease and COPD against the metabolomic clocks and against ProteomicAge (Hamilton) (**Supplementary Note Figure 13A**).

#### **7.3 Incremental predictive value**

We also repeated the incremental analysis of **Figure 4B** with BMI included in every model, so that the demographic baseline became chronological age, sex and BMI. Adding MetAgeFormer to this baseline still improved the C-index for every disease endpoint, with a mean gain of 0.030 compared with 0.047 when BMI was absent from the baseline. The smaller gain is expected, since BMI itself now supplies part of the metabolic information that ΔAge would otherwise contribute. Adding MetAgeFormer on top of the baseline plus a competitor improved discrimination in 77 of 84 comparisons (mean gain 0.020 with BMI versus 0.026 without). For the two focus comparators shown in the main text, the full model containing both measures gave the highest C-index for 12 of 12 diseases against Frailty Index and for 11 of 12 against ProteomicAge (Hamilton), the single exception being osteoarthritis (ΔC-index = −0.001) (**Supplementary Note Figure 13B**).

Taken together, BMI adjustment changes the absolute discrimination of BMI-proximal endpoints, as expected, but does not alter the ranking of MetAgeFormer against any competitor for mortality, does not alter any of the substantial disease-level advantages, and does not alter the conclusion that metabolomic age acceleration carries prognostic information that is complementary to frailty and proteomic aging signals. The head-to-head results reported in the main text are therefore not an artefact of omitting BMI, and are retained in the age- and sex-adjusted form for comparability with the published clocks.

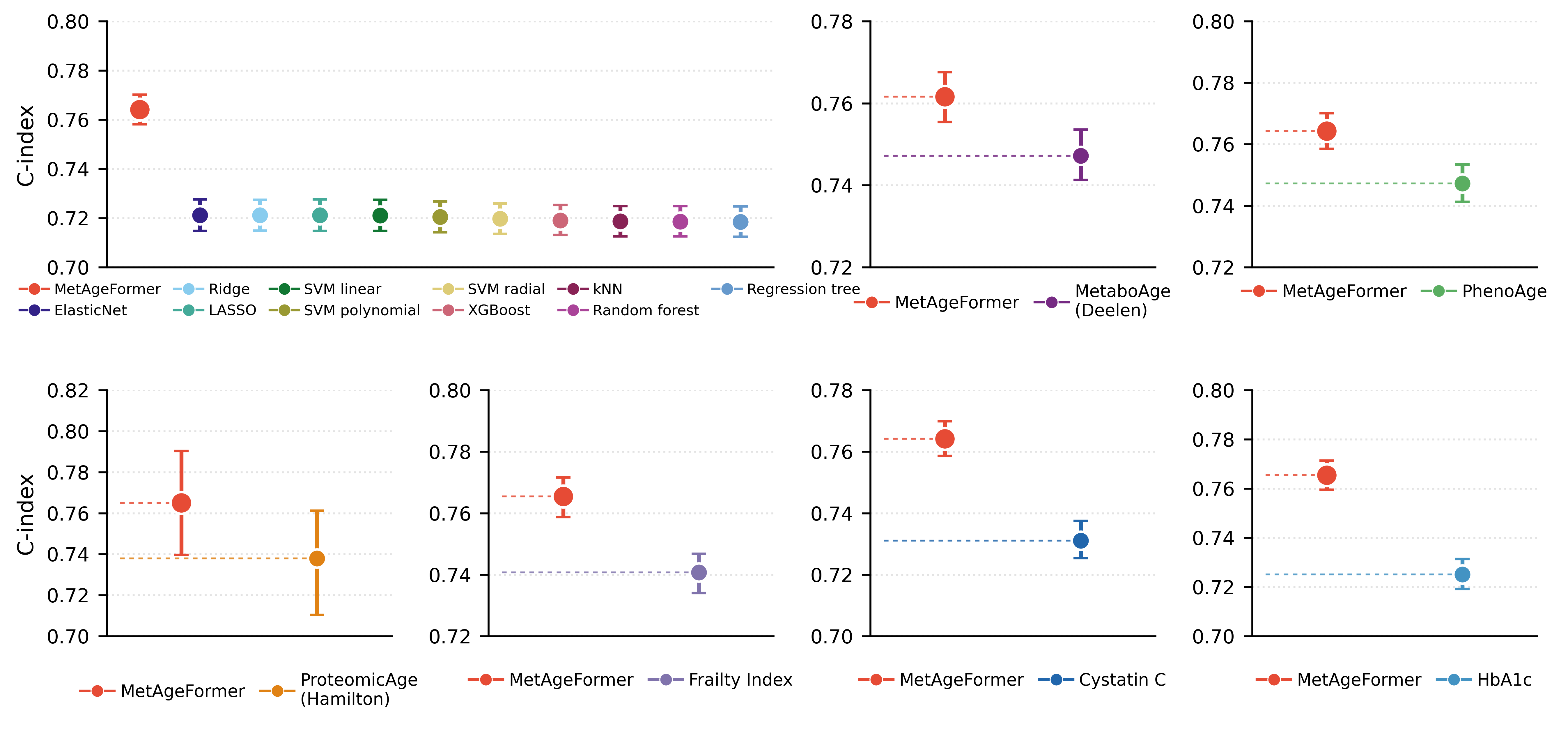

**Supplementary Note Figure 12.** Performance of aging clocks and markers in predicting all-cause mortality with additional BMI adjustment. Cox models are specified as z-scored ΔAge (or the aging marker, for non-clock comparators) plus chronological age, sex and z-scored BMI. C-index values are shown with 95% bootstrap confidence intervals. Panels correspond to those of Figure 3 of the main text, in which the same models are fitted without BMI.

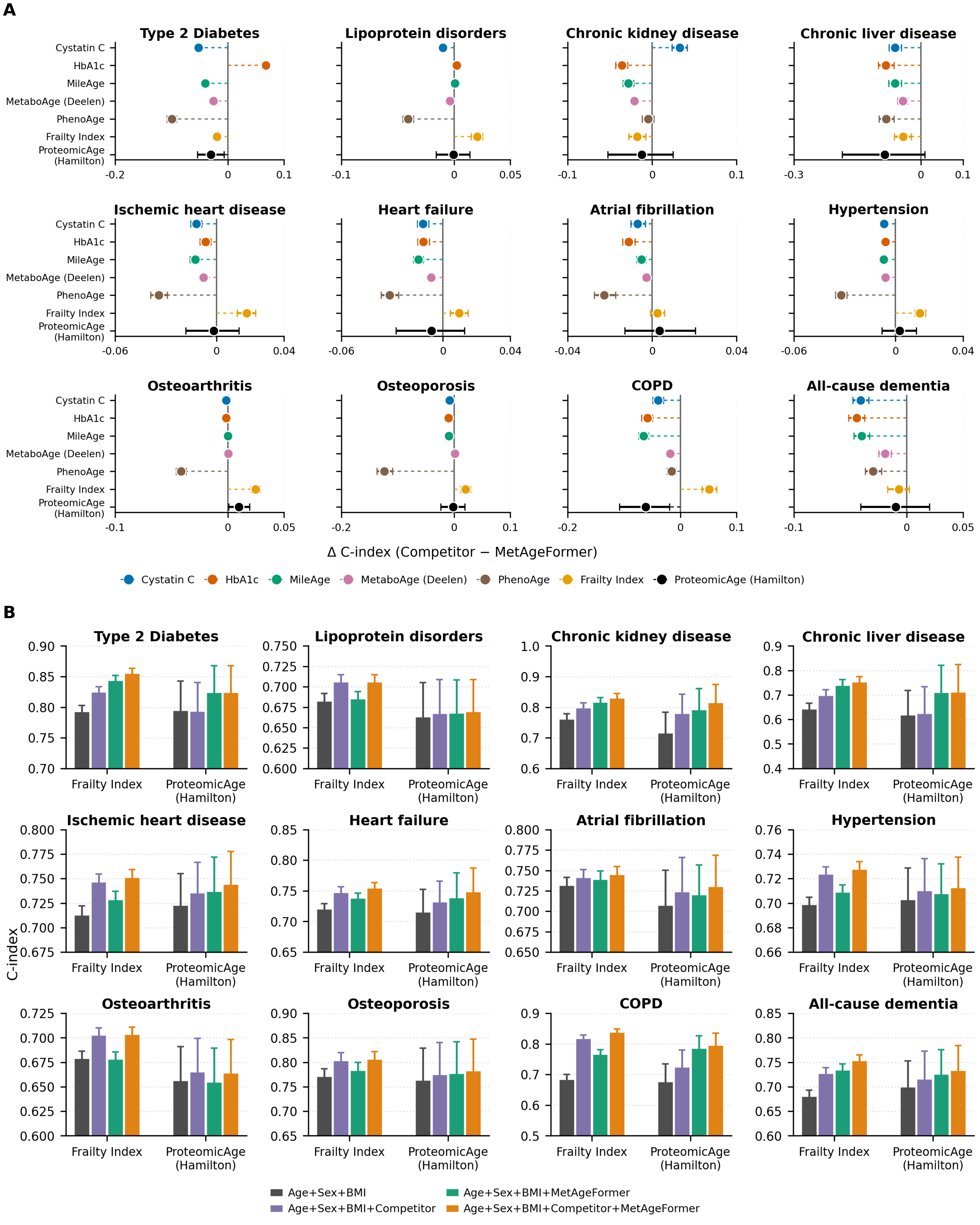

**Supplementary Note Figure 13.** Performance of aging clocks in predicting disease onset with additional BMI adjustment. (A) Difference in concordance index (ΔC-index; competitor minus MetAgeFormer) across the 12 age-related diseases, with all models additionally adjusted for z-scored BMI. Negative values indicate a higher C-index for MetAgeFormer. For MileAge, the highest-performing variant for each disease is shown. Points show estimates with 95% bootstrap confidence intervals (1,000 iterations). (B) Incremental predictive value of MetAgeFormer when added to models that already include chronological age, sex, BMI and either Frailty Index or ProteomicAge (Hamilton). Bars show mean C-index with standard deviations from 1,000 bootstrap iterations. Panels correspond to those of Figure 4 of the main text, in which the same models are fitted without BMI.
